# Identifying therapeutic drug targets for rare and common forms of short stature

**DOI:** 10.1101/2020.04.02.022624

**Authors:** Karol Estrada, Steven Froelich, Arthur Wuster, Christopher R. Bauer, Teague Sterling, Wyatt T. Clark, Yuanbin Ru, Marena Trinidad, Hong Phuc Nguyen, Amanda R. Luu, Daniel J. Wendt, Gouri Yogalingam, Guoying Karen Yu, Jonathan H. LeBowitz, Lon R. Cardon

## Abstract

While GWAS of common diseases has delivered thousands of novel genetic findings, prioritizing genes for translation to therapeutics has been challenging. Here, we propose an approach to resolve that issue by identifying genes that have both gain of function (GoF) and loss of function (LoF) mutations associated with opposing effects on phenotype (Bidirectional Effect Selected Targets, BEST). Bidirectionality is a desirable feature of the best targets because it implies both a causal role on the phenotype in one direction and that modulating the target activity might be safe and therapeutically beneficial in the other.

We used height, a highly heritable trait and a model of complex diseases, to test our approach. Using 34,231 individuals with exome sequence data and height, we identified five genes (*IGF1R, NPPC, NPR2, FGFR3*, and *SHOX*) with evidence for bidirectional effects on stature. Rare protein-altering variants significantly increased risk for idiopathic short stature (ISS) (OR=2.75, p= 3.99×10^−8^). These genes are key members of the only two pathways successfully targeted for short stature: Growth Hormone/Insulin-like growth factor 1 axis and C-type Natriuretic peptide (CNP) for Achondroplasia, a monogenic form of dwarfism. We assayed a subset of *NPR2* mutations and identified those with elevated (GoF) and diminished (LoF) activity and found that a polygenic score for height modulates the penetrance of pathogenic variants. We also demonstrated that adding exogenous CNP (encoded by *NPPC*) rescues the *NPR2* haploinsufficiency molecular phenotype in a CRISPR-engineered cell line, thus validating its potential therapeutic treatment for inherited forms of short stature. Finally, we found that these BEST targets increase the probability of success in clinical trials above and beyond targets with other genetic evidence. Our results show the value of looking for bidirectional effects to identify and validate drug targets.

## Introduction

Drug discovery has recently focused on target-based approaches to treat human disease instead of phenotypic screens. There are several ways in which potential targets are identified, the most common of which is the exploration of targets that have shown a mechanistic effect in either cell or animal models. The main disadvantage of these strategies is the high risk of failure when drug efficacy is tested in humans. Human genetic evidence for a target can improve the odds of success. A target with GWAS evidence has a two-fold higher probability of success compared to those targets with no evidence, and this is increased to 8-fold if the target is a gene that causes a Mendelian disorder.^1^ GWAS studies have begun to produce hundreds of hits for complex phenotypes,^2^ leading to challenges in prioritizing the large number of potential targets for those most likely to be amenable to therapeutic intervention. There are different approaches to prioritizing these genetic targets.^3–5^ The use of an allelic series of genetic variants that associate with the outcome of interest and mimic a dose-response curve can provide a direct indication that a promising target has been selected.^5^ It can also help to quantify the magnitude of the required modulation needed to achieve efficacy, and even inform us of potential adverse events.^5^

Some recent drug targets with strong human genetic evidence show an additional characteristic: they have allelic series that span a bidirectional effect. Gain-of-function (GoF) mutations are experiments of nature that might mimic the effect of a drug that increases the activity of a particular gene. Conversely, Loss-of-function (LoF, predicted LoF i.e., stop-gained, splice site disrupting, frameshift) mutations mimic a drug decreasing the activity of a gene product. The presence of mutations driving a phenotype into both supra- and sub-physiological levels supports the idea that a drug mimicking that effect may have similar clinical outcome and a tolerable safety profile. This also suggests that an entire disease population could be treated, even those patients with a different genetic etiology such as individuals with a high polygenic risk score for a complex disease. Therefore, the allelic series should ideally span both extremes to make a stronger case for druggability. For example, *PCSK9* GoF variants associate with higher LDL levels and higher cardiovascular disease (CVD) risk.^6^ Conversely, *PCSK9* LoF have the exact opposite effect (Fig 1. lower LDL levels, lower CVD risk).^7^ This observation led to the therapeutic hypothesis that anti-*PCSK9* neutralizing antibodies would lower LDL levels in all patients regardless of their *PCSK9* carrier status and also decrease risk for CVD. This strategy was eventually validated in successful Phase III randomized clinical trials.^8,9^ Similarly, GoF and LoF variants with bidirectional effects on two members of the WNT pathway (*LRP5* and *SOST*)^10^ led to the creation of a new therapy for osteoporosis using antibodies against *SOST*.^11^ This property of bidirectionality illustrated by the examples above should be generalizable to other complex phenotypes. We decided to test this hypothesis by using height as an example of a complex trait.

**Figure 1.**
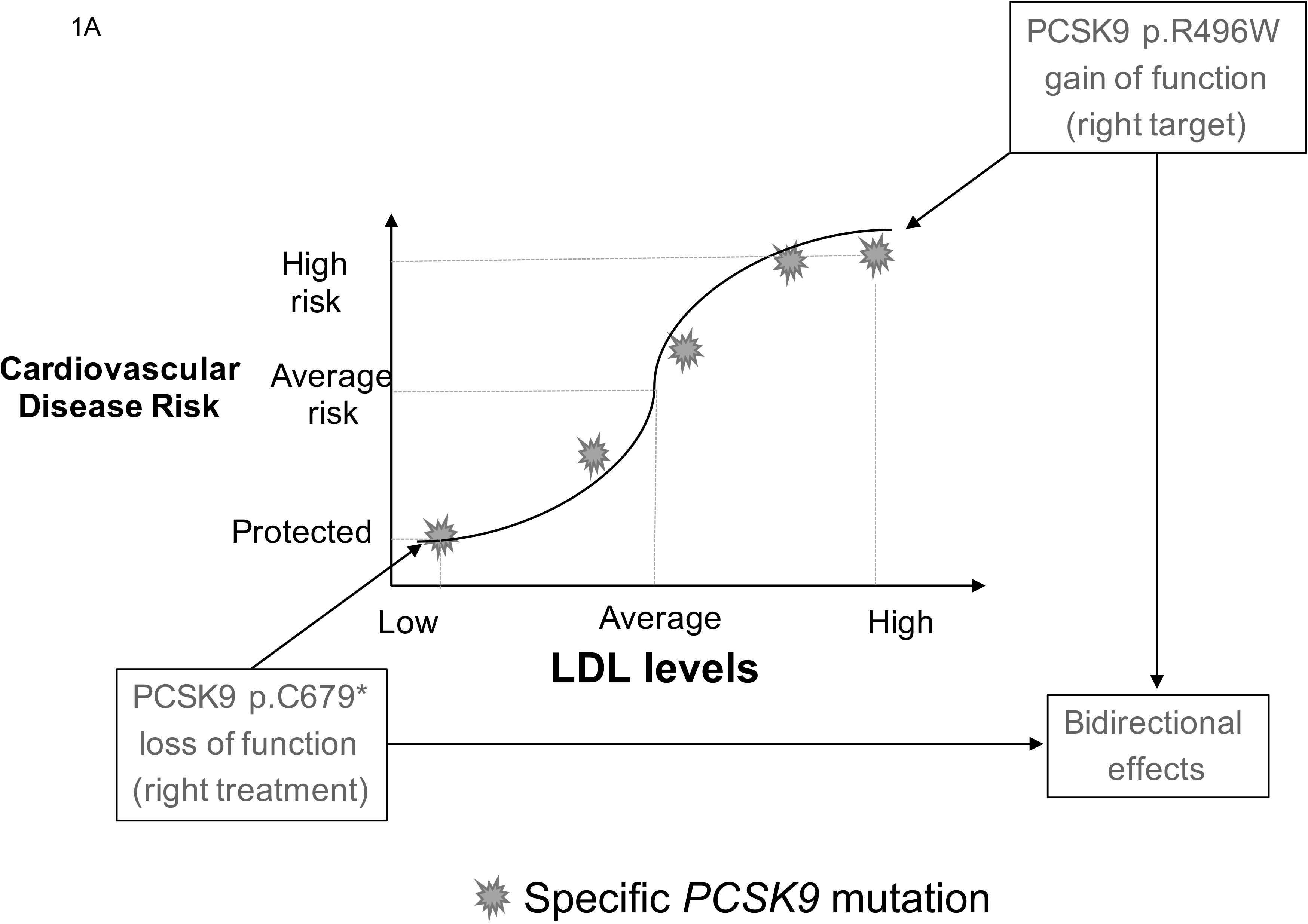
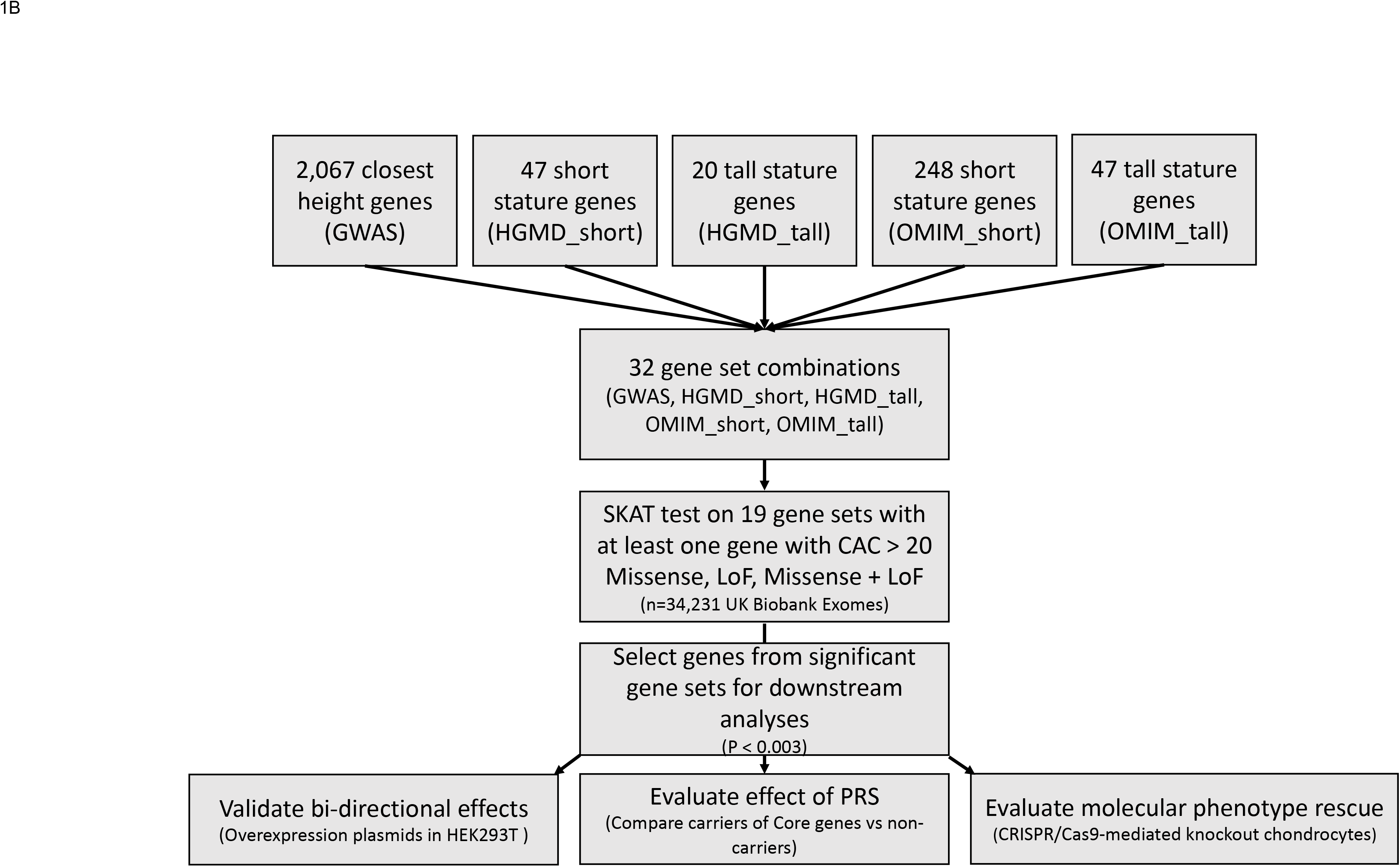
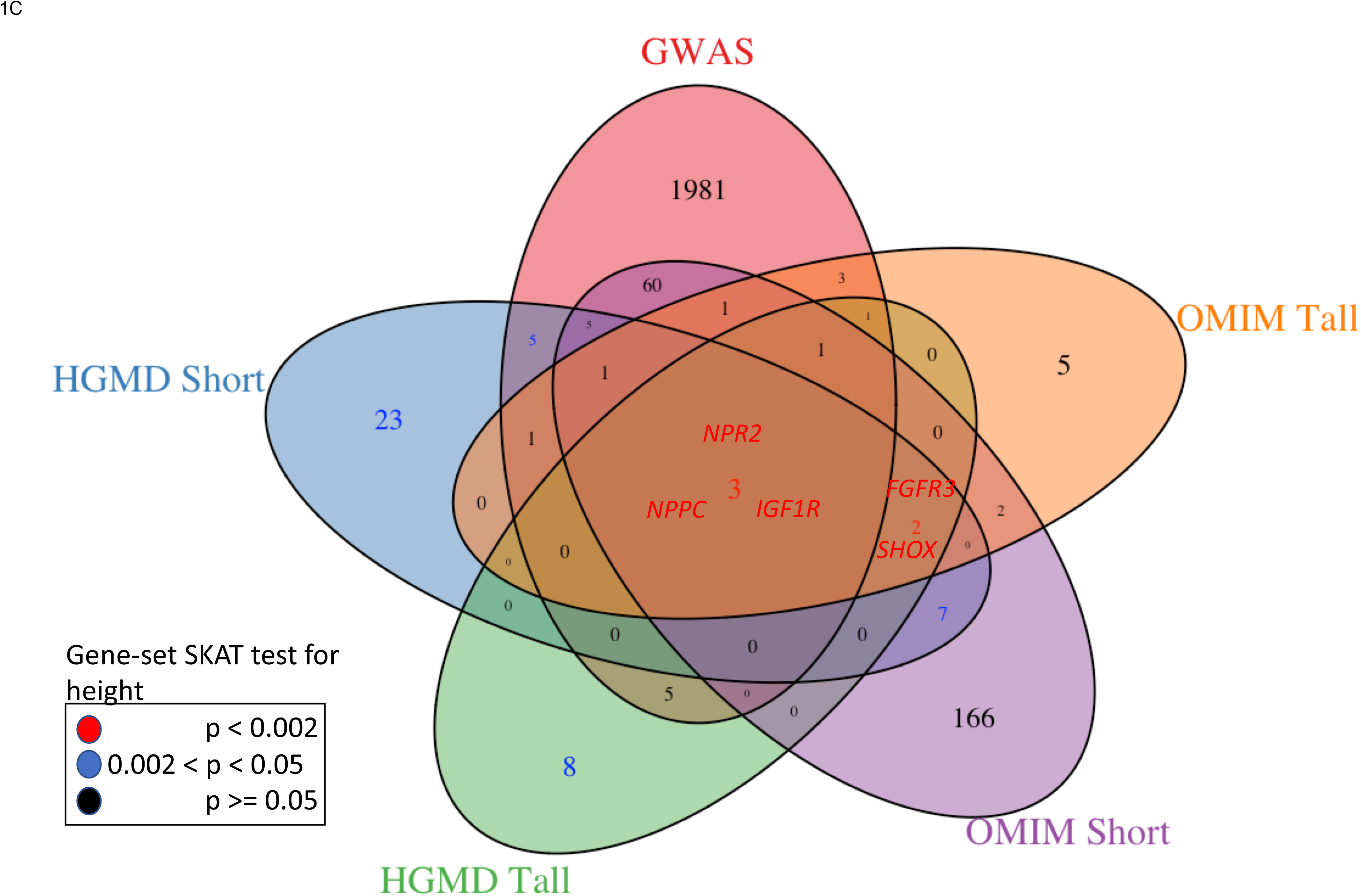
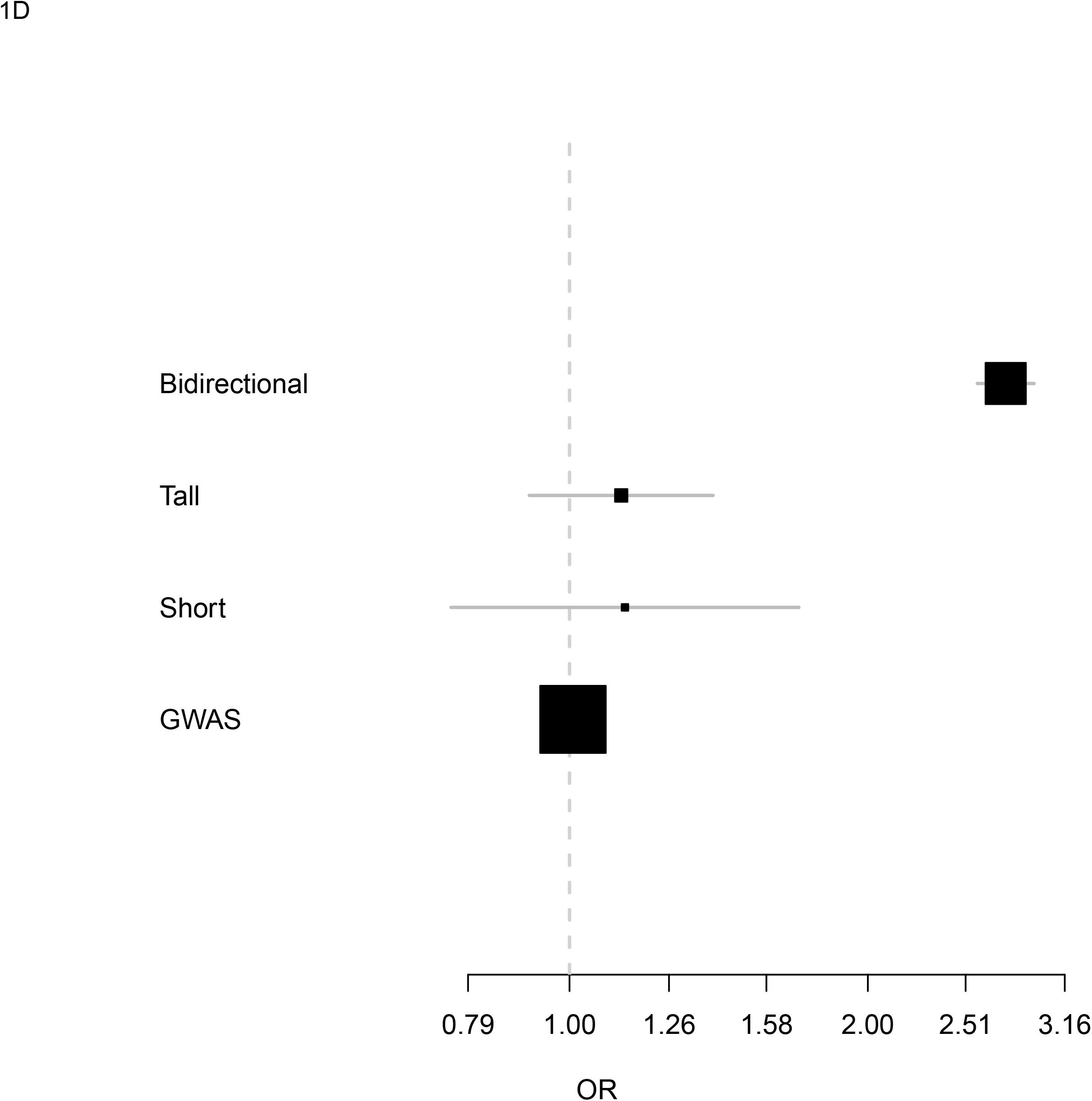
Strategy followed to identify bidirectional genes. A. Graphical representation of a gene (PCSK9) showing bidirectional effects on LDL and Cardiovascular disease risk. B. Flow chart describing the steps followed to identify causal genes. PRS, Polygenic Risk Score computed using previous meta-analysis of common variants. GWAS, Genome-Wide Association Study; HGMD, Human Gene Mutation Database; OMIM, Online Mendelian Inheritance in Man; SKAT SNP-set (Sequence) Kernel Association Test. C. Groups of genes analyzed to identify five genes that regulate height using both common and rare variation. The color codes of the gene names or the number of genes in the gene set corresponds to the SKAT test p-value for the association with height on 34,284 UK Biobank exome sequences. C. Odds Ratio for Idiopathic Short Stature (ISS) on highlighted gene-sets. Box represent estimate and 95% CI.

Height is a highly heritable (80%) and easily measured quantitative trait that has historically been used as a model to understand the genetic architecture of complex traits and diseases.^12^ At the low end of the height distribution, Idiopathic Short Stature (ISS, defined as having a height at least two standard deviations (SDs) below the mean)^12^ affects ~ 2% of the general population. Mutations in several genes have been described in studies of familial ISS as well as in cohorts of ISS patients.^13^

Mendelian forms of extreme short stature (height at least 5 SDs below the mean) such as Achondroplasia, Acromesomelic dysplasia type Maroteaux (ADMD) and Osteogenesis Imperfecta type III, are less frequent (1/15,000, 1/1 million, unknown respectively).^14–16^ There is no approved treatment for these extreme forms of short stature, and while growth hormone has been used for idiopathic short stature, response to treatment is variable.^17^ Thousands of loci have been associated with height using GWAS.^18–23^ Linking those association signals to genes, and those genes to mechanisms that may ultimately yield a new drug target has been challenging despite large scale efforts to map regulatory regions.^24,25^

Here, we aimed to prioritize height-associated genes to identify a subset of genes most likely to be good therapeutic targets by selecting those genes exhibiting bidirectional effects on height (Bidirectional Effect Selected Targets, BEST). The attributes of these BEST targets included: having significant GWAS signals, known bidirectional effects on rare protein coding variants with an extreme phenotype, and population-level evidence of association with the trait using rare variant association tests.

To test this approach, we used data for adult height from published and unpublished sources and tested its relevance in 33,204 adults with exome sequence data from the UK Biobank project. We functionally validated genetic variants and compared the predicted effect of these coding variants with their functional impact at a cellular level as well as with a clinical readout. We then conducted experiments to validate the use of a therapeutic approach in a cell model to rescue the molecular phenotype.

Finally, to determine whether the bidirectionality approach was truly generalizable, we curated a large database of known mutations to identify 98 genes with other bidirectional effects on other human diseases. We were then able to show that drugs targeting these genes are more likely to be approved in clinical trials. This trend was consistent across multiple stages of regulatory approval and a wide range of disease indications.

## Results

### Growth genes

First, we obtained summary statistics from the largest published meta-analysis for height based on imputed GWAS data from ~700,000 individuals.^18^ In total, the 2067 closest genes to genetic signals (within 1-MB window) were obtained (Supplementary Tables 1 and 2, Methods). Other approaches to map GWAS signals to genes such as co-localization using eQTL data were not suitable since they require gene expression data in the tissue most relevant to the phenotype under study. For height, this would be limb bone growth plates and there are currently no gene expression datasets available for this tissue.^26,27^

**Table 1.**
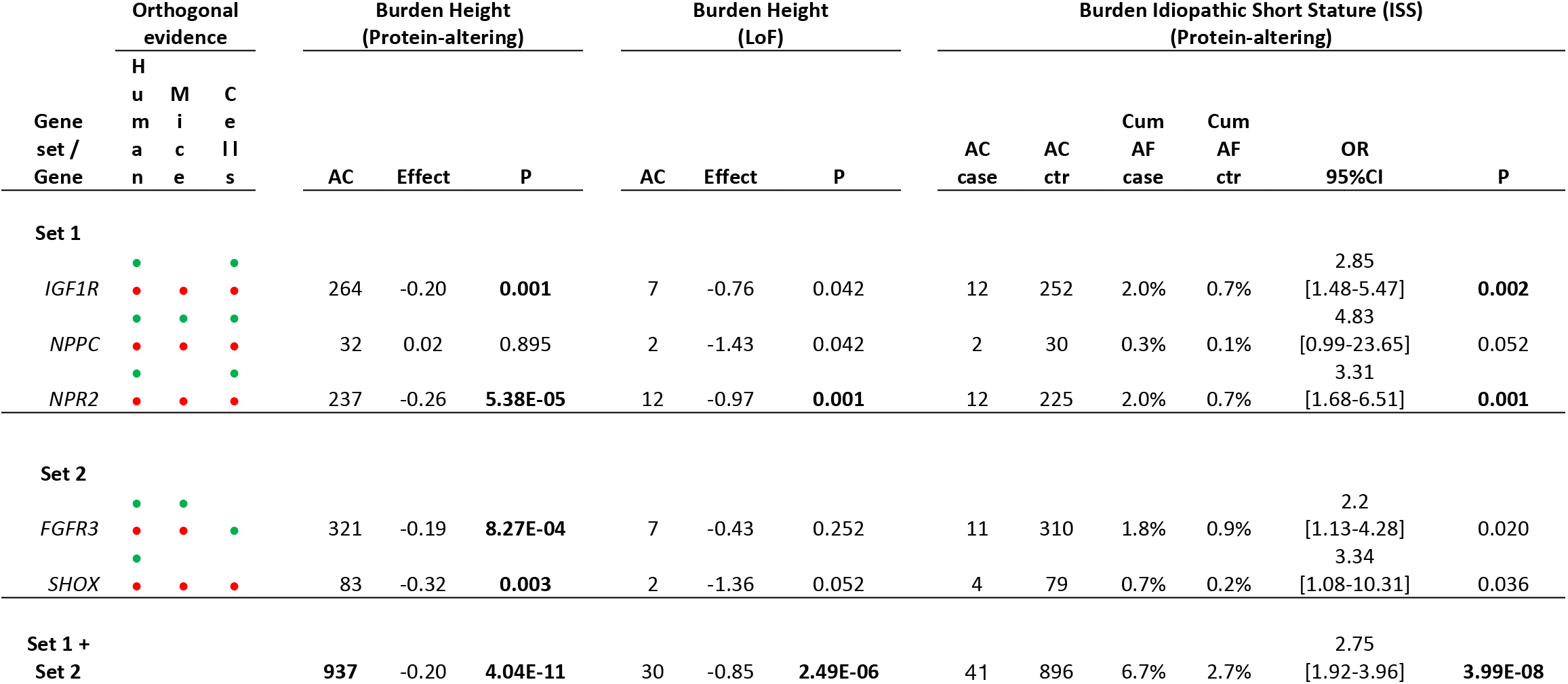
Individual gene association statistics for Height and Idiopathic Short Status (ISS, height < −2 SD below the mean) on rare protein-altering or LoF variants. Bold p-values represent significant associations after adjusting for five tests. AC, Allele count; LoF, Loss of Function; Effect, Effect estimate from regression; P, Nominal P-value estimate; Green and Red dots on the orthogonal evidence column reflects the presence of gain-of-function and Loss of function mutations in the respective model with corresponding evidence of effect on growth (For more details, see Supplementary Table 7)

To identify genes that are regulators of growth, we analyzed the intersections of five gene lists including the list of genes from the GWAS. Growth regulators would be the most likely to contain rare coding mutations with bidirectional effects (i.e. short stature or skeletal dysplasia AND tall stature or overgrowth). First, we queried the Human Gene Mutation Database (HGMD version v2019_2) for genes associated with short or tall stature.^28^ There were 47 genes annotated with at least one pathogenic variant reported in the literature to cause “short stature”. Only 20 genes were annotated as tall stature or overgrowth genes (Fig. 1 and Supplementary Table 2). Secondly, we used a manually curated list of 258 OMIM genes (248 short, 20 tall) which was created using the keywords: short stature, overgrowth, skeletal dysplasia, brachydactyly.^20^ Third, we looked at the intersection of these four lists with the list of genes from GWAS (Fig. 1 and Supplementary Table 2). We found at the intersection of these five lists contained three genes known to be associated with height (Set 1: *IGF1R*, *NPPC*, *NPR2*).

Two additional genes were present in the intersection of all lists except GWAS (Set 2: *FGFR3*, *SHOX*). *FGFR3* was not in Set 1 as it was not the closest gene in the region of any GWAS signal and *SHOX* was not tested in the GWAS study as it is located on the X-chromosome (Supplementary Figs. 1-4). Thus, at least for height, at this point there was no added value on using the intersection with the list of genes coming from GWAS. The *NPPC*/*NPR2* and the *IGF1R*/growth hormone pathways have been implicated in GWAS of height.^21,22^

Literature on extreme phenotypes such as short or tall stature is susceptible to publication bias. A randomly ascertained study population is required to obtain an unbiased assessment of the effects of rare coding variation of these genes on height on the general population. For that purpose, we used exome sequence data from 33,204 white British individuals from the UK Biobank project of which 572 (1.67%) reached the criterion of ISS (Supplementary Table 3. See Methods). After quality control, we identified 245,083 rare (allele frequency (AF) < 0.01%) missense and LoF variants in 1,964 genes with at least one rare protein coding variant. 1,918 genes with a mean cumulative allele count (AC) >= 20 remained after removing those with a cumulative AC < 20 (See methods).

### Gene set analysis

To evaluate if any of the annotations (GWAS signal adjacent, short stature, tall stature, both short and tall, *etc*.) had a significant association with height, gene set analysis was performed using the combined effect of any rare protein modifying variants using the SNP-set (Sequence) Kernel Association Test (SKAT).^29^ There were 32 possible intersections of the 5 annotations described above and 25 of those contained at least one gene (Fig. 1 and Supplementary Table 4). Of these, surprisingly, only the two aforementioned gene sets (Set 1 and Set 2) showed significant association with height. They both contained bidirectional effects as reported by both HGMD and OMIM, namely: Set 1: *NPR2*, *NPPC*, *IGF1R* (p=6.82×10^−7^) and Set 2: *FGFR3 and SHOX (p=1.75×10^−5^)* (Table 1, Fig. 1 and Supplementary Table 4). Given that both sets of genes share the property of having bidirectional effects on height, we combined all 607 independent rare protein-altering variants for the five genes in Set 1 or Set 2 using burden tests. The new group of five genes significantly decreased height (β=−0.20, p=4.04×10^−11^) and significantly increased risk for Idiopathic Short Stature (ISS) (OR=2.75, 95%CI [1.92 - 3.96], p = 3.99×10^−8^). Interestingly, this set of bidirectional genes has a much stronger risk for ISS as compared to the sets of genes reported for solely short stature or tall stature (Fig. 1C). Restricting the test to rare LoF variants resulted in an association with a larger decrease in height (β=−0.85, p=2.49×10^−6^). However, there were insufficient LoF carriers to calculate a precise odds ratio for ISS. Of the 607 variants, 158 were singletons and 192 (31.63%) were not reported in any of the public human databases and thus may represent novel pathogenic variants (Supplementary Table 5).

### Associated genes

While most of these mutations are too rare to be significantly associated individually, each of the five associated genes (*FGFR3, IGF1R, NPPC, NPR2* and *SHOX*) were still nominally (P <0.05) associated with height when considered individually, reflecting that no single gene was driving the gene set signal (Table 1, Online Methods). The direction of effects was consistent with LoF variants having Bonferroni-adjusted association with shorter stature (β=−0.97, P=0.001, for *NPR2*) and nominal associations in the same direction for *IGF1R* and *NPPC*. Rare missense variants were associated with shorter height (β=−0.22, P=0.002; β=−0.18, P= 0.003; and β=−0.18, P=0.001) for *NPR2*, *IGF1R* and *FGFR3* respectively. Combined LoF and missense variants in *NPR2* and *IGF1R* were also associated with increased risk for ISS (OR=3.31, P= 0.001, OR= 2.85, P=0.002, respectively) (Table 1).

Entire gene deletions and/or mutations causing loss of protein function in *SHOX*, *IGF1R*, *NPPC*, *NPR2* have been reported in familial short stature with various degrees of severity (Table 1, Supplementary Table 6).^*15,30–33*^ Conversely, duplications, deletions of repressor regions, translocations and missense mutations leading to GoF in these genes were reported in individuals with tall stature or overgrowth.^*34–38*^ GoF mutations are hard to identify from in-silico predictions and usually require functional validation. GoF mutations in *FGFR3* cause Achondroplasia, the most common form of dwarfism.^*39*^ For both *NPPC* and *IGF1R*, only large translocations and duplications have been reported with the overgrowth phenotype. In contrast, at least four independent *NPR2* missense mutations have been reported to be GoF.^32,36,37,40^ Importantly, mutations in these genes in animal and cell models also show bidirectional effects that are consistent with the directions of effects observed in humans. (Supplementary Table 7).

### Polygenic Risk Scores

The polygenic nature of complex traits suggests that the combined effect of thousands of variants may be an independent risk factor to rare variants with strong effects possibly modifying the overall risk. To test this hypothesis, we calculated polygenic risk scores (PRS) for height using the largest published GWAS meta-analysis for height that did not include any samples from the UK Biobank project (see Methods).^20^ We divided our cohort into five equally-sized (n=6,824) PRS quintiles (PRS 1 being the lowest height, PRS 5 the tallest height). As expected, there was a dose-dependent relationship between increasing PRS score and mean height (β=0.30 per each PRS quintile increase) (Fig. 2A). Carriers of LoF variants in the five genes were consistently shorter than non-carriers across the five different PRS backgrounds (Fig. 2A and Supplementary Figs 5-9. Our data suggest that the combined effect of PRS and rare protein variants is consistent with an additive model: polygenic effects modulated height in both carriers and non-carriers. The mean effect of having a rare protein coding variant in any of the five genes was similar across PRS subgroups (I^2^ = 30%, P=0.21; I2=0, P_het_=0.62 for missense and LoF variants respectively) (Fig. 2A, Supplementary Figs. 10-16). We then calculated the risk for ISS across PRS groups using individuals at PRS3 who were non-carriers of mutations in the five genes as a reference. Non-carriers of the lowest PRS group were associated with increased risk for ISS and non-carriers of the highest PRS group with a decreased risk (Fig. 2B; OR=5.81, P=2.74×10^−32^; OR=0.25, P=6.09×10^−6^ for PRS 1 and PRS 5 respectively). We then evaluated the effect of rare coding variants of the five genes for ISS stratified by PRS group. Carriers of any of the five genes were at increased risk for ISS in the first three quintiles (OR=15.39, P=4.44×10^−25^; OR=4.76, P=1.34×10^−4^; OR=5.96, P=1.21×10^−6^; OR=2.16, P=0.19 Fig. 2B). There were not enough ISS individuals in the highest PRS group (14 ISS for PRS 5, all non-carriers).

**Figure 2.**
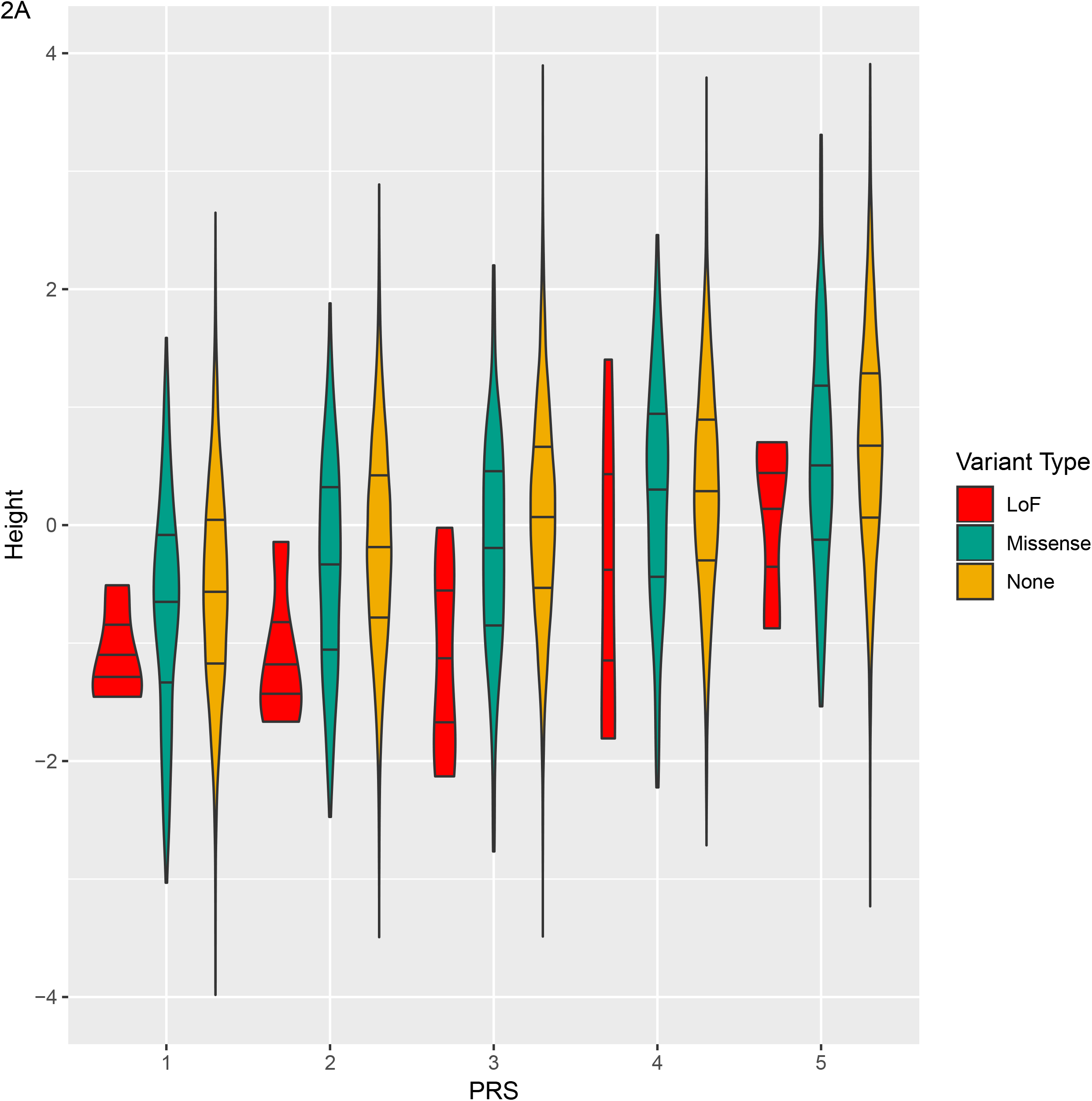
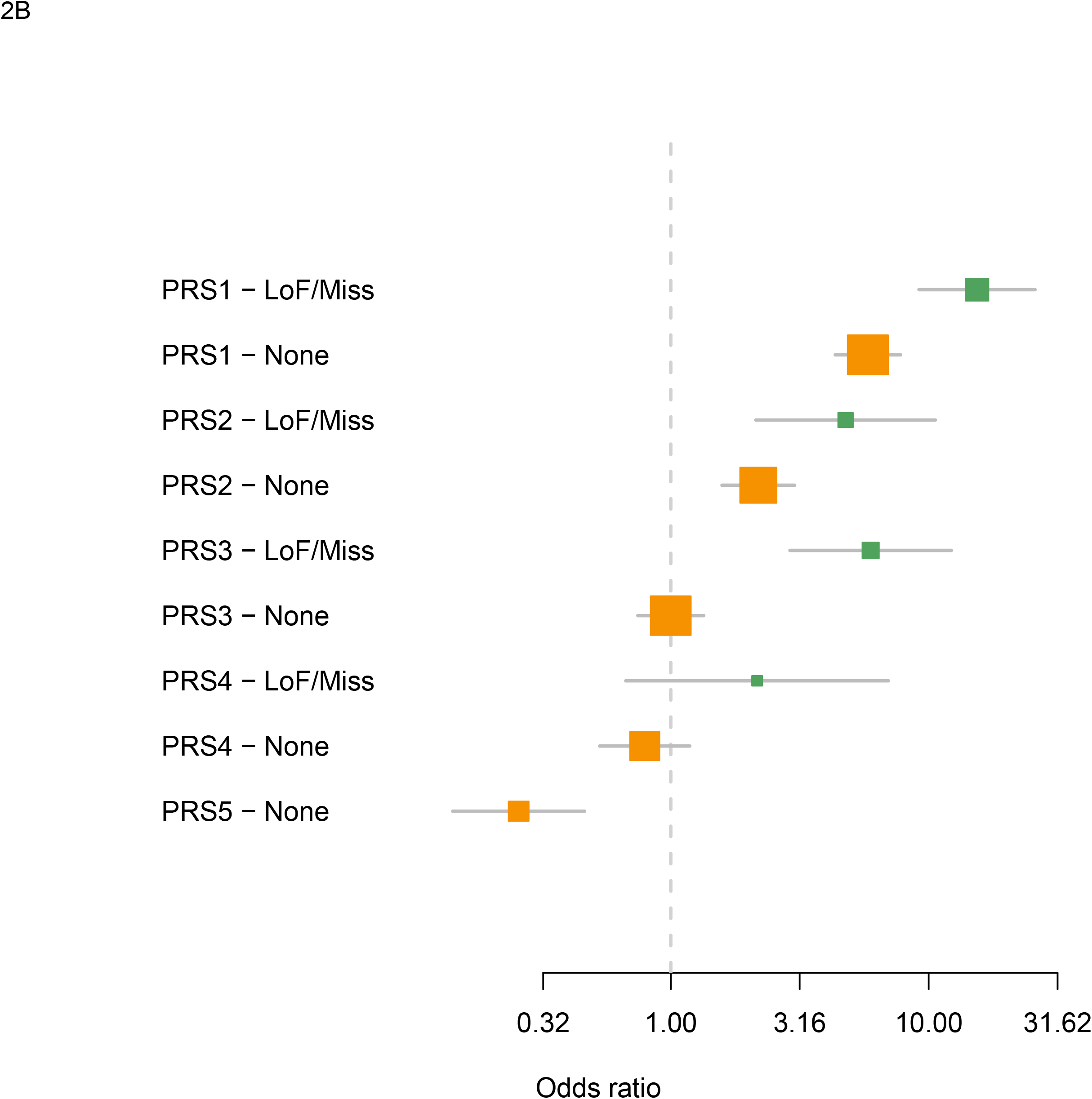
Combined effect of PRS and rare coding variants on height. A. Effects on height as a quantitative trait. Samples were divided in five groups based on their PRS. Horizonal lines in violin plots represent the 25%, 50% and 75% percentile of height. Samples were grouped by carrying status of missense (green), loss of function (red) or None (orange) in any of the five core genes. B. Effect reflected on Odds ratios for “Idiopathic Short Stature” or ISS. Odds for ISS using Non-Carriers of Core 5 genes with PRS =3 as reference vs the other PRS groups. Box colors represent: orange, non-carriers of LoF/missense variants in the five genes; green, carriers of LoF/missense variants in the five genes.

### Functional assessment of protein coding variants

A C-type natriuretic peptide analog (CNP, encoded by the *NPPC* gene which was also on the list of genes with bidirectional effects, Tables 1-2) is being investigated in a phase III Randomized Clinical Trial for Achondroplasia.^41^ Binding of CNP to its receptor (*NPR2)*, triggers endochondral and skeletal growth via cGMP production.^42^ Furthermore, even wild-type rhesus monkeys exhibit increased growth velocity when treated with CNP.^43^ Loss-of-function mutations in *NPR2* are responsible for dwarfism in mice and a lack of an intracellular cGMP response to CNP in cultured chondrocytes.^44^

In order to explore the effect of gain of function mutations in *NPR2* on height, which should mimic the effect of adding exogenous CNP, we assessed the function of multiple missense mutations across the full spectrum of NPR2 activity. We selected 39 variants for functional analysis: 9 *NPR2* protein-altering mutations that were present in UKBiobank ISS individuals, 7 NPR2 missense mutations present in UKBiobank individuals with height > 1.2 SD, 5 *NPR2* variants in individuals with a mean height around 0 (in SDs), and 4 additional *NPR2* mutations with higher carrier number to complement the allelic series. Additionally, we tested 13 *NPR2* mutations that were reported in the literature for ISS and/or a more severe recessive form of short stature (Methods and Supplementary Table 8). To distinguish functional *vs.* neutral *NPR2* missense variants, we performed a series of experiments in cellular models (Methods and Fig. 3A). Constructs were tested in triplicate and the mean and SD of the functional readout (cGMP) was reported (Methods and Supplementary Table 8). We first confirmed the dynamic range of the assay: 10 out of 11 previously reported pathogenic variants had mean cGMP < 0.20 relative to wildtype (wt), 2 out of 3 previously reported gain of function variants showed a mean cGMP of > 1 relative to wt (Supplementary Table 8). From the list of ISS patients found in our study, 10 out of 12 tested mutations were confirmed as low activity (mean cGMP < 0.2 relative to wt). (Fig. 3B, Supplementary Table 8). There was a significant association with increasing cGMP levels and increasing height (β=0.91 per 100% change in cGMP relative to wt, P=2.7×10^−7^; Fig. 3C). This assay data served as a better classifier for height than a predictor (Combined Annotation Dependent Depletion or CADD^45^, (β=−0.02, p=0.04 for CADD unit change score; Fig. 3D).

**Figure 3.**
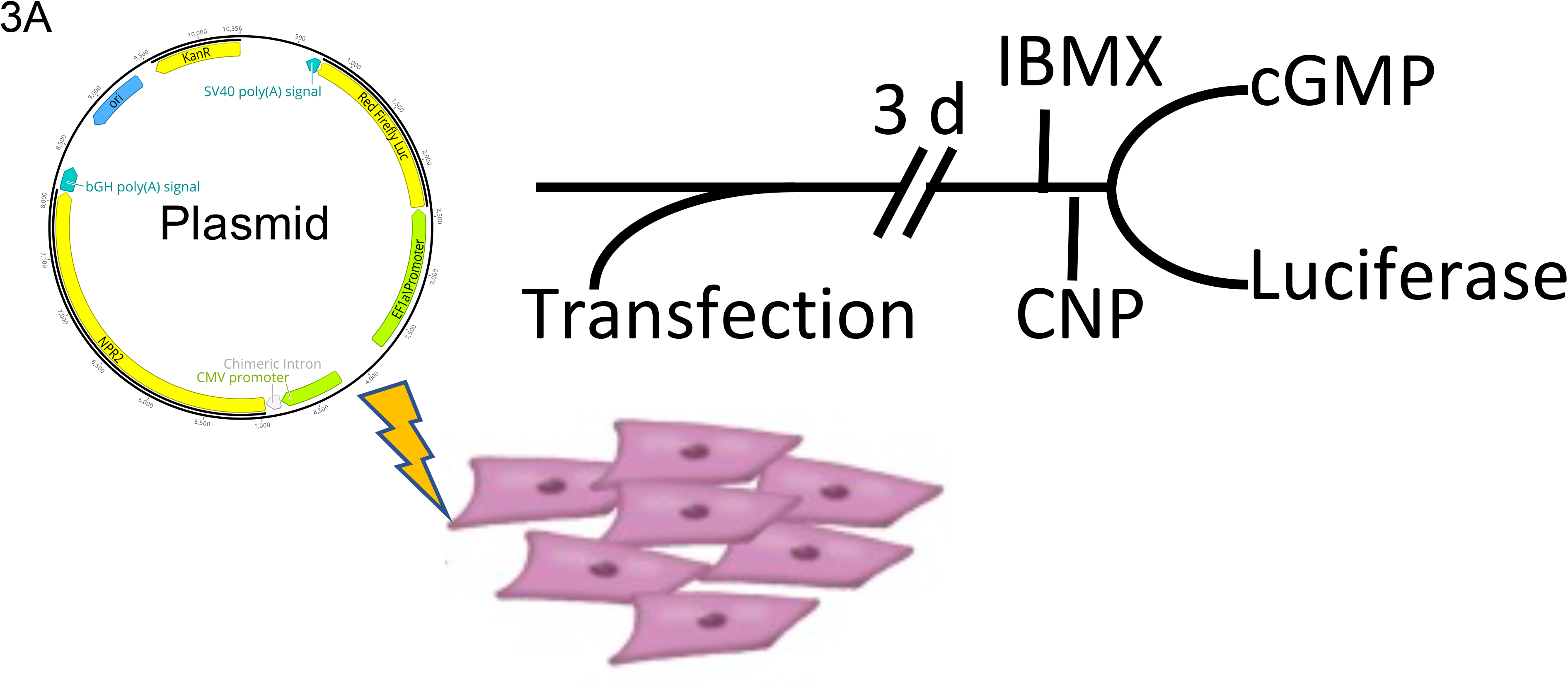
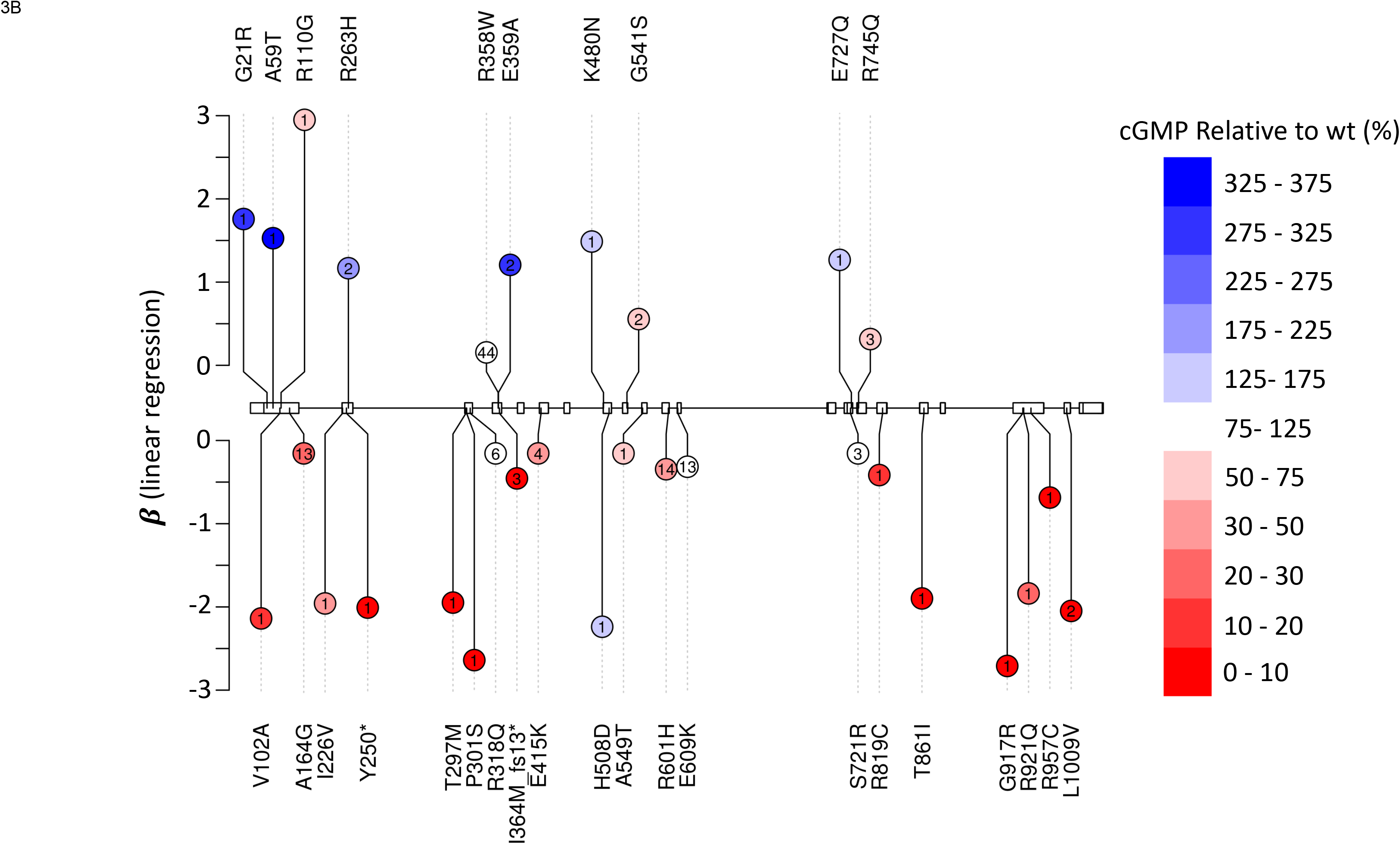
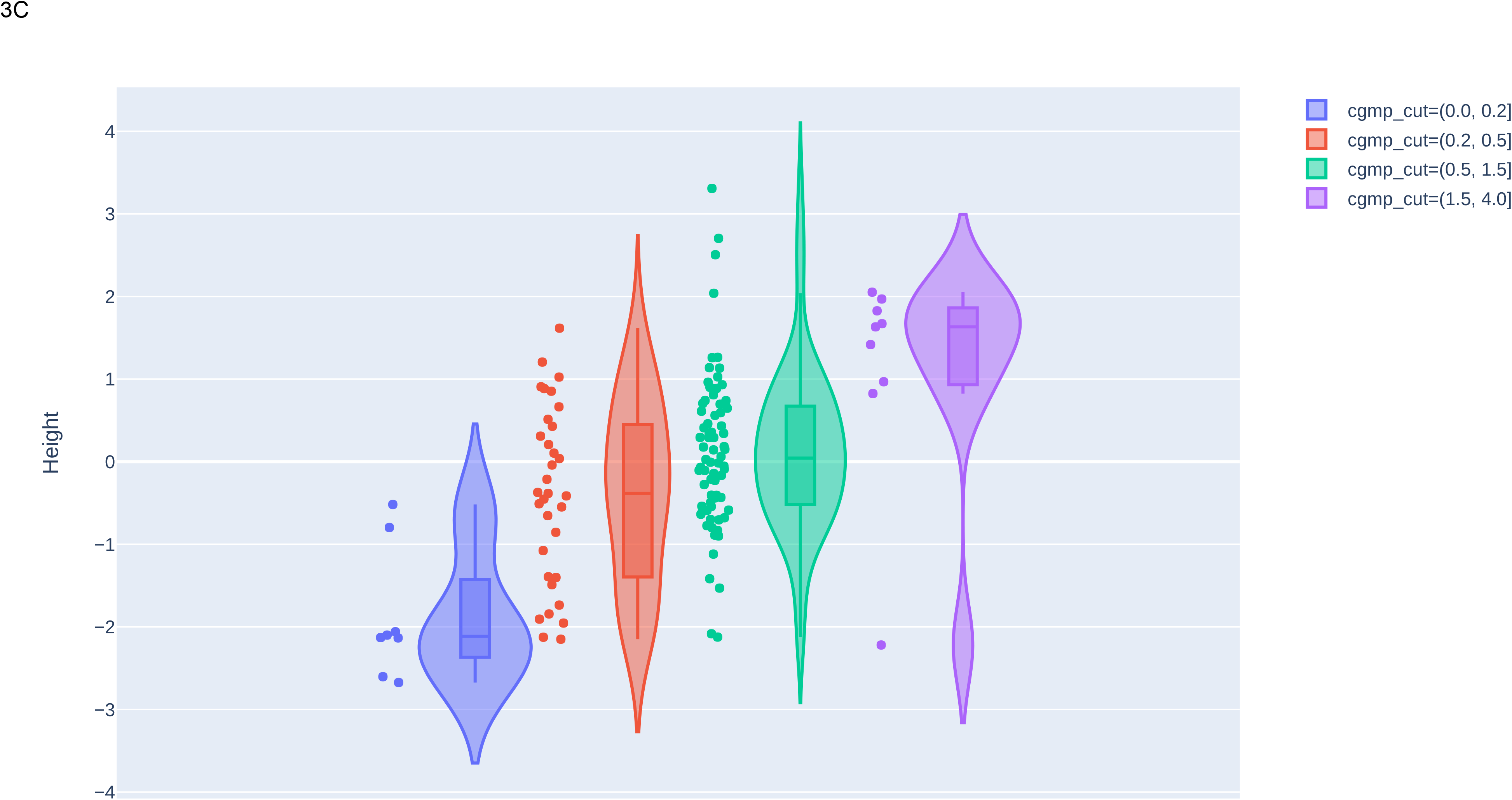
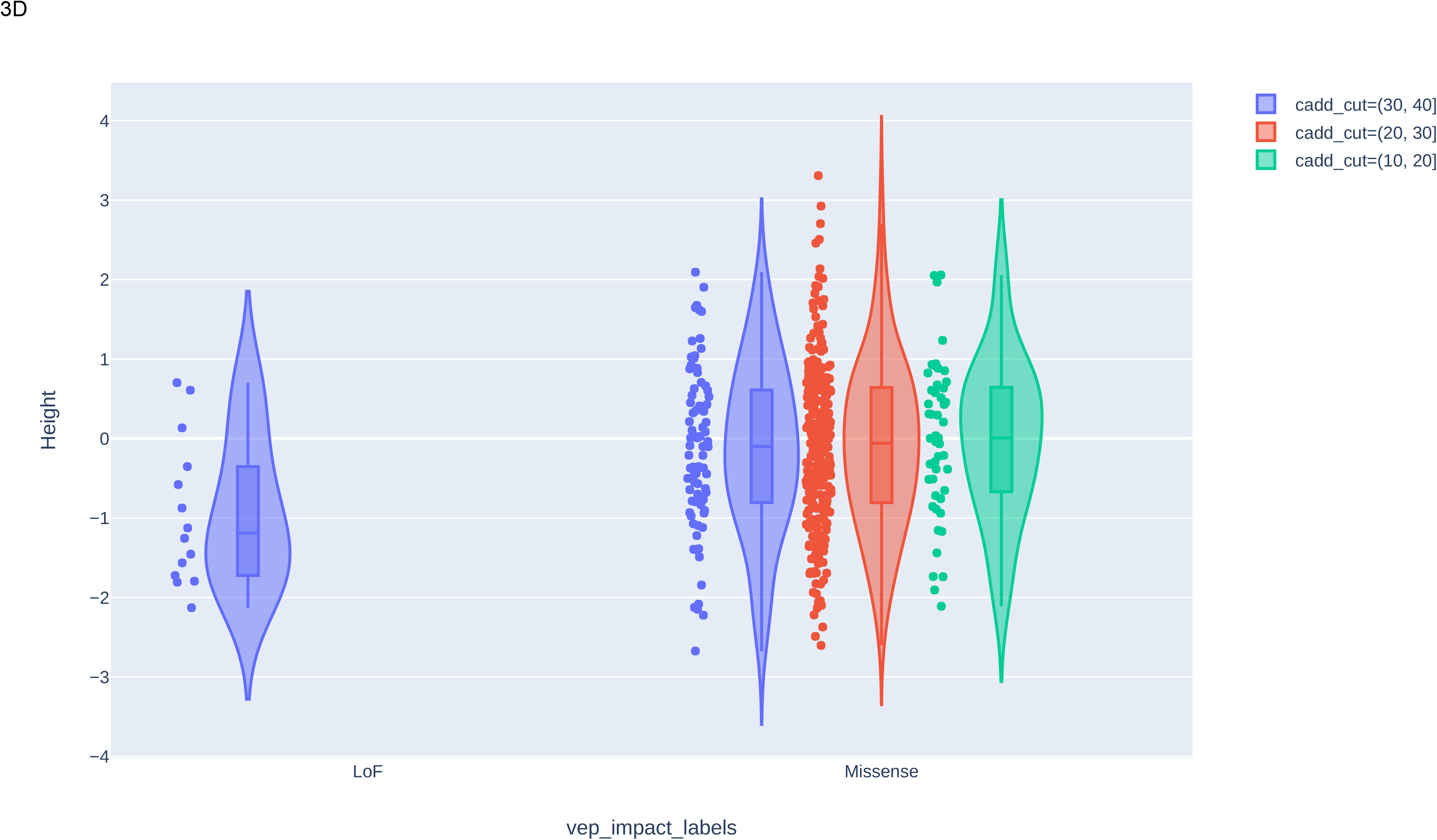
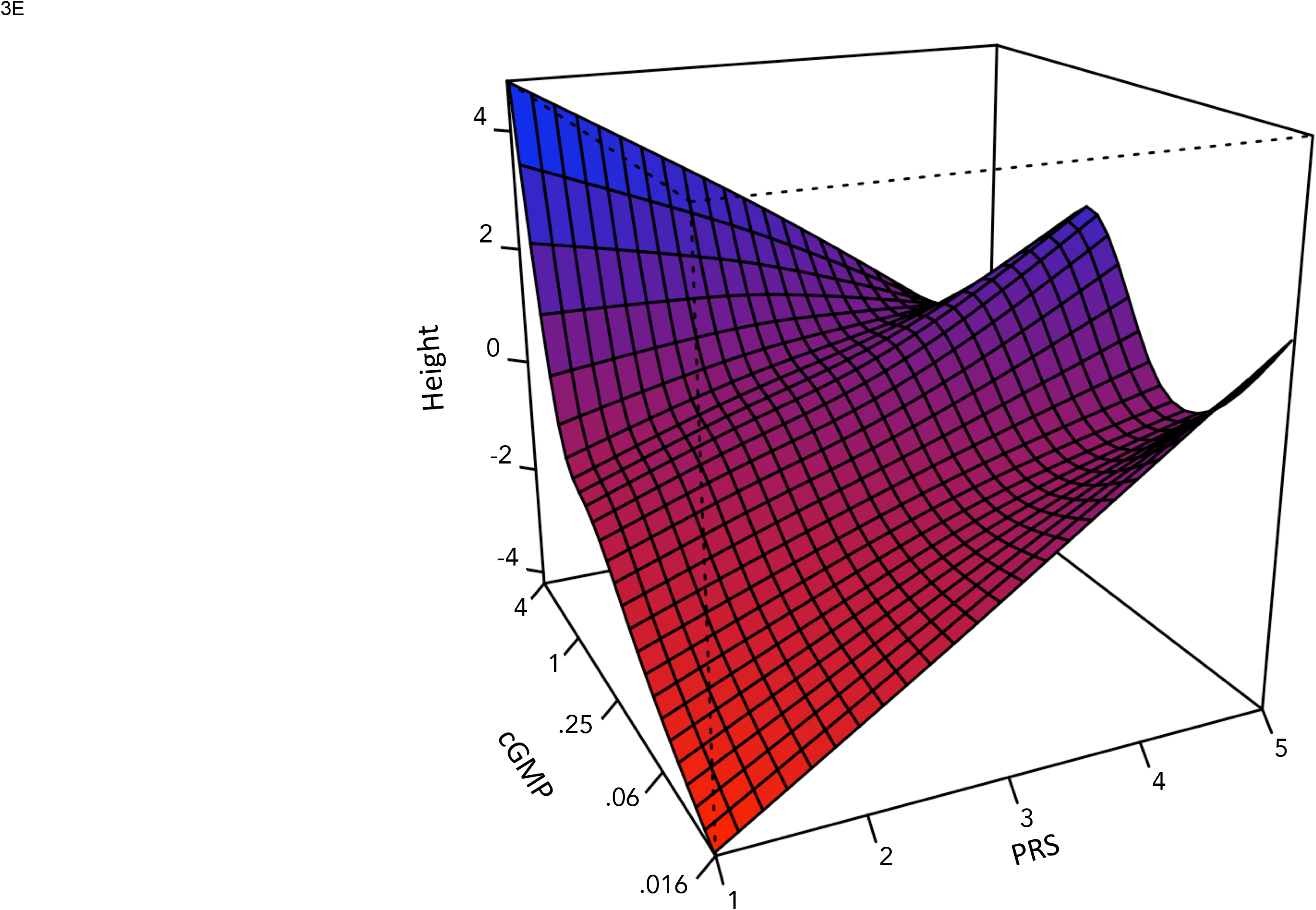
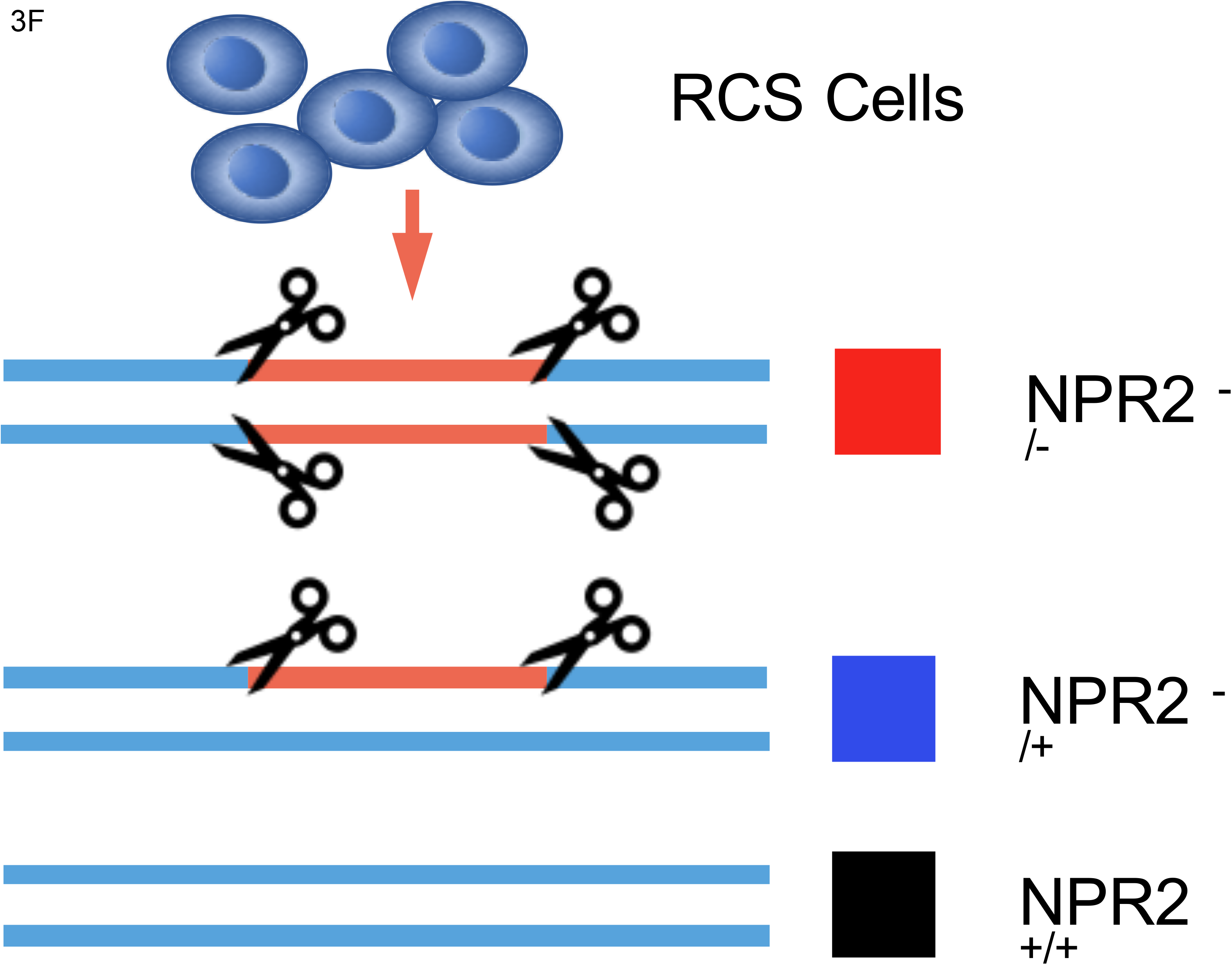
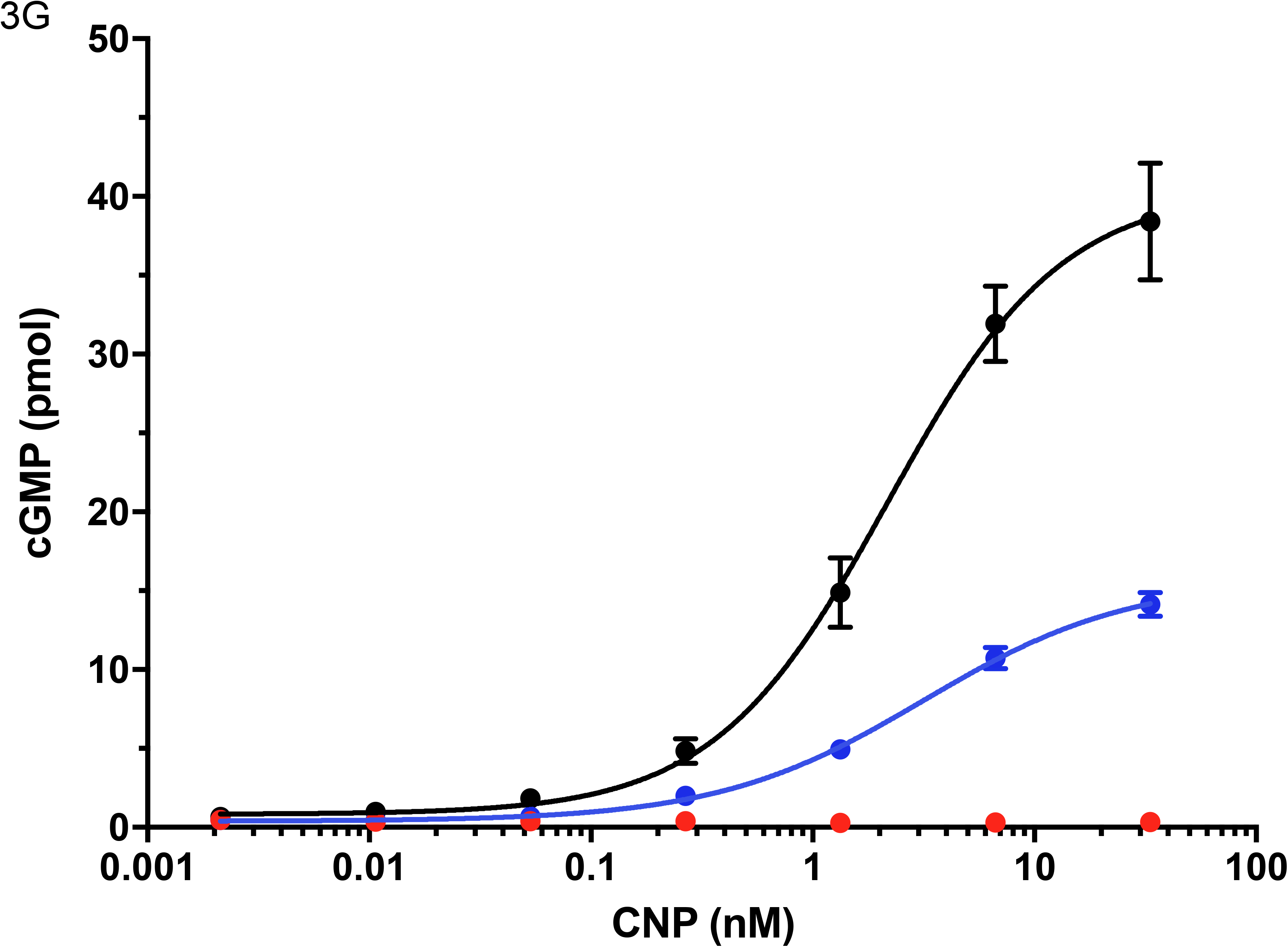
Functional effects of NPR2 protein coding variants. A) Schematic for evaluation of missense variants. The bicistronic plasmid with a CMV promoter driving production of NPR2 variants and an EF1a promoter driving production of red firefly luciferase was transfected into HEK293T cells. Three days post transfection cells were treated with IBMX, incubated with CNP, and analyzed for luciferase and cGMP activity. cGMP activity was normalized of variants relative to luciferase, n = 3 biologically independent samples. B) Diagram of NPR2 protein altering variants identified in the UKBiobank study. X-axis indicates genomic position. Y-axis indicates effect size (β) of a linear regression of that variant on height in standard deviations. Each circle represents a different variant with the number of carriers inside the circle, amino acid change of the variant is labeled at the end of the circle. Color code indicates cGMP activity of variants relative to luciferase. C) Violin plots representing height of carriers of the NPR2 variants tested in functional analyses. Box-plots inside violin-plots represent the median and inter-quantile range of height in the group. Whiskers are 95% CI. Color categories represent binned cGMP functional readout. D) Violin plots representing height of carriers of all *NPR2* variants found in the study. Box-plots inside violin-plots represent the median and inter-quantile range of height in the group. Whiskers are 95% CI. Color categories represent binned computer algorithm scores (CADD). E) Predicted measured height (Z-transformed) as a function of their PRS and the percent of wild-type cGMP observed for their *NPR2* variant. The color maps to height (blue= highest, red=lowest). F) Schematic for generating NPR2 deficient cells using CRISPR/Cas9. G) cGMP dose response to CNP treatment with *NPR2* knockout (red), heterozygous knockout (blue) or wild-type (black) cells, n = 3 biological independent samples; mean ± s.d.

One interesting outlier was identified carrying the *NPR2* variant R110G. While this variant showed a modest loss of function (cGMP = 0.70), the individual carrying it was very tall (height=3.10 SDs). A closer examination of this individual revealed that they were in the 99.85^th^ percentile of the PRS for height (Supplementary Fig. 17). This prompted us to examine PRS groups in *NPR2* carriers of the variants we functionally tested. We grouped *NPR2* carriers by their PRS group and their residual *NPR2* activity. Despite having a small number of individuals in each group, as shown before there was an additive relationship between PRS and *NPR2* functional variants on height (Fig. 3E, Supplementary Table 9). This additive relationship was consistent with the combination of multiple risk factors of PRS and rare variants with strong effects

### Phenotype rescue

In order to test whether a CNP-analog can rescue a LoF mutation in *NPR2*, we examined the response to CNP of cells engineered to have heterozygous or homozygous LoF mutations. We measured the amount of cGMP produced by CNP stimulation in CRISPR/Cas9-mediated knockout chondrocytes (Methods and Fig. 3F). cGMP production by NPR2 triggers a signaling cascade via activation of protein kinase cGMP-dependent 2 (*PRKG2*). Previous activation data reports cGMP EC_50_ in the range of 40 to 360nM for activation of *PRKG2*.^23,46,47^ In the heterozygous *NPR2* knockout cells, a CNP dose >0.163nM is able to achieve an intracellular concentration exceeding the EC_50_ range for PRKG2 activation (Fig. 3G). Whereas, in wild-type cells a CNP dose of 0.040nM is able to achieve the same cGMP concentration. These results demonstrate that CNP supplementation can achieve the cGMP levels necessary for PRKG2 activation and growth in cells with loss-of-function mutations of *NPR2*.

### Generalizability to other diseases

Next, we wanted to assess the generalizability of our BEST strategy for other human traits or diseases. To address this, we manually identified 98 genes in HGMD that showed evidence of bidirectional effects for different gene variants (Online Methods, Supplementary Table 10). We then calculated the odds of clinical trial approval for those BEST target-indication pairs using a recently published database of clinical trial outcomes.^48^ Finally, we compared those odds to the odds of targets with purely unidirectional evidence from OMIM and targets with evidence from GWAS (Online Methods, Supplementary Table 11). We found that overall, the BEST target-indications that have reached Phase I have higher probability of successful transition to Approval as compared to other non-BEST target-indications across all MeSH similarity thresholds (Fig. 4A) and it reached a maximum OR=3.81 95%CI [2.81 - 4.74] at MeSh >= 0.83. Importantly, that probability was significantly higher than that of other sources of genetic evidence (Fig. 4B, OR=2.12 95%CI [1.73 - 2.52], OR=1.12 95%CI [0.82 - 1.46] for OMIM unidirectional and GWAS targets respectively). BEST target-indications showed significant improvement in the probability of successful transition at all steps of the clinical development as compared to non-BEST target-indications (Fig. 4B). Interestingly, the phase where BEST target-indications showed the strongest effect was the transition from Phase II to Phase III (OR=1.46 95%CI [1.25 - 1.66]), a phase that is generally used as proof of concept or initial clinical efficacy.

**Figure 4.**
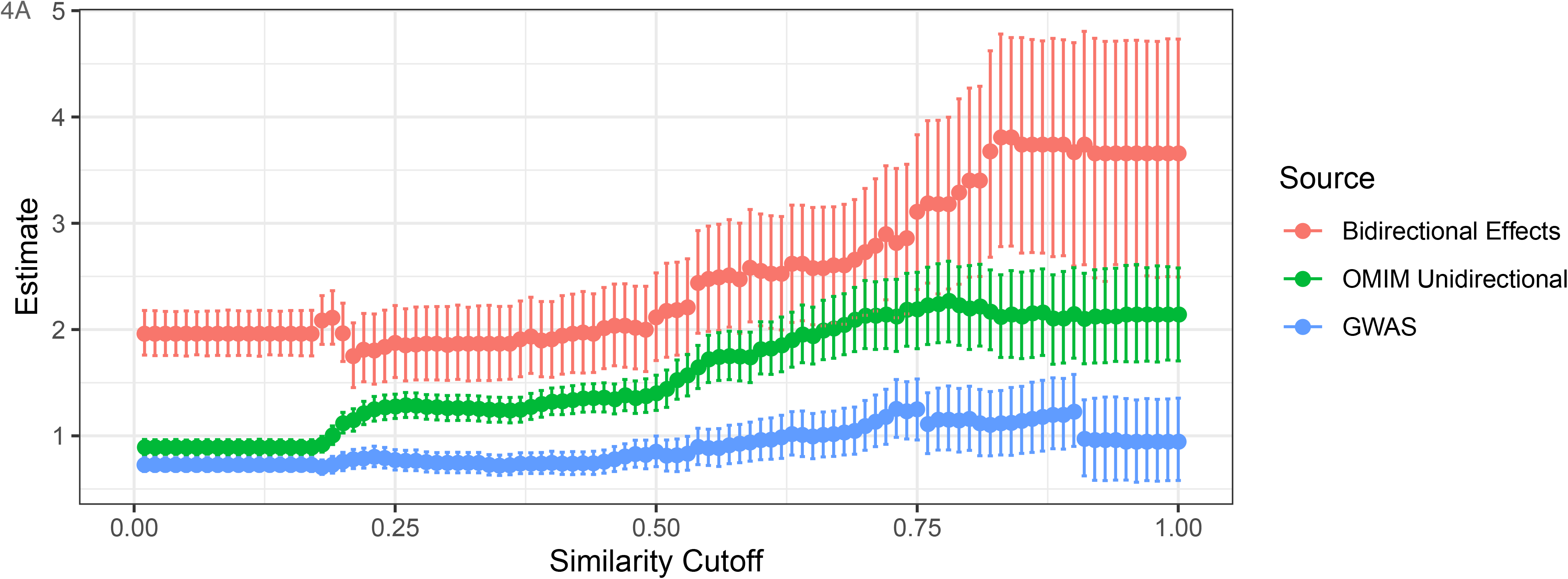
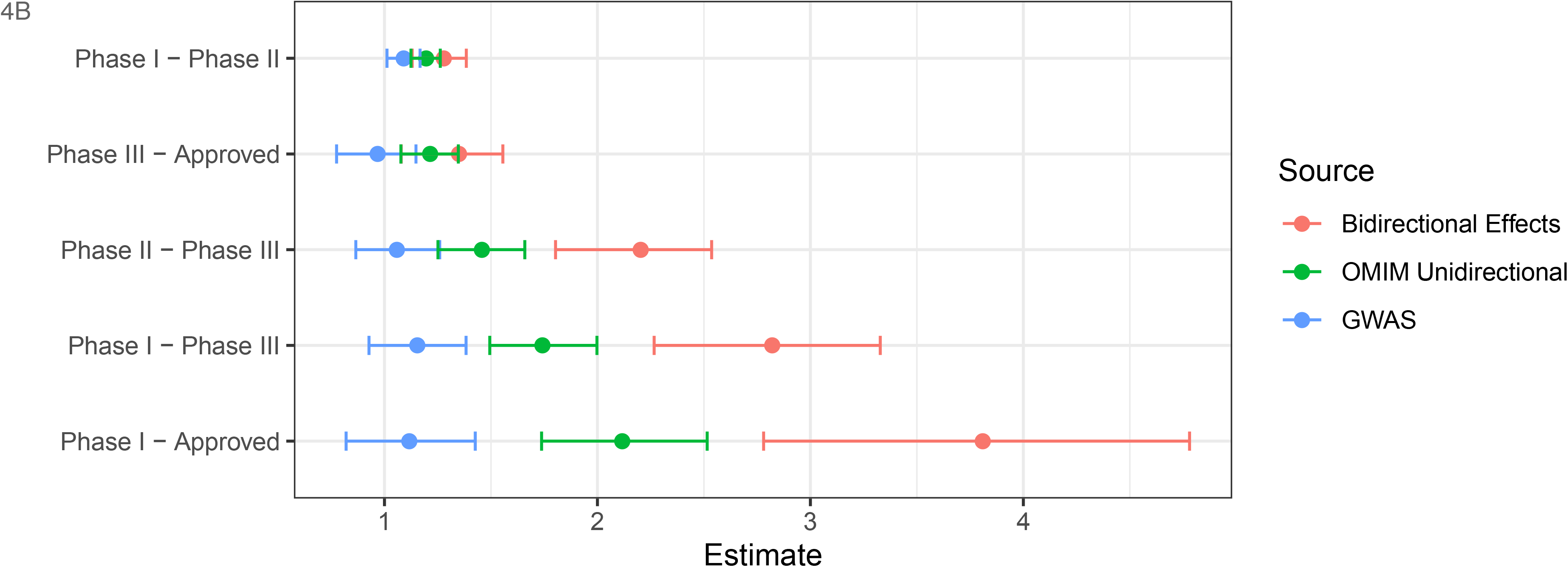
Generalizability of bidirectional effects for increasing probability of success on other indications. A. Estimated odds ratio for transition from Phase I to Approved. We calculated the odds ratio for transition from Phase I to approval and 95% confidence, for the subset of target-indications with genetic support from GWAS, OMIM associations with unidirectional effects, and bidirectional effects curated from HGMD. This process was repeated using all thresholds of MeSH term similarity from 0 to 1 by increments of 0.01. Plotted points represent the estimated odds ratios and the error bars represent the 95% confidence intervals. Bidirectional effect supported data is shown in red, OMIM unidirectional support is shown in green, and GWAS support is shown in blue. B. Estimated odds ratio for transitions between various phases at MeSH similarity >= 0.83. We calculated the odds ratio for transition and 95% confidence, for the subset of target-indications with genetic support from GWAS, OMIM associations with unidirectional effects, and bidirectional effects curated from HGMD. Plotted points represent the estimated odds ratios and the error bars represent the 95% confidence intervals. Bidirectional effect supported data is shown in red, OMIM unidirectional support is shown in green, and GWAS support is shown in blue.

## Conclusions

In this study, our goal was to identify genes most likely to be drug targets for ISS by showing mutation bidirectionality (GoF and LoF), as well as GWAS effects with moderate effect sizes. We identified a set of five height genes with bidirectional effects and characterized the effect of these mutations across different height PRS backgrounds. Height has been used as model for common complex traits and diseases and its genetics are characterized by a large polygenic architecture. Despite this polygenic nature, there are critical genes where single LoF/GoF variants have large effects with magnitudes approaching the aggregated effects of thousands of common variants with small effects (as calculated by PRS). We demonstrate that genetic insults to these five genes contribute not only to rare skeletal malformations (Table 1, Supplementary Table 6), but also to common forms of short stature. Additional evidence from both human and animal models support the notion that the *NPPC*-*NPR2*-FGFR3-SHOX pathway is a key modulator of growth (Table 2). In the same way, *IGF1R*, a key member of the growth hormone receptor pathway, points toward a second group of genes that would include IGF-I which is currently a short stature therapy for Growth Hormone non-responsive patients.^49^

The additive effects of common genetic variation (i.e. PRS), predicted 20.1% of the variance in height in our dataset. These effects appeared to have similar magnitudes on the carriers of rare coding variation of genes as compared to non-carriers. This observation indicates that PRS may be a strong contributor in the differences in penetrance of rare pathogenic variants (especially in models of haploinsufficiency such as the ones described here). Supporting this idea, we saw that two of eight *NPR2* variant carriers with low NPR2 activity had a low-normal height. One of them was in the 4^th^ PRS quintile and the other was in the 5^th^ PRS quintile, the remaining 6 were in quintiles 1^st^, 2^nd^ and 3^rd^ (Fig. 3E, Supplementary Table 9). This data suggests that most ISS individuals possessing mutations in *NPR2* may also have a polygenic background that made them more susceptible to the pathogenic effect of losing *NPR2* activity. This observation could be interpreted as supporting for the omnigenic model,^50,51^ where core genes would define a module that when disrupted, has large effects. Similarly, the omnigenic model predicts that the effect of core genes is modulated by multiple weaker common genetic variants driving regulatory networks. However, it is evident that in our data PRS has a strong effect on height even in non-carriers of the five genes we identified.

It is possible that that other genes will appear as key contributors of height and ISS as sample sizes increases. Our study was 63% powered to identify a gene set association that explained the proportion of variance we observed in the set of five genes we describe. A follow-up study would require over 400,000 samples to have 80% power to identify individual genes at exome-wide significance levels (See Methods).

Our results have important implications not only for drug discovery but also for drug repurposing. The presence of genes driving disease in the general population suggests the possibility of tailoring therapies to the small fraction of individuals with gene defects. Also, the fact that such genes have allelic series with bidirectional effects and additive effects across PRS groups, suggests that such tailored therapies may be effective for any patient irrespective of the genetic lesion. Importantly, we have taken initial steps to show that having bidirectionality increases the probability of success (4-fold) in clinical trials above and beyond just having human genetic evidence (2-fold).

As mentioned earlier, CNP-analogs are being tested as a treatment for Achondroplasia (incidence 1 in 10,000).^41^ Here we demonstrate that carriers of variants in any of the five genes are at ~ 3-fold increased risk for ISS and account for 6.7% of the total ISS population.

Furthermore, we showed dose-dependent rescue of *NPR2* signaling in a cell model of *NPR2* haploinsufficiency after adding exogenous CNP. These results support the idea that these CNP-based treatments could be effective in NPR2 haploinsufficient patient populations if the right dose can be delivered. Further, our results showing a significant correlation of cGMP levels and height in *NPR2* carriers of the general population suggest that targeting this receptor with CNP analogs could be an effective therapy for all ISS individuals.

In summary, we highlight a strategy to identify genes driving disease by leveraging a combination of human genetic databases and validation using large-scale exome sequencing studies followed by functional studies. We also highlight potential opportunities for drug target discovery and repurposing in this type of genetic targets.

## Online content

Supplementary Figures and Tables are available online.

## Acknowledgements

We thank all participants of the UK Biobank project for sharing their data. We also thank Dr. Pavel Krejčí (Masaryk University) for providing us access to the Rat Chondrosarcoma Cell line.^52^ We are grateful to Drs. Emily King and Wade Davis for their advice and assistance calculating the odds of drug approval based on the support of genetic evidence.

## Author contributions

KE, LC, JL designed the study. SF, MT, HN, AL, GY designed and executed the functional experiments. KY, KE, AW, CB, TS, WC obtained and worked with the exome sequence data. DW provided access to the rat chondrosarcoma cell line and provided input for the experiments. KE drafted the manuscript and all co-authors contributed to it.

## URLS. Additional information

UK Biobank website and data access http://ukbiobank.ac.uk/

Genomics PLC wecall: https://github.com/Genomicsplc/wecall

Hail: https://github.com/hail-is/hail/releases/tag/0.2.16

SKAT: https://www.hsph.harvard.edu/skat/

Genetic-evidence-approval: https://github.com/AbbVie-ComputationalGenomics/genetic-evidence-approval.

## Methods

### Human databases

#### GWAS

We extracted 2,067 non-repeating closest genes for each of the 3,290 independent genetic variants reported by a large GWAS meta-analysis of height using ~700,000 individuals (Supplementary Tables 1 and 2).^18^ We considered this the most comprehensive GWAS for height.

#### HGMD

We queried the “allmut” table from HGMD version v2019_2 looking for all pathogenic variants labelled as “DM” having either “short stature” and “tall stature or overgrowth” in the same genes (Supplementary Table 2).

#### OMIM

The list of OMIM genes related to growth disorders was previously described and was created using the keywords: short stature, overgrowth, skeletal dysplasia, brachydactyly(Supplementary Table 2).^20^

### Samples

#### UKBiobank

Data was accessed under application 41232. The demographics and patient characteristics for the 50,000 exome sequenced individuals has been previously described.^53^ In short, individuals from the UK aged 45 to 75 were invited to participate. A baseline questionnaire and several measurements were taken. Electronic health records were also available. Standing height (field id 50) was measured to all individuals as part of the baseline assessment (Supplementary Table 3). We used GNU parallel to rapidly extract individual phenotype fields to CSV files.^54^

### Sequencing and variant calling

Sequencing protocol and details are described elsewhere.^53^ In short, exomes were captured using a modified version of the IDT xGen Exome Research Panel 81 v1.0 and sequenced in pooled multiplexed samples using 75 base pair paired-end reads with two 10 base pair index reads on the Illumina NovaSeq 6000 platform using S2 flow cells. Reads were aligned to the GRCh38 genome reference with BWA-mem2. The WeCall variant caller was used to generate gVCF files (See URLs). A PVCF was created using the GLnexus joint genotyping tool.^55^

### Software

Genetic data analysis was performed on Hail 0.2.16 running on Apache Spark version 2.3.4.

### Exome sequencing data

We downloaded UK Biobank exome sequencing genotypes in Plink format for 49,960 individuals and 10,448,724 variants (“SBP” data) in May 2019.

We annotated and filtered individuals with data provided by UK Biobank. Specifically, we included individuals with the genetic ethnic grouping “Caucasian” (field 2206), no sex chromosome aneuploidies (field 22019) (effectively removing any classical Turner syndrome patients) and with a PCA-corrected heterozygosity between 0.17 and 0.21 (field 22019). After applying these filters, our dataset contained 41,190 individuals.

We removed related individuals by excluding samples meeting the criteria for relatedness and kinship set by UK Biobank (2nd degree relatives or closer) and that are listed in fields 22011, 22012 and 22013. After applying this filter, our dataset contained 35,990 individuals.

We computed sample and variant QC metrics and applied additional filters that excluded sample with more than 80,000 non-reference variants, more than 200 singletons, a het/hom ration of less than 1.3 or more than 1.85, and transition/transversion ratio of more than 2.5, a call rate of less than 0.985 or a proportion of heterozygous sites in sequencing data that are also heterozygous in chip data of less than 0.98. We also excluded variants that had a call rate of less than 0.99 or a Hardy-Weinberg p value of less than 10^−10^.

We annotated variants using the Variant Effect Predictor (VEP)^56^ and extracted rare high impact variants. We defined those as variants with a minor allele frequency of less than 1% and whose impact as predicted by VEP is either high (protein truncation) or moderate (missense). We observed that a subset of those high-impact variants had an unexpectedly high allele frequency compared to the allele frequency of non-Finnish European (NFE) individuals in the genome aggregation database (gnomAD).^57^ We removed those variants if the observed allele count in our dataset was significantly different to what would be expected based on the NFE allele frequency in gnomAD (p < 10-7, binomial test). After applying those filters, our dataset contained 581,392 variants. The assignment of variants to genes was accomplished by using canonical VEP gene models.

### Quality control

After applying the filters described above, our exome sequencing dataset contained 34,284 individuals and 10,173,231 variants. There was no apparent clustering of the post-filtering European ancestry individuals by genotyping eigenvectors (Supplementary Figs. 18-19). The sample call rate was between 0.999941 and 1.0 (median: 0.999984) and the variant call rate was between 0.989909 and 1.0 (median: 1.0) (Supplementary Figs. 20-21).

We observed between 37 and 111 (median: 68) rare high-impact variants per individual (Supplementary Figure 22) and between 0 and 36 (median: 9) singletons per individual (Supplementary Figure 23). Most individuals had a rare high-impact variant in less than 1% of genes (Supplementary Figure 24).

### Gene set analyses

We constructed a list of 2,355 genes (Supplementary Table 3) that had evidence from GWAS and/or HGMD/OMIM. To assign a GWAS signal to a gene, we used the latest meta-analysis for height and took the list of 3,290 predicted independent variants (Supplementary Table 1).^18^ Then we mapped the closest gene to each one of those lead variants and removed duplicates. That list of genes was compiled together with genes reported on HGMD and OMIM and genes were assigned one to five different annotations: GWAS, HGMD_SHORT, HGMD_TALL, OMIM-OVERGROWTH, OMIM-SHORT (Supplementary Table 2).

We then queried this list of genes into the UKBiobank exome dataset. Here 1,970 genes had at least 20 carriers (combined rare missense and Lof) and were considered for the analyses. Genes had a mean of 204 rare protein coding variants (Supplementary Fig. 20).

### Statistical association analyses

Standing height measurements were first split by gender and then normalized to standard deviations adjusted for age and first five principal components using a linear regression implemented on the R package. Linear and logistic regression analyses for single variants were computed using Hail 0.2.

We grouped rare missense and loss of function variation per gene using GENCODE canonical transcripts. We applied SKAT and burden tests on missense variants and predicted protein truncating variants (PTVs) with a minor allele frequency < 0.01% using Hail 0.2.

For the association analysis on gene-sets and ISS we grouped genes in the following way: Bidirectional: Genes present in HGMD_SHORT, HGMD_TALL, OMIM-OVERGROWTH, OMIM-SHORT. Tall: Genes present in HGMD_TALL or OMIM-OVERGROWTH but not in HGMD_SHORT or OMIM _SHORT; Short: Genes present in HGMD_SHORT and OMIM _SHORT but not in HGMD_TALL or OMIM-OVERGROWTH.

### SNP chip data and polygenic risk scores

We used summary statistics from a large GWAS of human height to compute a height polygenic risk score (PRS) for UK Biobank participants.^20^ Specifically, we downloaded UK Biobank imputed SNP chip genotypes in BGEN format for 487,409 individuals. We exported genotypes for 2,540,786 variants for which GWAS summary statistics were available to Plink format. We then used LDpred^58^ version 1.0.7 to compute a height PRS for 49,796 individuals for which exome sequencing data was available. For the coordination step of LDpred we used a linkage disequilibrium reference panel of 2,000 randomly selected UK Biobank participants of European ancestry for which exome data was not available. For variant filtering, generating SNP weights and PRS scoring we used default LDpred parameters. Linear regression analyses were used to estimate the effect of PRS as a categorical variable (1-5) on height. Logistic regression analyses were used to estimate the effect of PRS as a categorical variable (1-5) on ISS, groups with low number of ISS cases (standard error > 100) were removed.

### Selection of NPR2 variants for Functional assays

A total of 39 NPR2 protein coding variants were selected to investigate their functional effect at a cellular level (Supplementary Table 7). Fourteen of those variants had been previously described in the literature and aren’t explicitly found in our study. Seven variants were found in our exome dataset in single individuals with height > 1 SD above the mean and were labeled as potential gain of function. Nine variants were found in individuals with height < 2 SD below the mean and were labeled as potential loss of function. Three variants with neutral effect on height were tested. Four variants with MAF <0.1% (p.A164G, p.R601H, p.E609K, p.R358W) were added to complement the allelic series.

### Transfection Functional assays and Assay for cGMP analyses

HEK293T cells were cultured in Dulbecco’s modified Eagle’s medium (DMEM) supplemented with 10% fetal bovine serum, penicillin/streptomycin and Glutamax (Gibco) at 37 °C with 5% CO_2_. Transfection was performed in triplicate using a 96-well Shuttle nucleofector (Lonza). Briefly, 250 ng of plasmid per 2 × 10^5^ cells was transfected using Amaxa solution SF (Lonza) and program CA-138. Three days after transfection, cells were then incubated in DMEM containing 0.75 mM IBMX (3-isobutyl-1-methylxanthine) (Enzo life sciences) for 15 min. The cells were next treated with 6.67 nM of BMN 111 (CNP)^43^ and incubated for another 15 min. The reaction was terminated with 40 μl of lysis solution, and the cGMP concentration was measured by a competitive enzyme immunoassay (Molecular Devices, R8075). Luciferase levels were measured using the ONE-Glo Assay (Promega) and the results are presented as the cGMP/luciferase normalized to the wild-type plasmid transfected cells. For the dose-response curves, the CNP concentrations assayed in triplicate were 33.33, 6.67, 1.33, 0.27, 0.053, 0.0107, and 0.002133 nM. The cGMP concentrations were assayed as above and cellular concentrations calculated assuming 1.6E5 cells/well with a diameter of 35μm.

### PRS and NPR2 functional data modeling

For all individuals carrying a rare variant in *NPR2*, for which functional assay data was available, we fit a generalized additive model to predict measured height (Z-transformed) as a function of their PRS for height (Z-transformed) and the percent of wild-type cGMP observed for their *NPR2* variant (log base 2 transformed). Models were fit using the gam() function from package mgcv v1.8-31 in R version 3.6.1. A gamma parameter of 0.3 was chosen to minimize the amount of smoothing. Values smaller than this resulted in unstable models that produced unrealistic height predictions.

### CRISPR/Cas9 knockout of *NPR2* in rat chondrosarcoma chondrocytes

The swarm rat chondrosarcoma (RCS) cell line^52^ was cultured with DMEM containing 10% fetal bovine serum, penicillin/streptomycin, and Glutamax (Gibco) at 37 °C with 5% CO_2_. RCS cells were harvested with TrypLE Express Enzyme (Life Technologies) and transfected with RNPs formed with guide RNA containing ATTO 550 labelled tracrRNA (IDT 1075928) and crRNA targeting NPR2, with PAM in bold:

gRNA1 5ʹ-CCCACTACTTCACCATCGAG**GGG**-3ʹ
gRNA2 5ʹ-CATTACACGGGCACTTCAAT**TGG**-3ʹ

Briefly, equimolar ratios of crRNA and tracrRNA were annealed and 12pmol was complexed with 104pmol HiFi Cas9 (IDT 1081061) according to the manufacturers protocol. 2E5 cells were transfected with 500ng of GFP mRNA (TriLink Biotechnologies, cat# L-7201) or RNP using a 96-well Shuttle nucleofector (Lonza) with Amaxa solution SF (Lonza) and program CA-150. Post transfection, each well was split and seeded into two wells of a 96well plate. Three days after transfection, single cells were sorted into 96 well plates containing 200μl of media using a FACSMelody cell sorter (BD Biosciences); the plates were monitored biweekly by imaging (Cell Metric, Solentim) when confluent cells were expanded into 6 well plates and genotyped.

Genomic DNA was extracted using Quanta Extracta solution (Quanta Biosciences, cat #95091). DNA amplicons covering the gRNA target region were amplified and analyzed by PacBio sequencing following the manufacturer’s instructions (Pacific Biosciences, 101-791-800 Version 01). Amplicon libraries were prepared using primers with universal sequences and the locus specific primers FW 5ʹ-CATCCCTGCTACTGGTGGTG-3ʹ and REV 5ʹ-CTCCACCACCAAACCTGAACT-3ʹ.

Sequencing was performed using the Sequel System using the SMRT Cell 1M v3 and acquiring 10h movies. Demultiplexing and circular consensus sequence analysis and FASTQ generation was performed using the SMRTlink 6.0 software (Pacific Biosciences). To analyze amplicon sequencing, Geneious Prime 2019.0.4 software was used for quality assessment, quality filtering, reference alignment, and calculating indels (Supplementary Fig. 21).

### Power analysis

We used the PAGEANT^59^ method to estimate the power of the current study using the SKAT method on the following two scenarios: a. Our study design involved an alpha=0.002 (adjusting for 25 gene set tests), we considered a variance explained of 0.09%. b. A study would require 400,000 samples to identify single genes at α=2.5e-6, a maximum of 1000 causal genes, and considering 5% of variance explained for protein-altering variants genome-wide.

### Bidirectional screen for broader indications

The HGMD allmut table v2016.4 was used as the source for associations to identify bidirectional effect associated genes. The table was grouped by each combination of gene and disease. Any genes with less than two unique disease associations were excluded from further analysis. The remaining dataset contained 23,754 distinct gene-disease pairs between 4427 genes and 12302 diseases. This table was manually curated to identify genes with at least 2 associations that appeared to different directions of effect on a common underlying phenotype. For example, PSCK9 was identified as a bidirectional effect gene due to associations with “High LDL cholesterol” and “Low LDL cholesterol”. KCNJ2 was identified as a bidirectional effect gene due to associations with “Long QT syndrome” and “Short QT syndrome”. We also required that the disease field represent an actual disease or disease associated endophenotype such as a blood metabolite. Thus, associations between genes and molecular, cellular, or benign phenotypes (e.g. increased and decreased enzyme activity, cell permeability, or pigmentation) were also excluded.

### Probability of Drug Approval Analysis

Analysis of the odds of drug target-indication approval was conducted in R/3.5.2 following the methods of King et al.^48^ Since some OMIM disease genes were also associated with bidirectional phenotypic effects, we first reran their analysis after removing those genes to estimate the odds of approval for drugs that target OMIM genes with unidirectional effects.

We then repeated this analysis with our set of gene-trait associations with bidirectional effects replacing the OMIM genes in the original King et al. data set. We called the function annotate_target_indication_table_with_genetic_evidence() to match the set of target-indications with the best supporting genetic evidence. We then used the function replicate_table1() to estimate the odds of transition between different phases of clinical development using a range of MeSH term similarity thresholds (0 to 1 by increments of 0.01) to link drug indications with genetically associated diseases or traits.

The maximum odds of transition from Phase I through approval occurred for indications with MeSH similarity greater than 0.83.

