## Supplementary Information for "Identifying therapeutic drug targets for rare and common forms of short stature"

Supplementary Tables.

Supplementary Table 1. List of lead SNPs from GWAS meta-analysis of height (Online)

Supplementary Table 2. List of height genes and their respective annotations (Online)

### Supplementary Figures

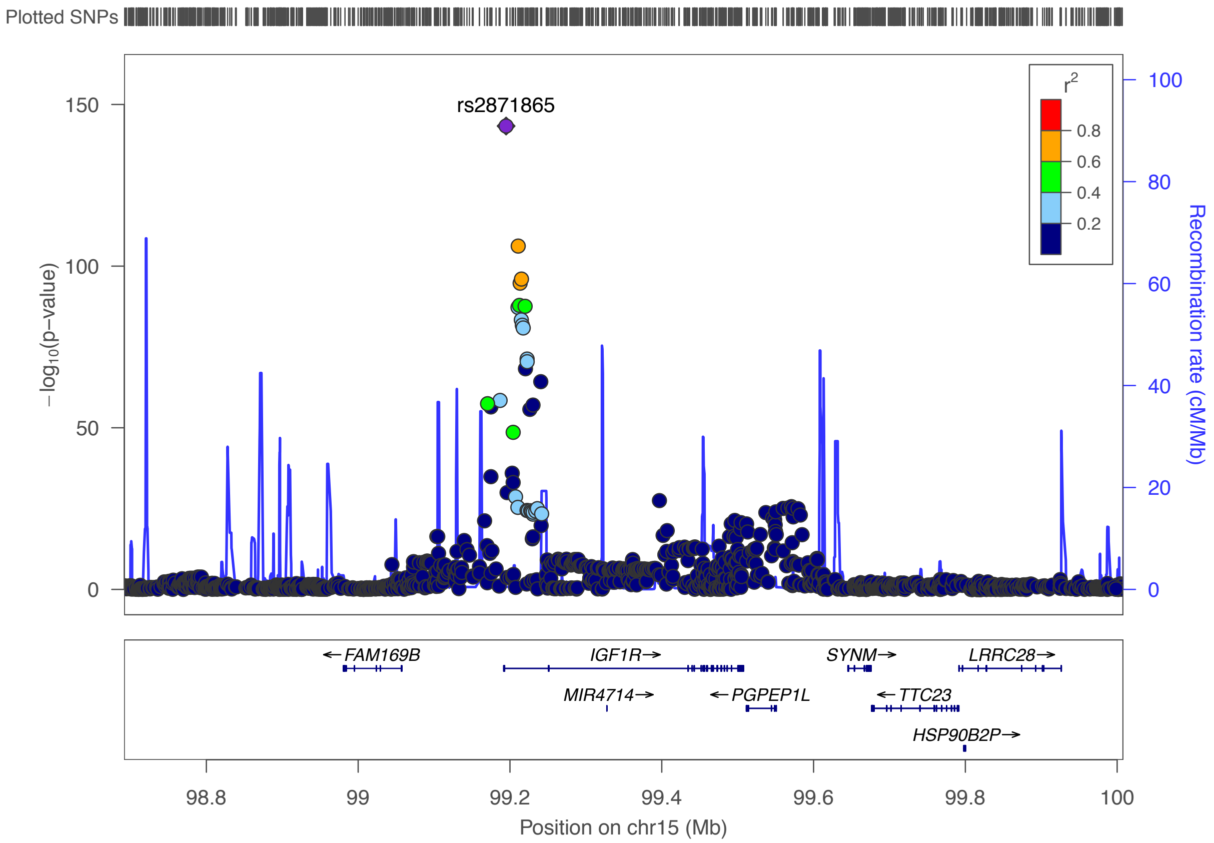

*Supplementary Figure 1. Regional plot near the IGF1R gene. The Y-axis represents the -log10 p-value for height in the Yengo et al. meta-analysis.*

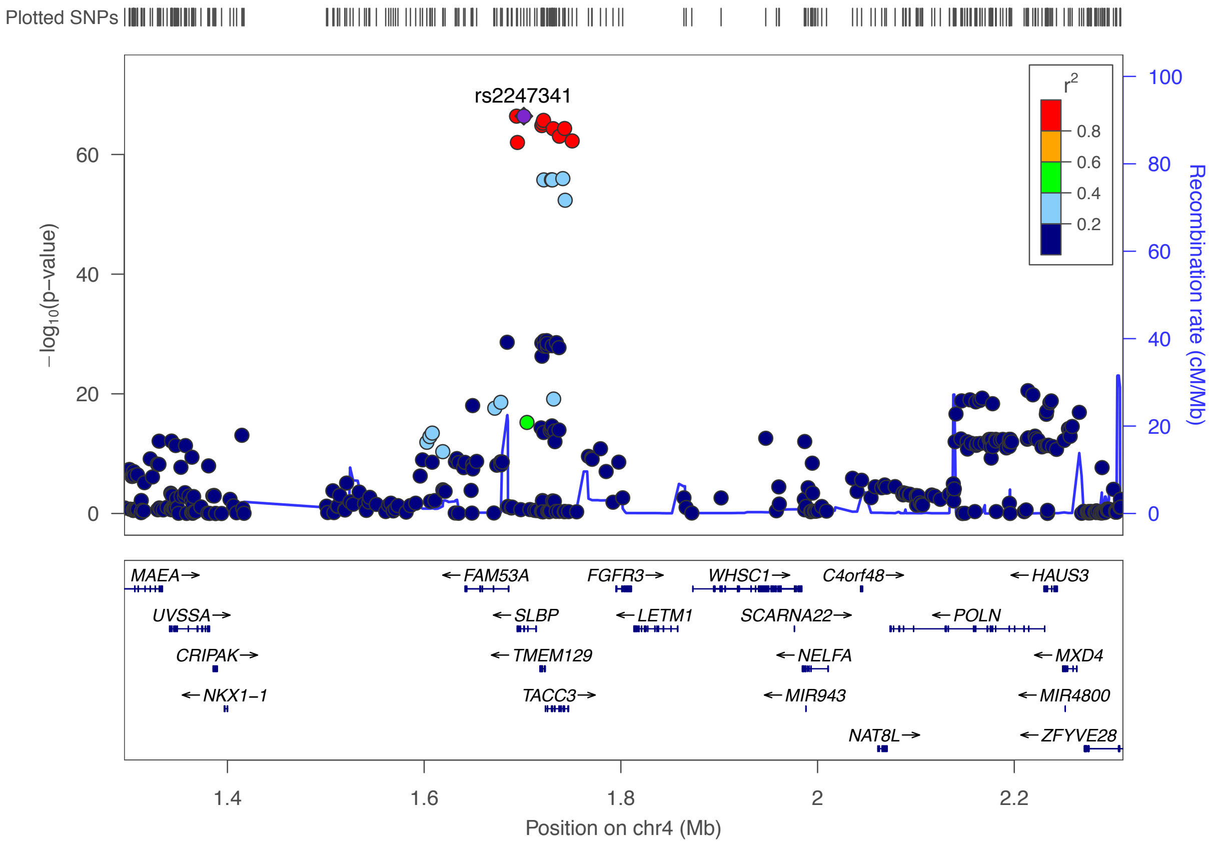

*Supplementary Figure 2. Regional plot near the FGFR3 gene. Y-axis represents the -log10 p-value for height in the Yengo et al. meta-analysis.*

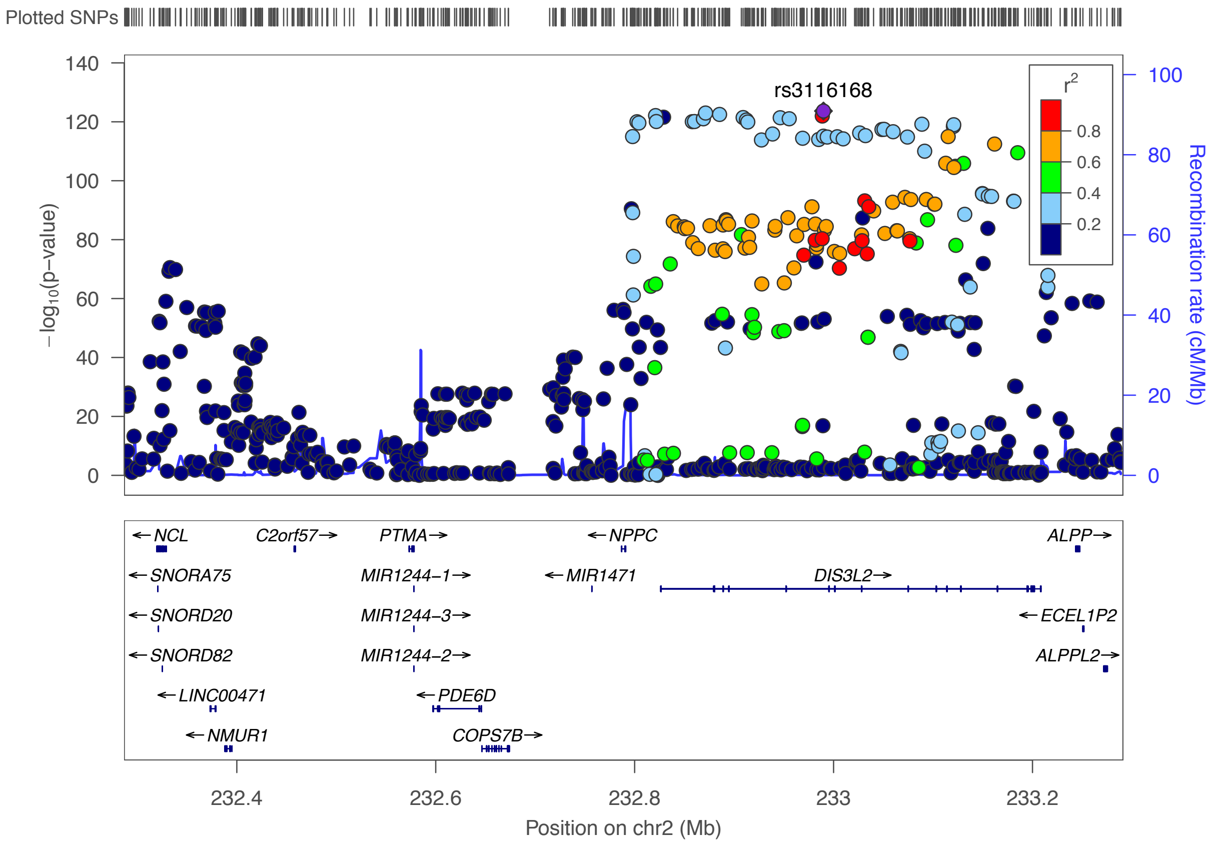

*Supplementary Figure 3. Regional plot near the NPPC gene. Y-axis represents the -log10 p-value for height in the Yengo et al. meta-analysis.*

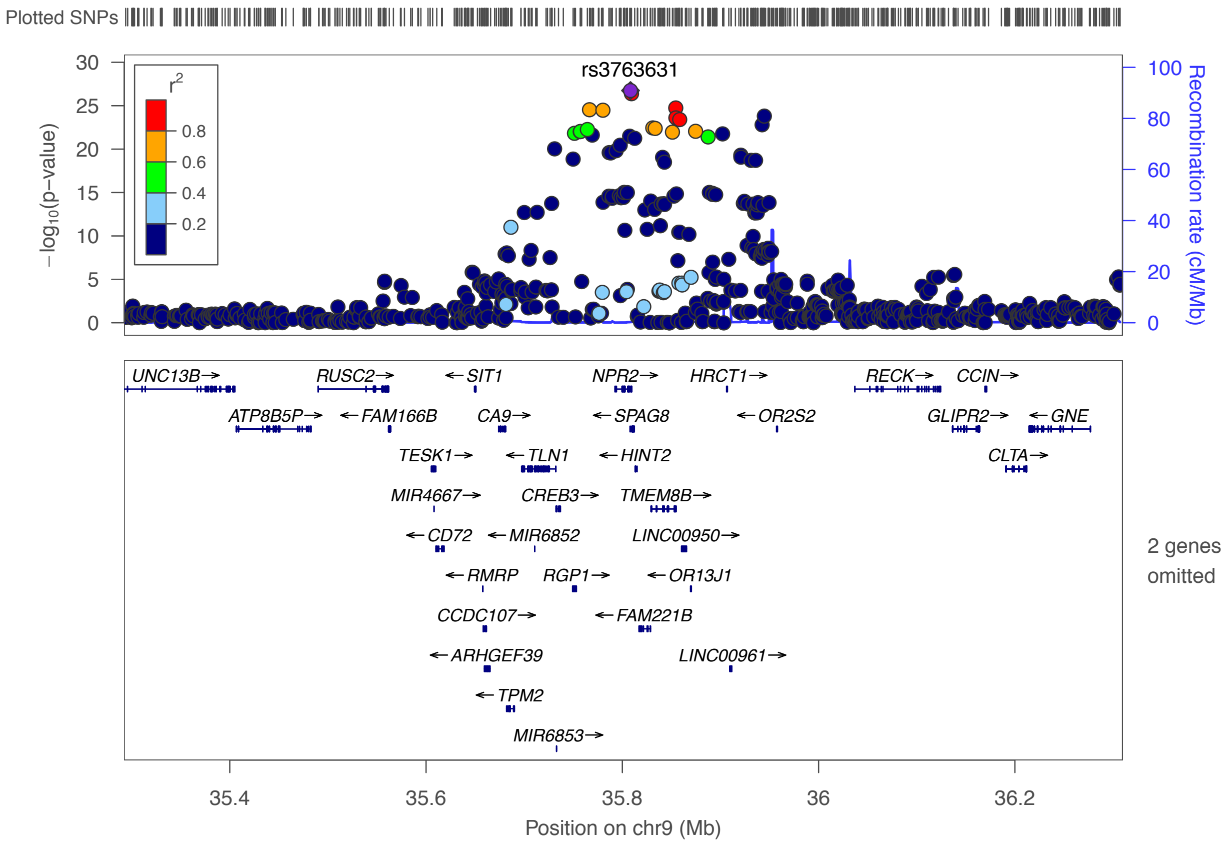

*Supplementary Figure 4. Regional plot near the NPR2 gene. Y-axis represents the -log10 p-value for height in the Yengo et al. meta-analysis.*

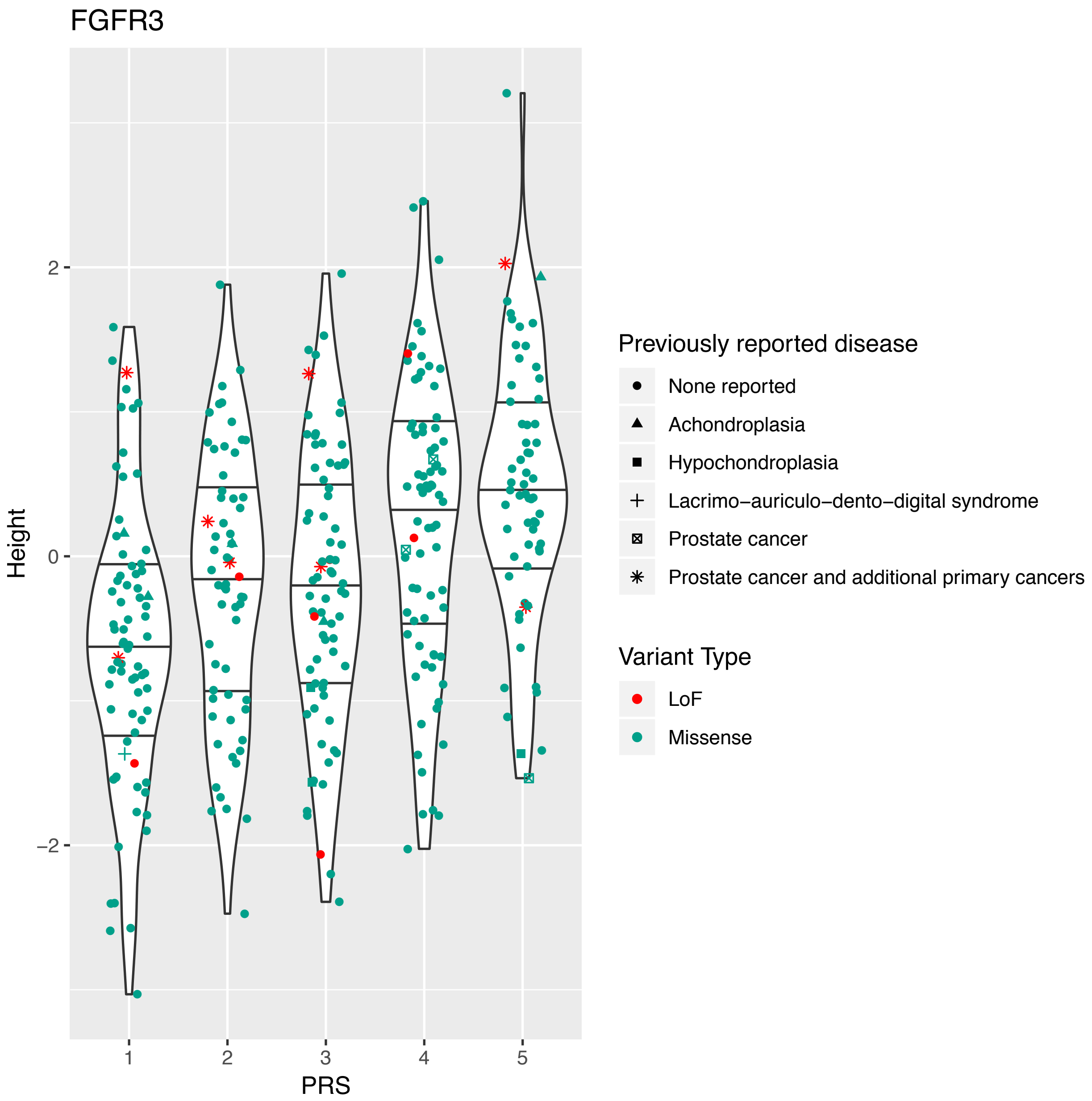

*Supplementary Figure 5. Effect of FGFR3 missense and loss of function variants on height stratified by PRS*

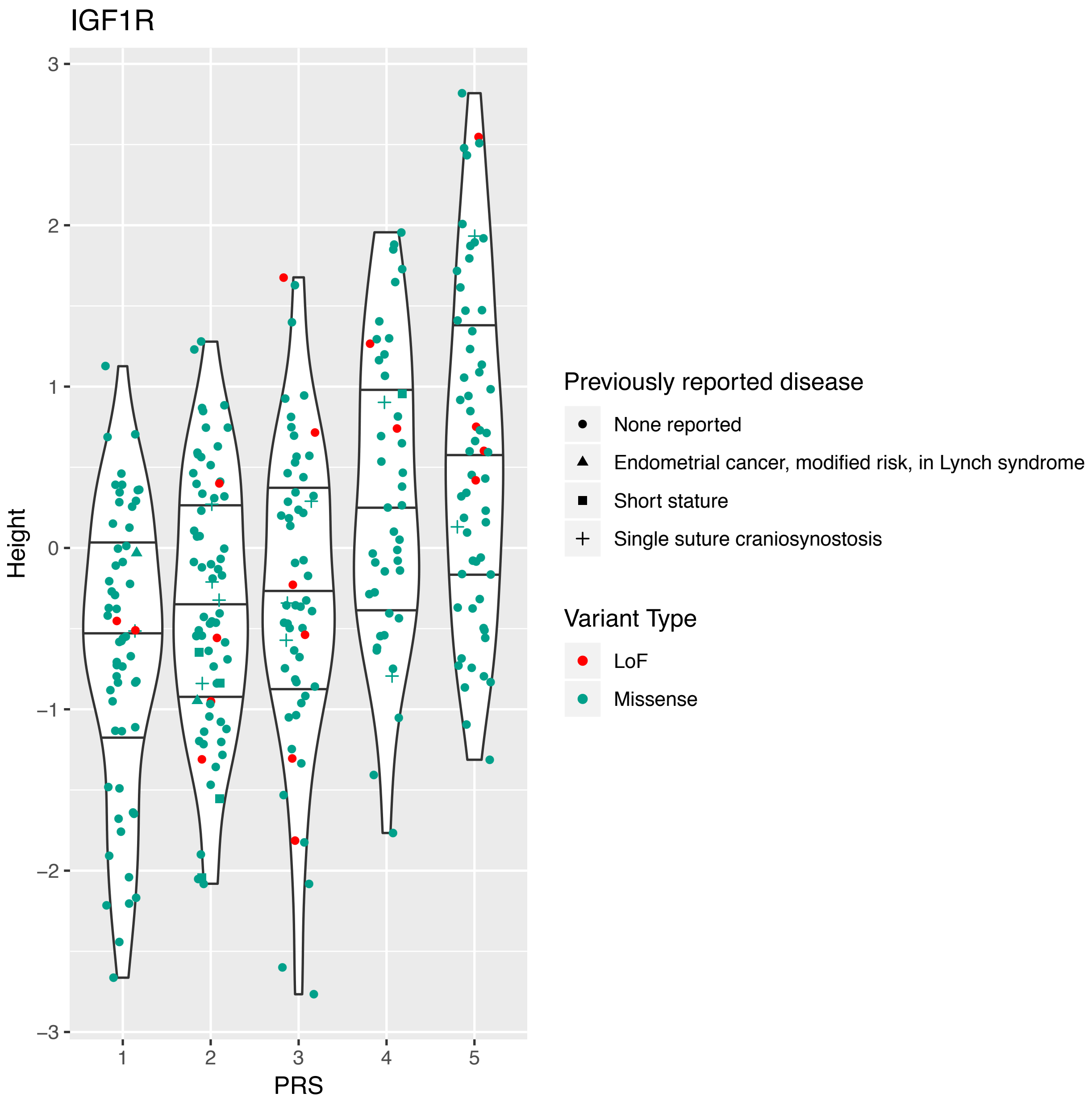

*Supplementary Figure 6. Effect of IGF1R missense and loss of function variants on height stratified by PRS.*

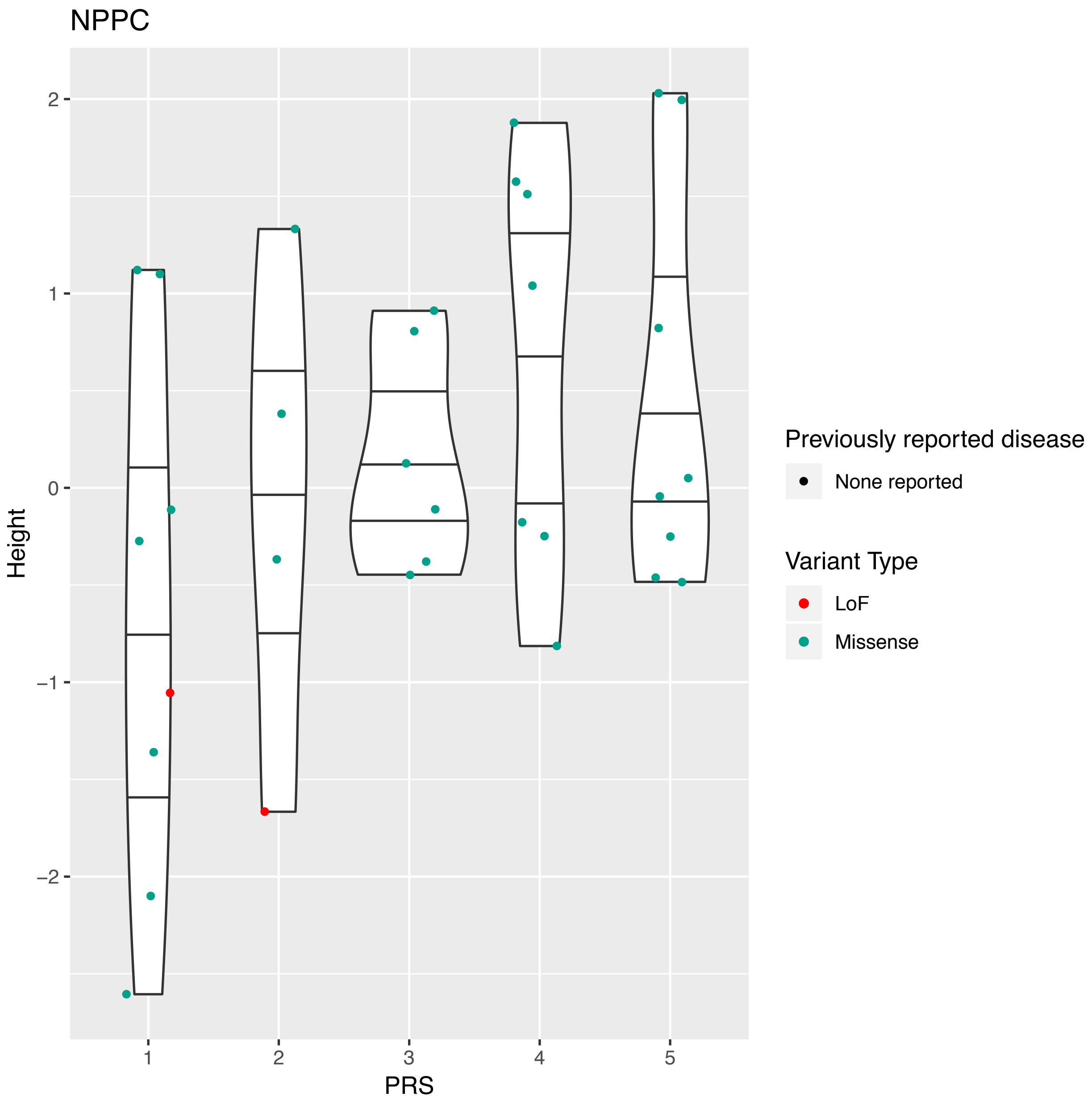

*Supplementary Figure 7. Effect of NPPC missense and loss of function variants on height stratified by PRS.*

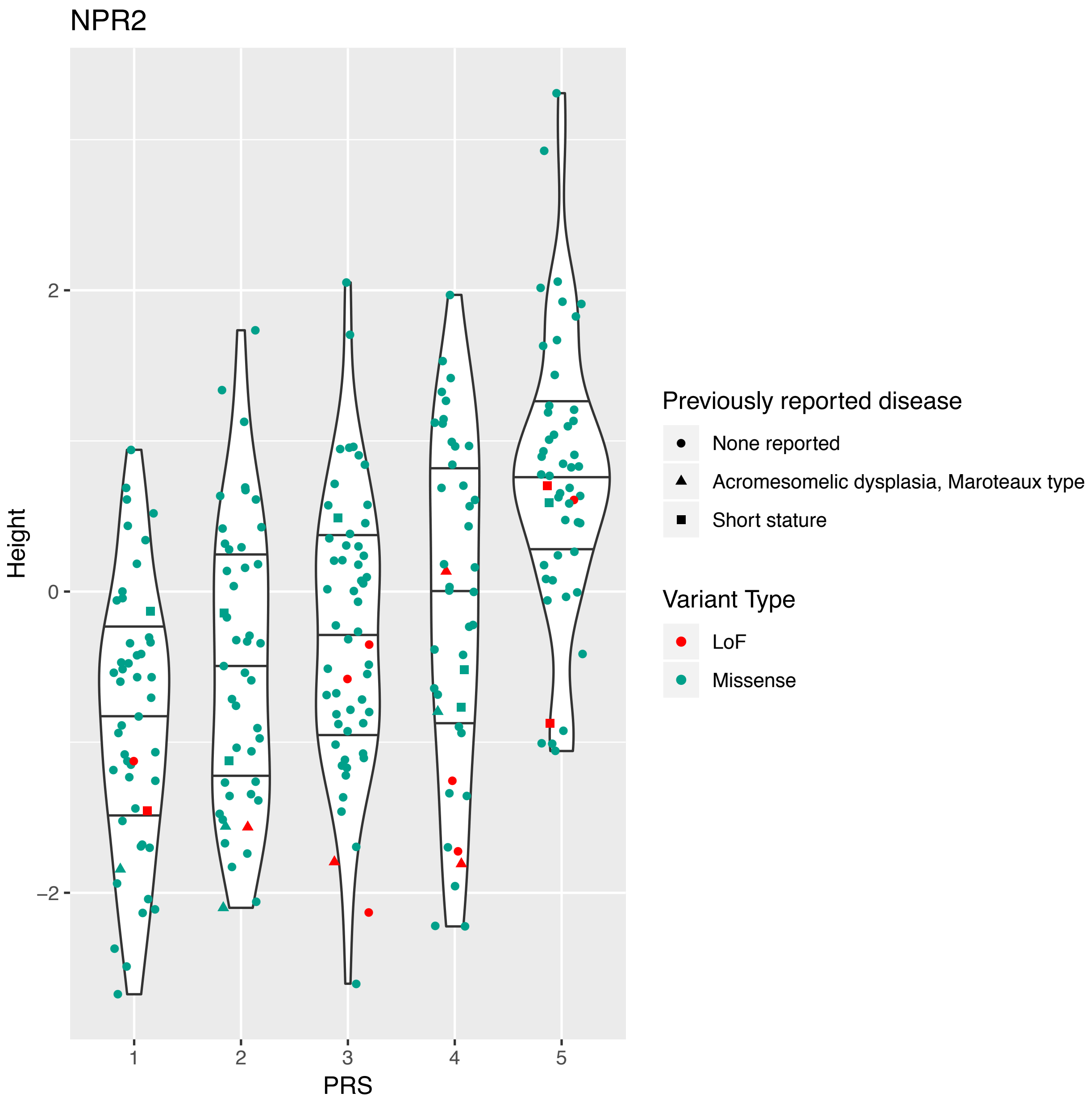

*Supplementary Figure 8. Effect of NPR2 missense and loss of function variants on height stratified by PRS.*

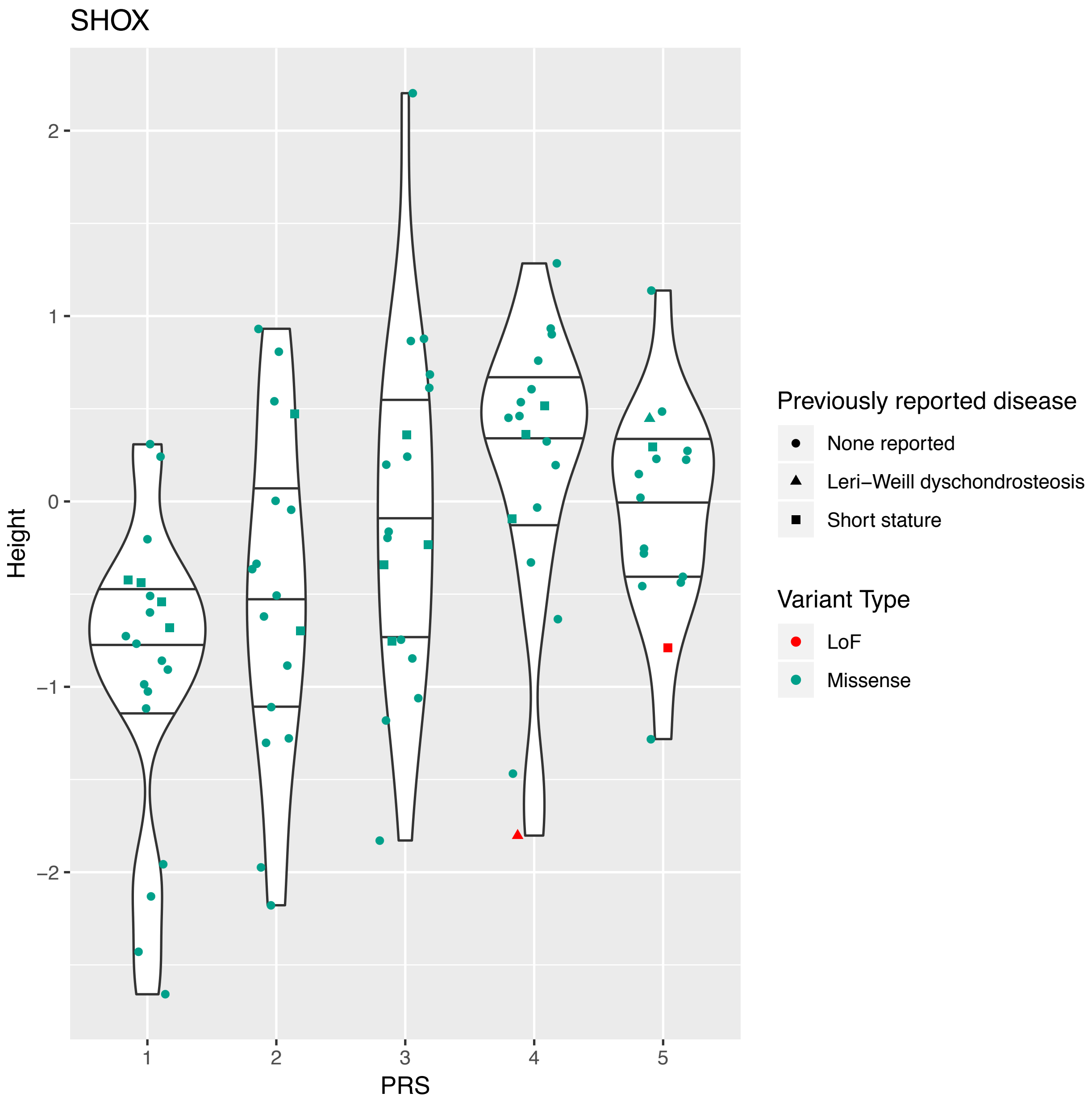

*Supplementary Figure 9. Effect of SHOX missense and loss of function variants on height stratified by PRS.*

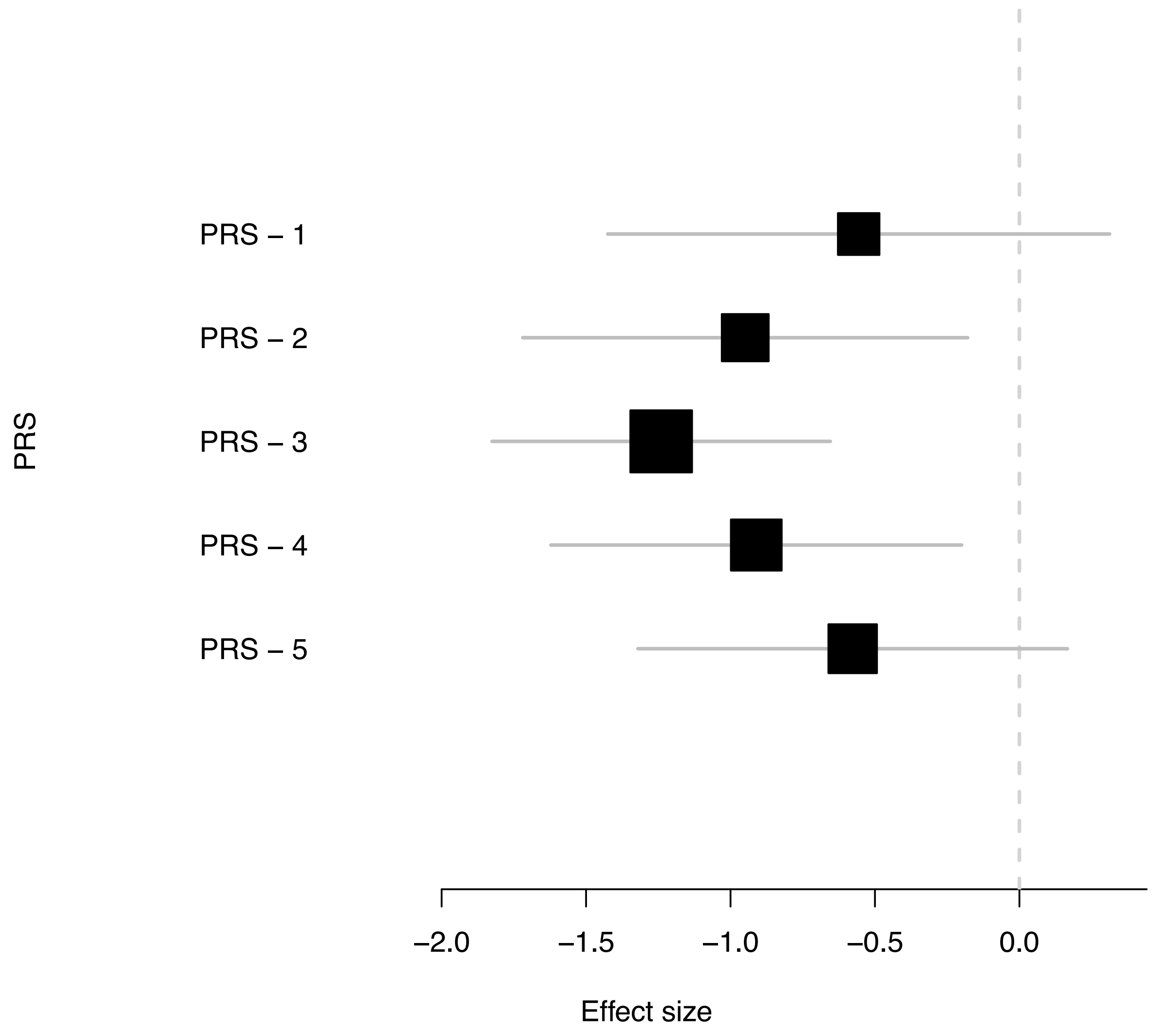

*Supplementary Figure 10. Mean effect on height of LoF status standardized by PRS quintiles*

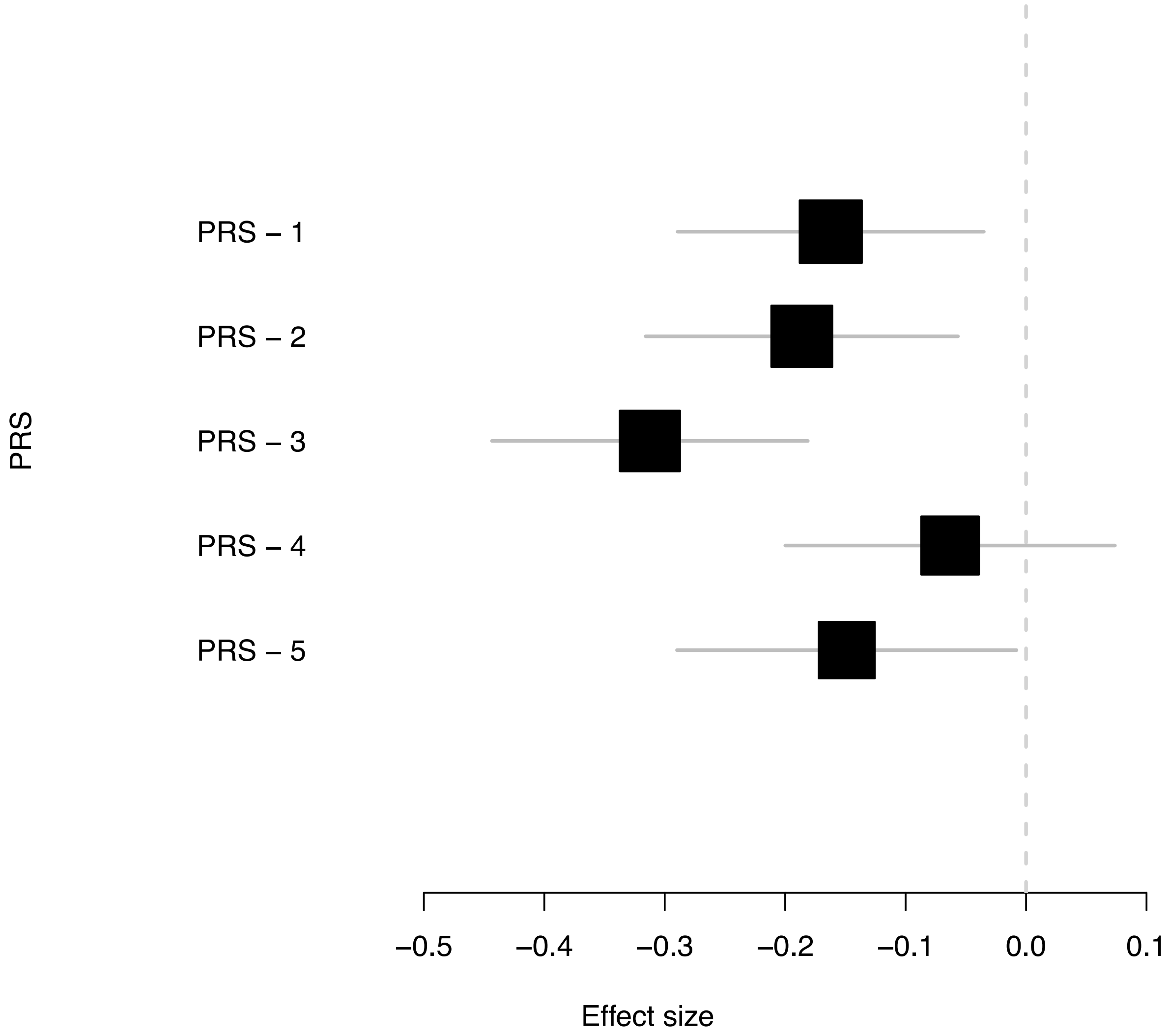

*Supplementary Figure 11. Mean effect on height of missense status standardized by PRS quintiles*

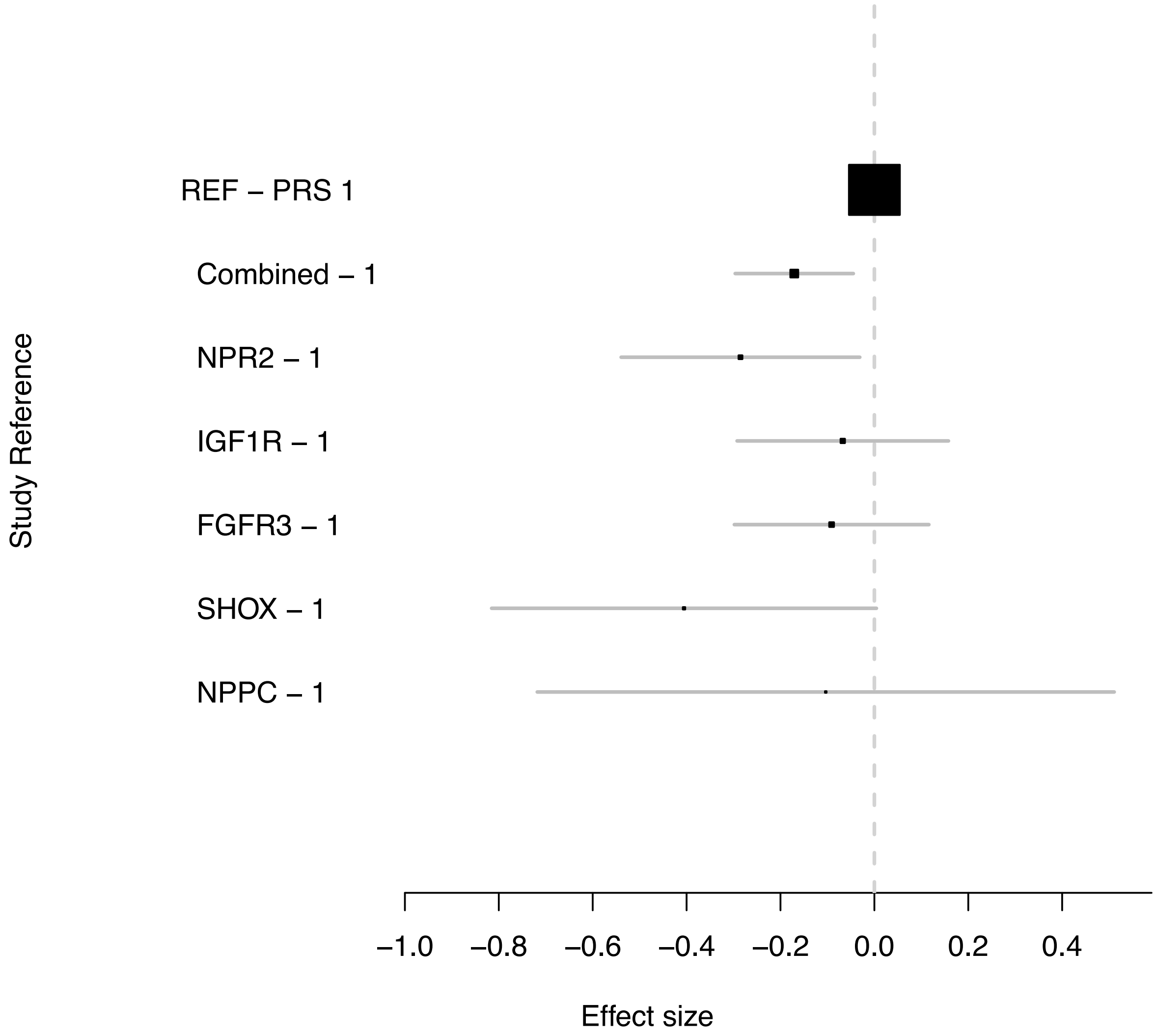

*Supplementary Figure 12. Effect of having a missense and/or Lof variant in five genes using non-carriers at PRS 1 quintile as reference*

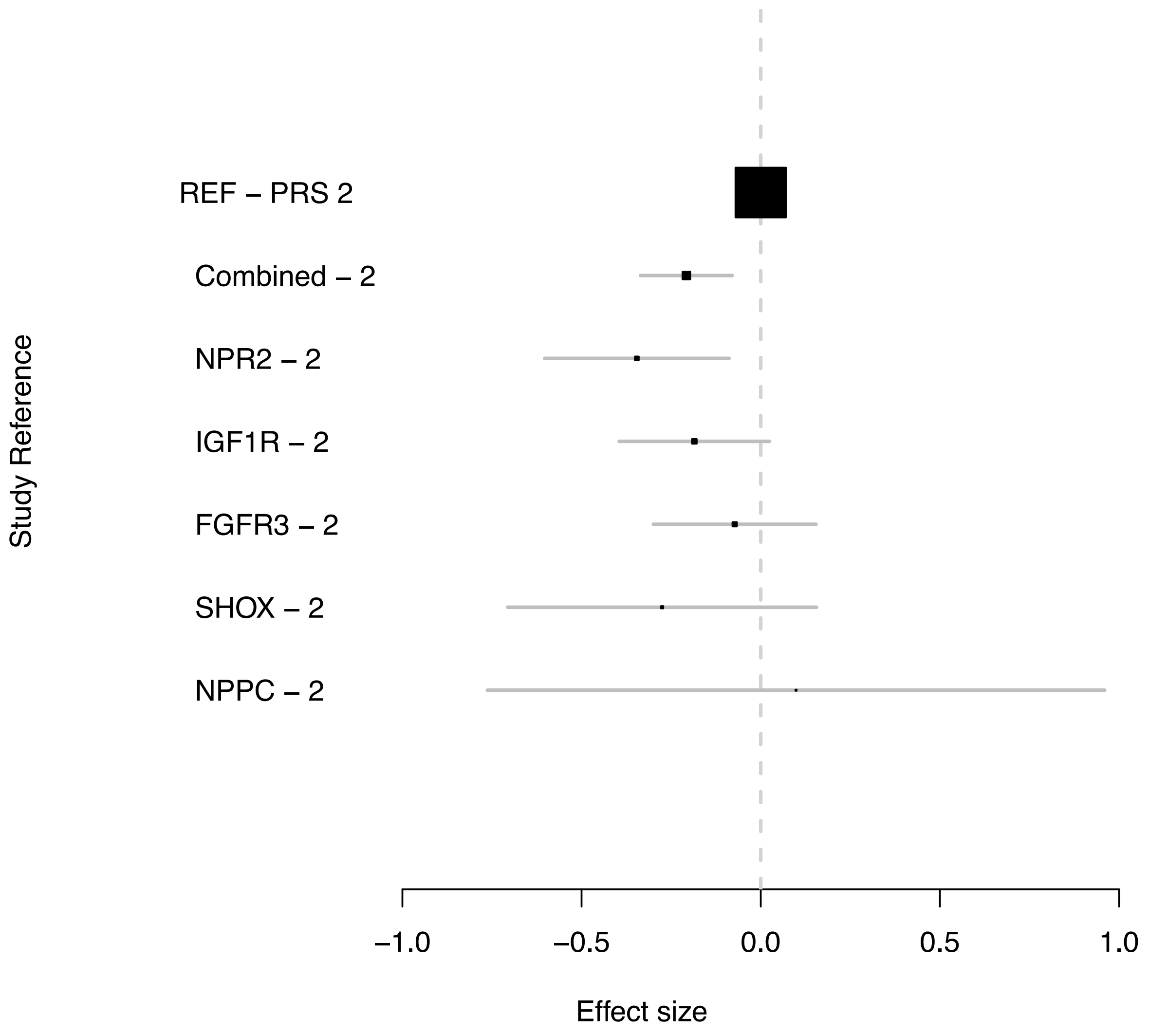

*Supplementary Figure 13. Effect of having a missense and/or Lof variant in five genes using non-carriers at PRS 2 quintile as reference*

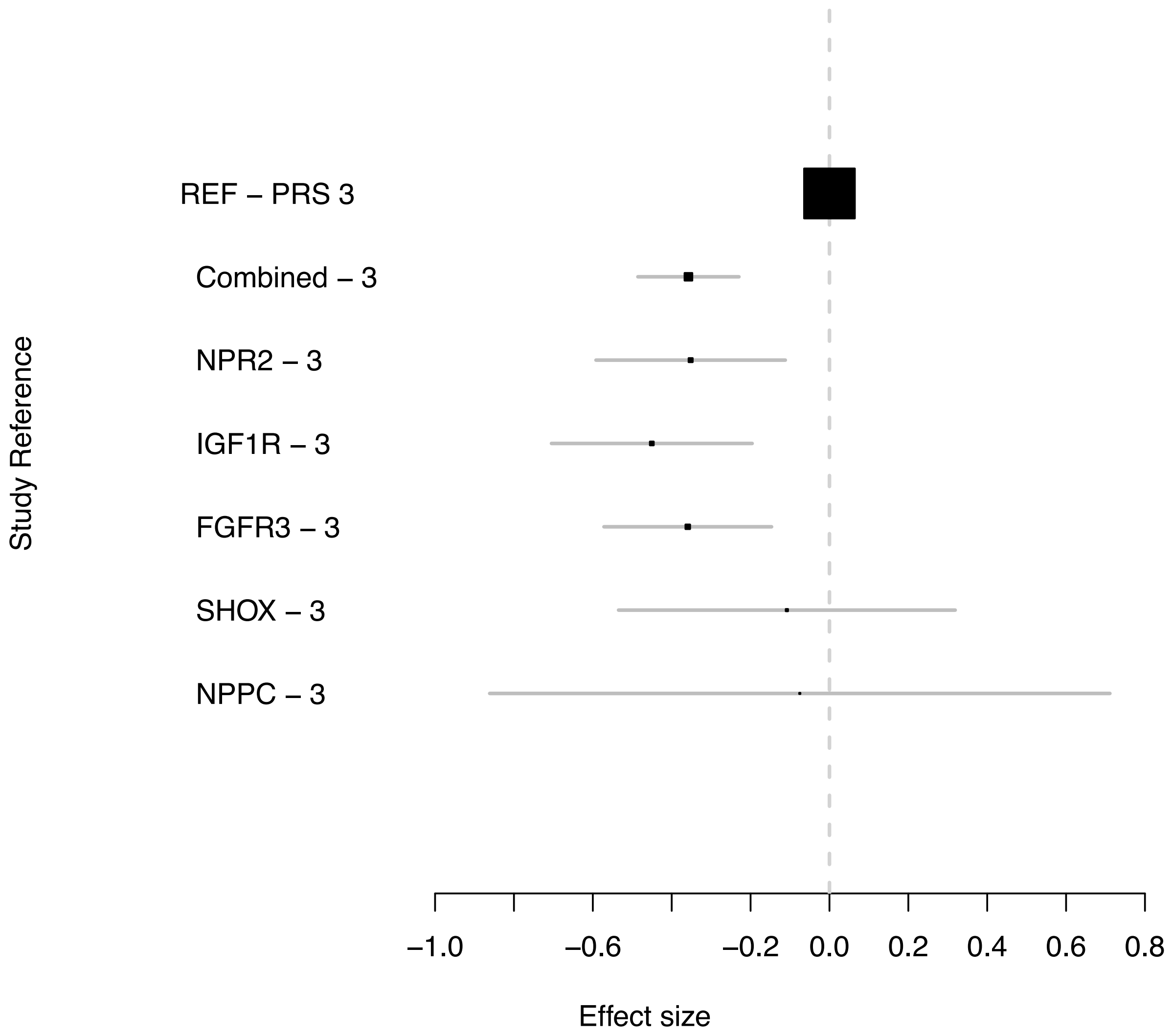

*Supplementary Figure 14. Effect of having a missense and/or Lof variant in five genes using non-carriers at PRS 3 quintile as reference*

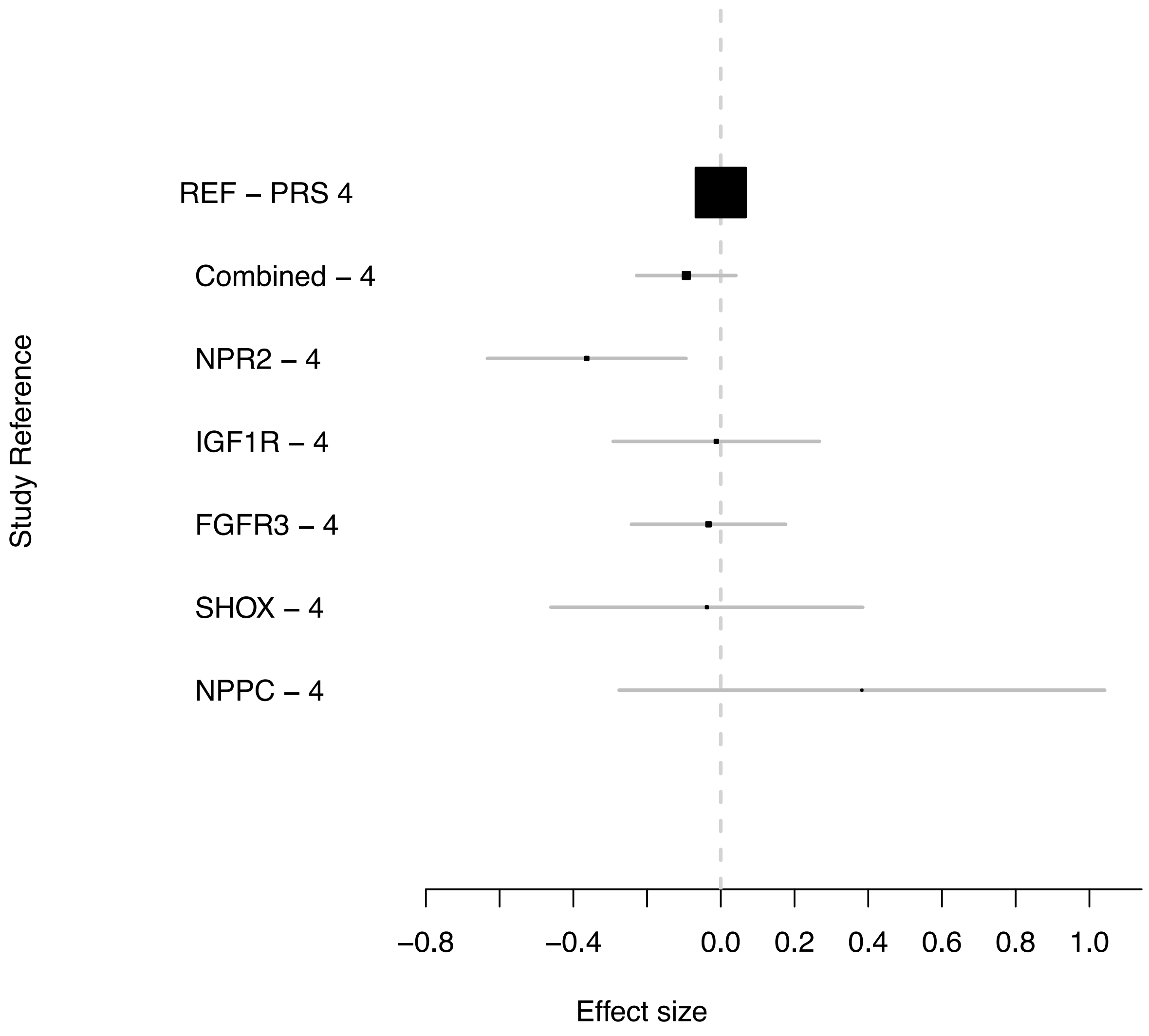

*Supplementary Figure 15. Effect of having a missense and/or Lof variant in five genes using non-carriers at PRS 4 quintile as reference*

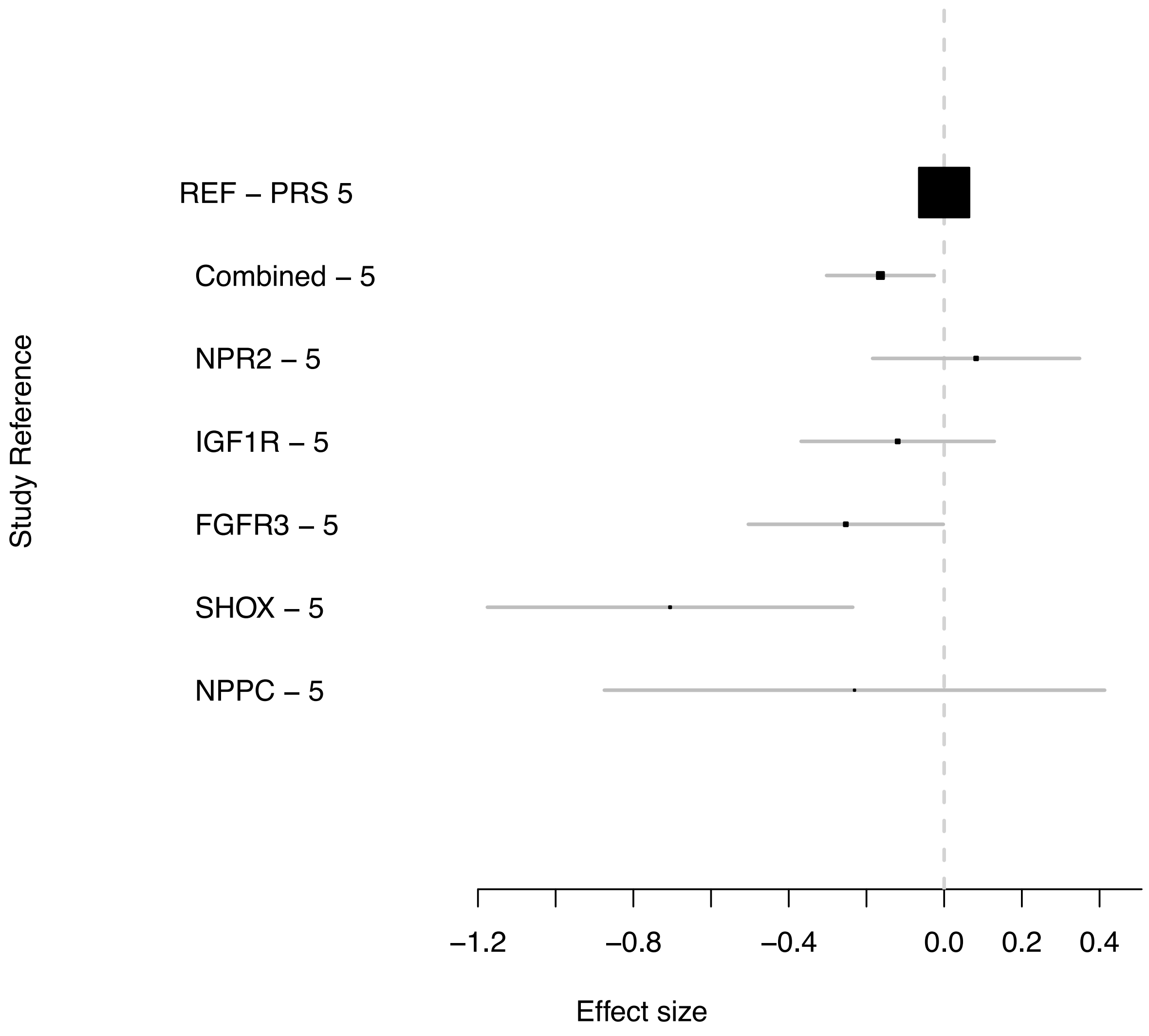

*Supplementary Figure 16. Effect of having a missense and/or Lof variant in five genes using non-carriers at PRS 5 quintile as reference*

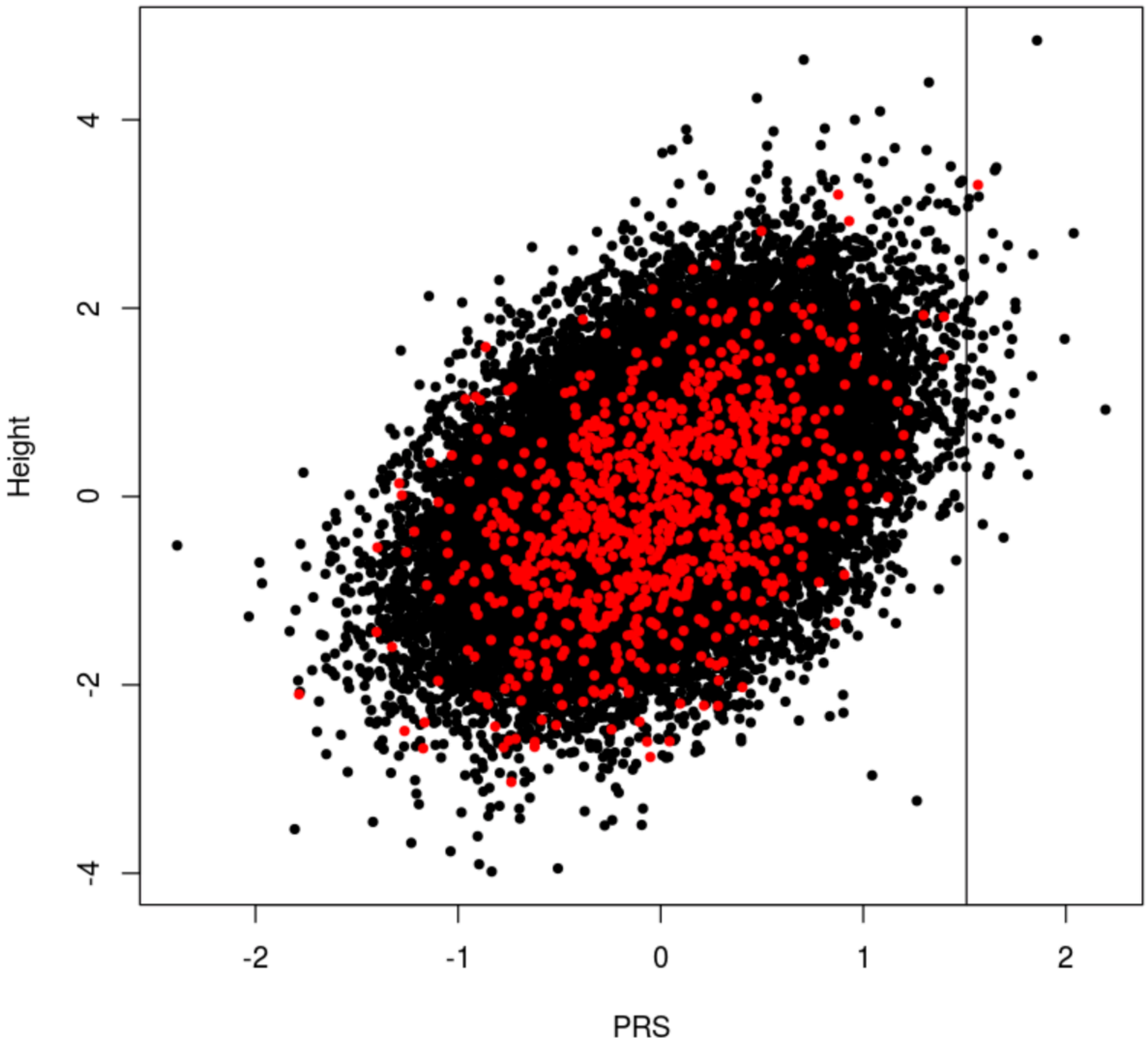

*Supplementary Figure 17. Plot of PRS vs Height in all exome sequenced individuals. Red dots represent carriers of any of the five genes. Black dots are non-carriers. Black vertical line represents 99.85% percentile of the PRS*

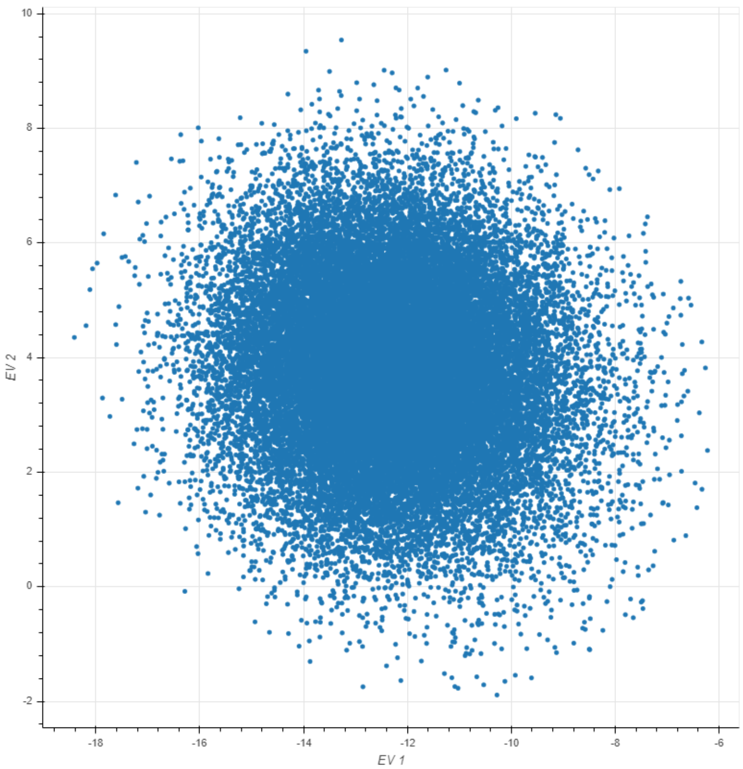

*Supplementary Figure 18. Eigenvectors 1 and 2 for 34,284 individuals with exome sequencing passing filters.*

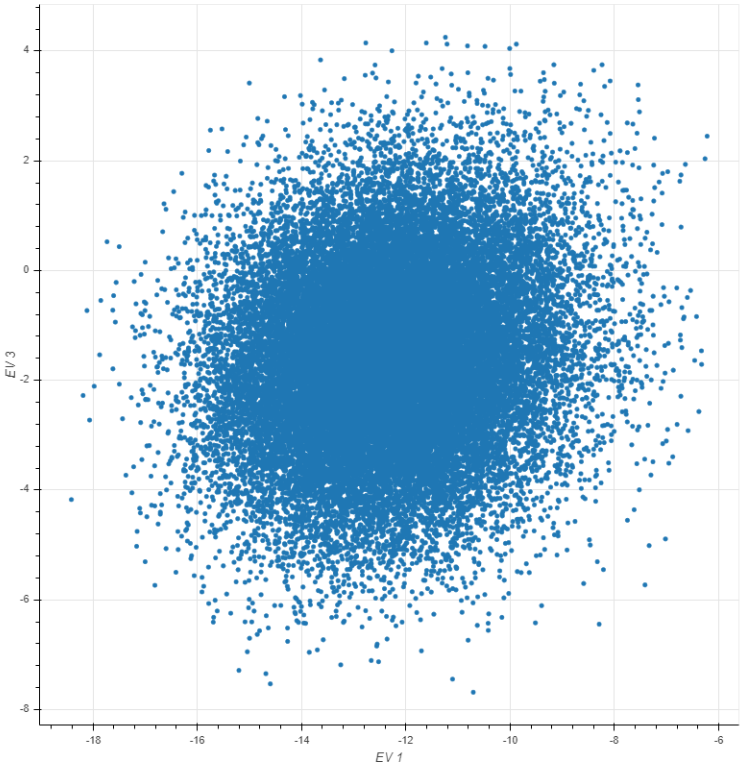

*Supplementary Figure 19. Eigenvectors 1 and 3 for 34,284 individuals with exome sequencing passing filters.*

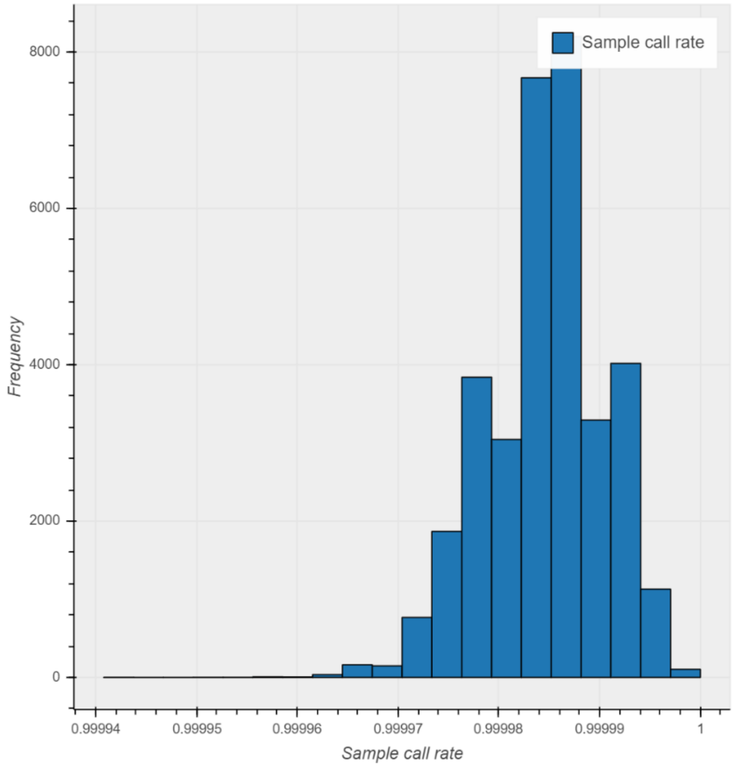

*Supplementary Figure 20. Call rate distribution for samples*

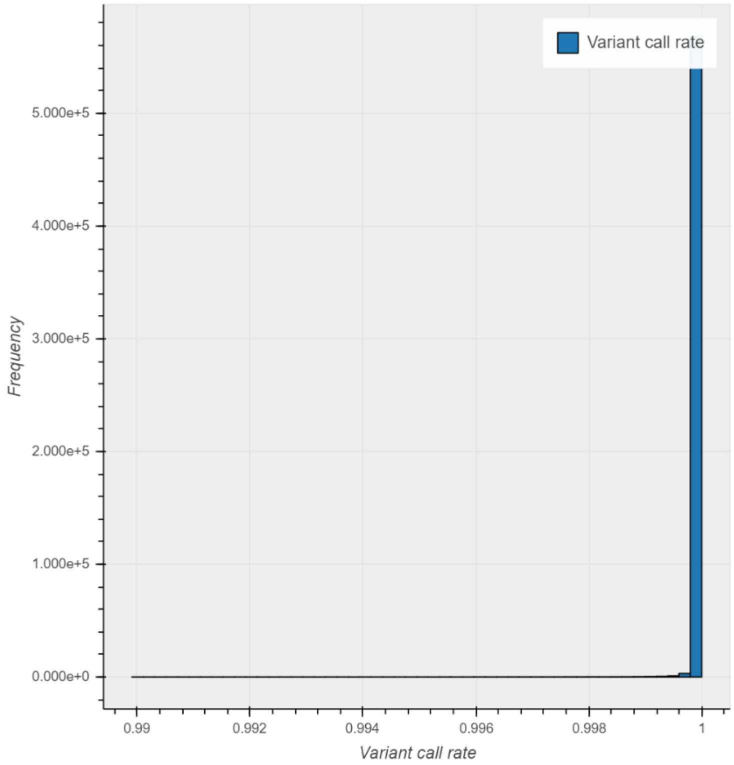

*Supplementary Figure 21. Call rate distribution for variants.*

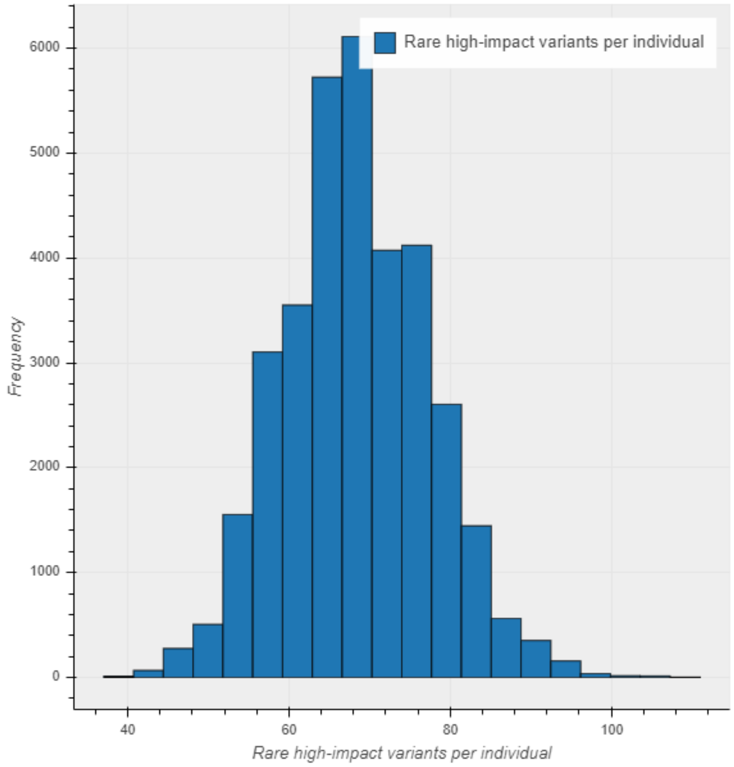

*Supplementary Figure 22. Rare high-impact variants per individual (autosomal variants only).*

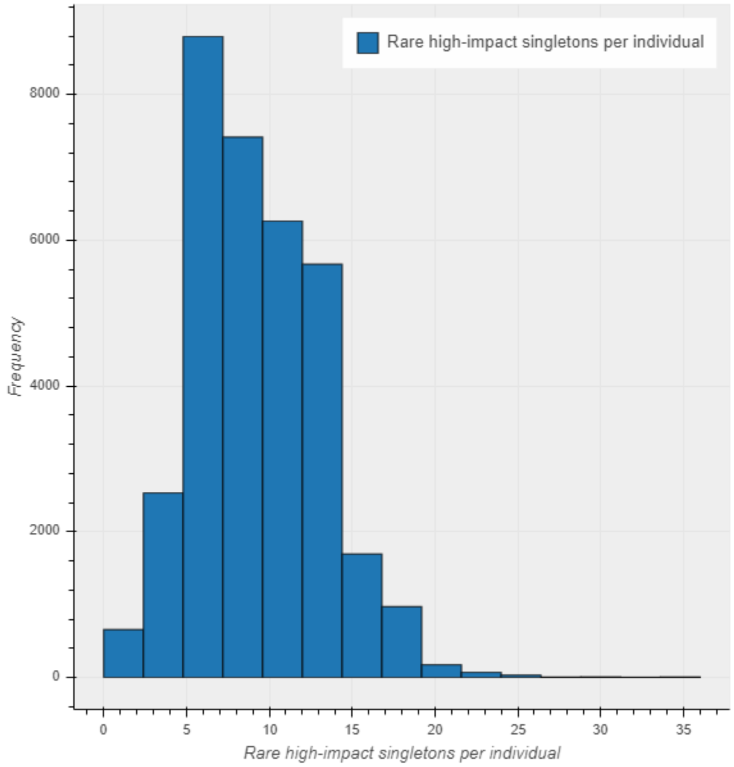

*Supplementary Figure 23. Rare high-impact singletons per individual (autosomal variants only).*

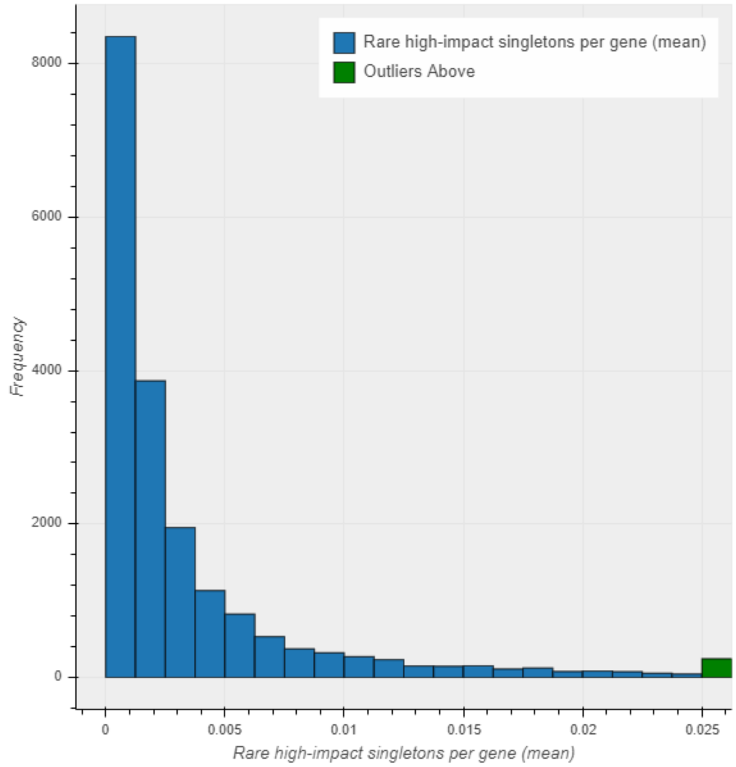

*Supplementary Figure  24. Proportion of individuals (out of 34,284) with a rare high-impact variant for each gene (n=19,117)*

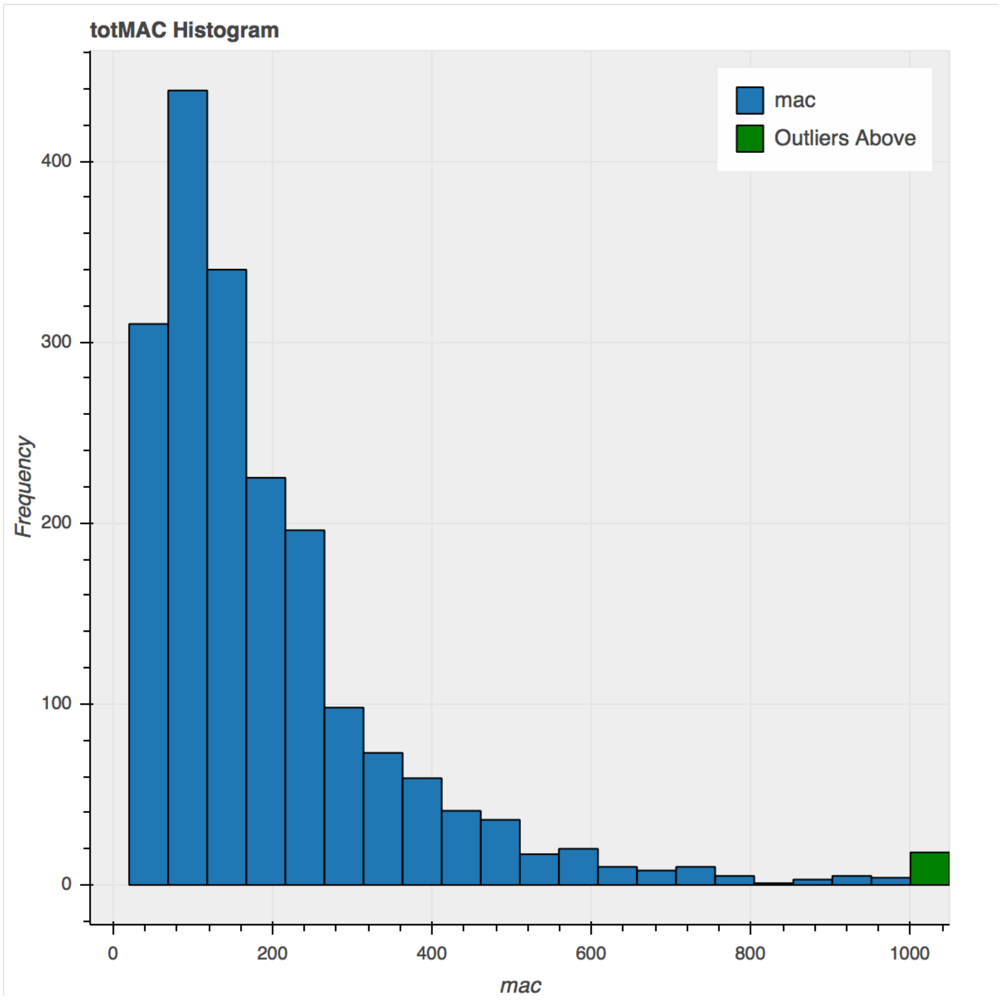

*Supplementary Figure 25. Distribution of minor allele counts in the 1918 genes included in gene set*

NPR2$^{+/+}$ clone, Allele 1:
ATGGCACTGCCATCCCTGCTACTGGTGGTGGCAGCCCTGGCAGGTGGGGTGCGTCCTCCGGGGGCACG
GAACCTGACGCTGGCGGTGGTGCTGCCAGAACACAACCTGAGCTATGCCTGGGCCTGGCCACGGGTGG
GTCCTGCTGTGGCACTGGCTGTGGAGGCGCTGGGCCGGGCACTGCCCGTGGACCTGCGGTTTGTCAGC
TCCGAACTAGACGGCGCCTGCTCTGAGTACCTGGCACCACTGCGCGCTGTGGATCTCAAGCTGTACCAT
GACCCCGACCTTCTGTTGGGCCCTGGTTGTGTGTACCCTGCTGCCTCTGTGGCTCGCTTTGCCTCGCACTG
GCACCTTCCCCTCCTGACTGCGGGTGCAGTGGCCTCTGGCTTTGCAGCTAAGAATGAGCATTATCGTACC
CTGGTTCGCACTGGCCCCTCTGCGCCCAAGCTGGGTGAGTTCGTAGTGACATTACACGGGCACTTCAATT
GGACTGCTCGGGCTGCTTTGCTGTATCTGGATGCTCGCACAGATGACCGGCCCCACTACTTCACCATCGA
GGGGGTGTTTGAGGCCCTGCAGGGCAGCAACCTCAGTGTGCAACACCAGGTGTATACCCGAGAGCCAG
GTGGCCCTGAGCAAGCCACCCACTTCATCAGAGCCAACGGGCGCA

NPR2$^{+/-}$ clone, Allele 1:
ATGGCACTGCCATCCCTGCTACTGGTGGTGGCAGCCCTGGCAGGTGGGGTGCGTCCTCCGGGGGCACG
GAACCTGACGCTGGCGGTGGTGCTGCCAGAACACAACCTGAGCTATGCCTGGGCCTGGCCACGGGTGG
GTCCTGCTGTGGCACTGGCTGTGGAGGCGCTGGGCCGGGCACTGCCCGTGGACCTGCGGTTTGTCAGC
TCCGAACTAGACGGCGCCTGCTCTGAGTACCTGGCACCACTGCGCGCTGTGGATCTCAAGCTGTACCAT
GACCCCGACCTTCTGTTGGGCCCTGGTTGTGTGTACCCTGCTGCCTCTGTGGCTCGCTTTGCCTCGCACTG
GCACCTTCCCCTCCTGACTGCGGGTGCAGTGGCCTCTGGCTTTGCAGCTAAGAATGAGCATTATCGTACC
CTGGTTCGCACTGGCCCCTCTGCGCCCAAGCTGGGTGAGTTCGTAGTGACATTACACGGGCACTTCAATT
GGACTGCTCGGGCTGCTTTGCTGTATCTGGATGCTCGCACAGATGACCGGCCCCACTACTTCACCATCGA
GGGGGTGTTTGAGGCCCTGCAGGGCAGCAACCTCAGTGTGCAACACCAGGTGTATACCCGAGAGCCAG
GTGGCCCTGAGCAAGCCACCCACTTCATCAGAGCCAACGGGCGCA

NPR2$^{+/-}$ clone, Allele 2:
ATGGCACTGCCATCCCTGCTACTGGTGGTGGCAGCCCTGGCAGGTGGGGTGCGTCCTCCGGGGGCACG
GAACCTGACGCTGGCGGTGGTGCTGCCAGAACACAACCTGAGCTATGCCTGGGCCTGGCCACGGGTGG
GTCCTGCTGTGGCACTGGCTGTGGAGGCGCTGGGCCGGGCACTGCCCGTGGACCTGCGGTTTGTCAGC
TCCGAACTAGACGGCGCCTGCTCTGAGTACCTGGCACCACTGCGCGCTGTGGATCTCAAGCTGTACCAT
GACCCCGACCTTCTGTTGGGCCCTGGTTGTGTGTACCCTGCTGCCTCTGTGGCTCGCTTTGCCTCGCACTG
GCACCTTCCCCTCCTGACTGCGGGTGCAGTGGCCTCTGGCTTTGCAGCTAAGAATGAGCATTATCGTACC
CTGGTTCGCACTGGCCCCTCTGCGCCCAAGCTGGGTGAGTTCGTAGTGACATTACACGGGCACTTCACA
ATTGGACTGCTCGGGCTGCTTTGCTGTATCTGGATGCTCGCACAGATGACCGGCCCCACTACTTCACCAT
CGAGGGGGTGTTTGAGGCCCTGCAGGGCAGCAACCTCAGTGTGCAACACCAGGTGTATACCCGAGAGC
CAGGTGGCCCTGAGCAAGCCACCCACTTCATCAGAGCCAACGGGCGCA

NPR2$^{-/-}$ clone, Allele 1:
ATGGCACTGCCATCCCTGCTACTGGTGGTGGCAGCCCTGGCAGGTGGGGTGCGTCCTCCGGGGGCACG
GAACCTGACGCTGGCGGTGGTGCTGCCAGAACACAACCTGAGCTATGCCTGGGCCTGGCCACGGGTGG
GTCCTGCTGTGGCACTGGCTGTGGAGGCGCTGGGCCGGGCACTGCCCGTGGACCTGCGGTTTGTCAGC
TCCGAACTAGACGGCGCCTGCTCTGAGTACCTGGCACCACTGCGCGCTGTGGATCTCAAGCTGTACCAT
GACCCCGACCTTCTGTTGGGCCCTGGTTGTGTGTACCCTGCTGCCTCTGTGGCTCGCTTTGCCTCGCACTG
GCACCTTCCCCTCCTGACTGCGGGTGCAGTGGCCTCTGGCTTTGCAGCTAAGAATGAGCATTATCGTACC

CTGGTTCGCACTGGCCCCTCTGCGCCCAAGCTGGGTGAGTTCGTAGTGACATTACACGGGCACTTCAATT
GGACTGCTCGGGCTGCTTTGCTGTATCTGGATGCTCGCACAGATGACCGGCCCCACTACTTCACCATC**<u>C</u>**G
AGGGGGTGTTTGAGGCCCTGCAGGGCAGCAACCTCAGTGTGCAACACCAGGTGTATACCCGAGAGCCA
GGTGGCCCTGAGCAAGCCACCCACTTCATCAGAGCCAACGGGCGCA

NPR2$^{-/-}$ clone, Allele 2:
ATGGCACTGCCATCCCTGCTACTGGTGGTGGCAGCCCTGGCAGGTGGGGTGCGTCCTCCGGGGGCACG
GAACCTGACGCTGGCGGTGGTGCTGCCAGAACACAACCTGAGCTATGCCTGGGCCTGGCCACGGGTGG
GTCCTGCTGTGGCACTGGCTGTGGAGGCGCTGGGCCGGGCACTGCCCGTGGACCTGCGGTTTGTCAGC
TCCGAACTAGACGGCGCCTGCTCTGAGTACCTGGCACCGGGGTGTTTGAGGCCCTGCAGGGCAGCAAC
CTCAGTGTGCAACACCAGGTGTATACCCGAGAGCCAGGTGGCCCTGAGCAAGCCACCCACTTCATCAGA
GCCAACGGGCGCA

*Supplementary Figure 26. Sequence of exon 1 of the NPR2 gene in CRISPR edited clones. NPR2$^{+/+}$ clone has one allele matching that of wild-type RCS cells. In the NPR2$^{+/-}$ clone allele 1 matches that of wild-type cells and allele 2 has a 2bp insertion. The NPR2$^{-/-}$ clone has a 1bp insertion and allele 2 has a 313bp deletion.*

#### Supplementary Tables

Supplementary Table 1. List of lead SNPs from GWAS meta-analysis of height (Online)

Supplementary Table 2. List of height genes and their respective annotations (Online)

Supplementary Table 3 Study Descriptives for UK Biobank samples with exome-sequence data

| Phenotype | Gender | n | Min. | 1st Q | Median | Mean | 3rd Q | Max. | ISS | % ISS |
|---|---|---|---|---|---|---|---|---|---|---|
| | Women | 18417 | 137 | 159 | 163 | 163 | 167 | 193 | 309 | 1.68% |
| Height | Men | 15814 | 148 | 172 | 176 | 176.3 | 181 | 203 | 263 | 1.66% |
| | Women | 18417 | 40 | 50 | 58 | 56.5 | 63 | 70 | | |
| Age | Men | 15814 | 40 | 52 | 59 | 57.59 | 64 | 70 | | |

Supplementary Table 4 SKAT Association results for the 19 gene sets tested on height

| Gene set id | labels | Miss+LoF | Miss | LoF |
|---|---|---|---|---|
| 1 | [1, 1, 1, 1, 1] | 6.67E-07 | 2.85E-05 | 1.27E-05 |
| 2 | [0, 1, 1, 1, 1] | 1.76E-05 | 5.12E-05 | 0.05 |
| 3 | [0, 1, 0, 0, 1] | 0.01 | 0.01 | 0.41 |
| 4 | [1, 1, 0, 0, 0] | 0.03 | 0.10 | 0.01 |
| 5 | [0, 0, 1, 0, 0] | 0.03 | 0.05 | 0.33 |
| 6 | [0, 1, 0, 0, 0] | 0.05 | 0.03 | 0.70 |
| 7 | [1, 0, 0, 1, 0] | 0.08 | 0.06 | 0.48 |
| 8 | [1, 0, 0, 0, 0] | 0.13 | 0.05 | 0.06 |
| 9 | [1, 0, 1, 1, 0] | 0.27 | 0.61 | 0.03 |
| 10 | [1, 1, 0, 1, 0] | 0.28 | 0.28 | NA |
| 11 | [1, 0, 1, 0, 0] | 0.42 | 0.44 | 0.80 |
| 12 | [1, 0, 0, 0, 1] | 0.44 | 0.46 | 0.84 |
| 13 | [0, 0, 0, 1, 1] | 0.53 | 0.53 | 1.00 |
| 14 | [0, 0, 0, 0, 0] | 0.68 | 0.85 | 0.37 |
| 15 | [1, 1, 0, 0, 1] | 0.69 | 0.88 | 0.00 |
| 16 | [0, 0, 0, 0, 1] | 0.70 | NA | 0.13 |
| 17 | [0, 0, 0, 1, 0] | 0.75 | 0.89 | 0.34 |
| 18 | [1, 0, 1, 1, 1] | 0.85 | 0.73 | 0.07 |
| 19 | [1, 0, 0, 1, 1] | 0.89 | 0.92 | 0.73 |

Labels represent wether gene is present in annotation (1=present, 0= not present)
Label Order: GWAS, HGMD_Short, HGMD_Tall, OMIM_overgrowth, OMIM_short
Bonferroni significance threashold: 0.003
Only gene sets with at least one gene are reported.

Supplementary Table 5 Genetic variants found in five height genes

| chr | position | alleles | AC | nt_qc.call_ | vep.consequence | vep.impact | vep.symbol | vep.hgvsc | vep.hgvsp | vep.gnom | vep.cadd_phred |
|---|---|---|---|---|---|---|---|---|---|---|---|
| 2 | 231925451 | ['C', 'G'] | 1 | 1.00 | ['missense_variant'] | MODERATE | NPPC | ENST00000409852.2:c.355G>C | ENSP00000387159.1:p.Gly119Arg | | 33 |
| 2 | 231925486 | ['G', 'GT'] | 1 | 1.00 | ['frameshift_variant'] | HIGH | NPPC | ENST00000409852.2:c.319_320insA | ENSP00000387159.1:p.Ser107TyrfsTer37 | | |
| 2 | 231925513 | ['T', 'C'] | 1 | 1.00 | ['missense_variant'] | MODERATE | NPPC | ENST00000409852.2:c.293A>G | ENSP00000387159.1:p.Tyr98Cys | | 23 |
| 2 | 231925514 | ['A', 'C'] | 2 | 1.00 | ['missense_variant'] | MODERATE | NPPC | ENST00000409852.2:c.292T>G | ENSP00000387159.1:p.Tyr98Asp | | 18 |
| 2 | 231925523 | ['C', 'A'] | 2 | 1.00 | ['missense_variant'] | MODERATE | NPPC | ENST00000409852.2:c.283G>T | ENSP00000387159.1:p.Ala95Ser | | 15 |
| 2 | 231925529 | ['G', 'A'] | 1 | 1.00 | ['missense_variant'] | MODERATE | NPPC | ENST00000409852.2:c.277C>T | ENSP00000387159.1:p.Pro93Ser | 4.04E-06 | 26 |
| 2 | 231925540 | ['A', 'T'] | 1 | 1.00 | ['missense_variant'] | MODERATE | NPPC | ENST00000409852.2:c.266T>A | ENSP00000387159.1:p.Leu89Gln | | 23 |
| 2 | 231925550 | ['C', 'G'] | 1 | 1.00 | ['missense_variant'] | MODERATE | NPPC | ENST00000409852.2:c.256G>C | ENSP00000387159.1:p.Ala86Pro | | 23 |
| 2 | 231925553 | ['A', 'G'] | 1 | 1.00 | ['missense_variant'] | MODERATE | NPPC | ENST00000409852.2:c.253T>C | ENSP00000387159.1:p.Trp85Arg | 4.06E-06 | 23 |
| 2 | 231925580 | ['G', 'T'] | 3 | 1.00 | ['missense_variant'] | MODERATE | NPPC | ENST00000409852.2:c.226C>A | ENSP00000387159.1:p.Arg76Ser | 6.96E-05 | 25 |
| 2 | 231925584 | ['G', 'C'] | 1 | 1.00 | ['missense_variant'] | MODERATE | NPPC | ENST00000409852.2:c.222C>G | ENSP00000387159.1:p.Asp74Glu | | 21 |
| 2 | 231925597 | ['C', 'A'] | 1 | 1.00 | ['missense_variant'] | MODERATE | NPPC | ENST00000409852.2:c.209G>T | ENSP00000387159.1:p.Arg70Leu | | 26 |
| 2 | 231925600 | ['G', 'T'] | 1 | 1.00 | ['stop_gained'] | HIGH | NPPC | ENST00000409852.2:c.206C>A | ENSP00000387159.1:p.Ser69Ter | | 38 |
| 2 | 231925639 | ['G', 'A'] | 7 | 1.00 | ['missense_variant'] | MODERATE | NPPC | ENST00000409852.2:c.167C>T | ENSP00000387159.1:p.Ala56Val | 3.95E-05 | 18 |
| 2 | 231925646 | ['C', 'A'] | 1 | 1.00 | ['missense_variant'] | MODERATE | NPPC | ENST00000409852.2:c.160G>T | ENSP00000387159.1:p.Asp54Tyr | | 23 |
| 2 | 231925649 | ['CCTT', 'C' | 3 | 1.00 | ['inframe_deletion'] | MODERATE | NPPC | ENST00000409852.2:c.154_156del | ENSP00000387159.1:p.Lys52del | 4.44E-06 | |
| 2 | 231925649 | ['C', 'T'] | 1 | 1.00 | ['missense_variant'] | MODERATE | NPPC | ENST00000409852.2:c.157G>A | ENSP00000387159.1:p.Gly53Ser | | 21 |
| 2 | 231925657 | ['T', 'C'] | 3 | 1.00 | ['missense_variant'] | MODERATE | NPPC | ENST00000409852.2:c.149A>G | ENSP00000387159.1:p.Gln50Arg | 1.81E-05 | 21 |
| 2 | 231925663 | ['C', 'A'] | 1 | 1.00 | ['missense_variant'] | MODERATE | NPPC | ENST00000409852.2:c.143G>T | ENSP00000387159.1:p.Gly48Val | | 24 |
| 2 | 231925669 | ['G', 'C'] | 1 | 1.00 | ['missense_variant'] | MODERATE | NPPC | ENST00000409852.2:c.137C>G | ENSP00000387159.1:p.Ala46Gly | | 23 |
| 2 | 231925708 | ['C', 'T'] | 1 | 1.00 | ['missense_variant'] | MODERATE | NPPC | ENST00000409852.2:c.98G>A | ENSP00000387159.1:p.Arg33Gln | 5.65E-06 | 24 |
| 2 | 231925711 | ['G', 'A'] | 1 | 1.00 | ['missense_variant'] | MODERATE | NPPC | ENST00000409852.2:c.95C>T | ENSP00000387159.1:p.Pro32Leu | | 13 |
| 2 | 231926189 | ['A', 'G'] | 1 | 1.00 | ['missense_variant'] | MODERATE | NPPC | ENST00000409852.2:c.61T>C | ENSP00000387159.1:p.Ser21Pro | 3.40E-05 | 23 |
| 2 | 231926200 | ['G', 'C'] | 1 | 1.00 | ['missense_variant'] | MODERATE | NPPC | ENST00000409852.2:c.50C>G | ENSP00000387159.1:p.Ser17Cys | | 28 |
| 2 | 231926201 | ['A', 'AGA( | 1 | 1.00 | ['inframe_insertion'] | MODERATE | NPPC | ENST00000409852.2:c.40_48dup | ENSP00000387159.1:p.Thr14_Leu16dup | | |
| 4 | 1793974 | ['G', 'GTG( | 1 | 1.00 | ['inframe_insertion'] | MODERATE | FGFR3 | ENST00000340107.8:c.42_47dup | ENSP00000339824.4:p.Ala15_Ile16insMetAla | | |
| 4 | 1794009 | ['G', 'C'] | 1 | 1.00 | ['missense_variant'] | MODERATE | FGFR3 | ENST00000340107.8:c.75G>C | ENSP00000339824.4:p.Leu25Phe | | 8 |
| 4 | 1794016 | ['G', 'C'] | 2 | 1.00 | ['missense_variant'] | MODERATE | FGFR3 | ENST00000340107.8:c.82G>C | ENSP00000339824.4:p.Glu28Gln | | 22 |
| 4 | 1794043 | ['G', 'A'] | 1 | 1.00 | _variant', 'splice_regi( | MODERATE | FGFR3 | ENST00000340107.8:c.109G>A | ENSP00000339824.4:p.Glu37Lys | | 22 |
| 4 | 1799274 | ['G', 'A'] | 4 | 1.00 | ['missense_variant'] | MODERATE | FGFR3 | ENST00000340107.8:c.130G>A | ENSP00000339824.4:p.Gly44Ser | 0.000225 | 0 |
| 4 | 1799289 | ['T', 'G'] | 1 | 1.00 | ['missense_variant'] | MODERATE | FGFR3 | ENST00000340107.8:c.145T>G | ENSP00000339824.4:p.Leu49Val | 4.06E-06 | 5 |
| 4 | 1799292 | ['G', 'T'] | 7 | 1.00 | ['missense_variant'] | MODERATE | FGFR3 | ENST00000340107.8:c.148G>T | ENSP00000339824.4:p.Val50Phe | | 22 |
| 4 | 1799292 | ['G', 'A'] | 1 | 1.00 | ['missense_variant'] | MODERATE | FGFR3 | ENST00000340107.8:c.148G>A | ENSP00000339824.4:p.Val50Ile | | 13 |
| 4 | 1799303 | ['C', 'A'] | 1 | 1.00 | ['missense_variant'] | MODERATE | FGFR3 | ENST00000340107.8:c.159C>A | ENSP00000339824.4:p.Ser53Arg | | 11 |
| 4 | 1799304 | ['G', 'A'] | 1 | 1.00 | ['missense_variant'] | MODERATE | FGFR3 | ENST00000340107.8:c.160G>A | ENSP00000339824.4:p.Gly54Arg | 4.06E-05 | 23 |
| 4 | 1799310 | ['G', 'A'] | 1 | 1.00 | ['missense_variant'] | MODERATE | FGFR3 | ENST00000340107.8:c.166G>A | ENSP00000339824.4:p.Ala56Thr | | 0 |
| 4 | 1799316 | ['G', 'A'] | 3 | 1.00 | ['missense_variant'] | MODERATE | FGFR3 | ENST00000340107.8:c.172G>A | ENSP00000339824.4:p.Glu58Lys | 8.13E-06 | 15 |
| 4 | 1799328 | ['C', 'A'] | 1 | 1.00 | ['missense_variant'] | MODERATE | FGFR3 | ENST00000340107.8:c.184C>A | ENSP00000339824.4:p.Pro62Thr | | 1 |
| 4 | 1799332 | ['C', 'G'] | 5 | 1.00 | ['missense_variant'] | MODERATE | FGFR3 | ENST00000340107.8:c.188C>G | ENSP00000339824.4:p.Pro63Arg | 3.27E-05 | 3 |
| 4 | 1799337 | ['G', 'A'] | 3 | 1.00 | ['missense_variant'] | MODERATE | FGFR3 | ENST00000340107.8:c.193G>A | ENSP00000339824.4:p.Gly65Arg | 0.001076 | 11 |
| 4 | 1799389 | ['T', 'C'] | 2 | 1.00 | ['missense_variant'] | MODERATE | FGFR3 | ENST00000340107.8:c.245T>C | ENSP00000339824.4:p.Val82Ala | | 2 |
| 4 | 1799416 | ['C', 'T'] | 1 | 1.00 | ['missense_variant'] | MODERATE | FGFR3 | ENST00000340107.8:c.272C>T | ENSP00000339824.4:p.Pro91Leu | 0.000176 | 8 |
| 4 | 1799422 | ['G', 'A'] | 1 | 1.00 | ['missense_variant'] | MODERATE | FGFR3 | ENST00000340107.8:c.278G>A | ENSP00000339824.4:p.Arg93Gln | 1.69E-05 | 12 |
| 4 | 1799457 | ['G', 'A'] | 2 | 1.00 | ['missense_variant'] | MODERATE | FGFR3 | ENST00000340107.8:c.313G>A | ENSP00000339824.4:p.Gly105Arg | 9.26E-06 | 24 |
| 4 | 1799472 | ['C', 'T'] | 1 | 1.00 | ['missense_variant'] | MODERATE | FGFR3 | ENST00000340107.8:c.328C>T | ENSP00000339824.4:p.Arg110Trp | 1.50E-05 | 22 |
| 4 | 1799473 | ['G', 'A'] | 2 | 1.00 | ['missense_variant'] | MODERATE | FGFR3 | ENST00000340107.8:c.329G>A | ENSP00000339824.4:p.Arg110Gln | 7.05E-05 | 17 |
| 4 | 1799479 | ['G', 'A'] | 3 | 1.00 | ['missense_variant'] | MODERATE | FGFR3 | ENST00000340107.8:c.335G>A | ENSP00000339824.4:p.Arg112Gln | 0.000229 | 9 |
| 4 | 1799491 | ['G', 'A'] | 5 | 1.00 | ['missense_variant'] | MODERATE | FGFR3 | ENST00000340107.8:c.347G>A | ENSP00000339824.4:p.Arg116His | 4.18E-05 | 3 |
| 4 | 1799493 | ['G', 'A'] | 3 | 1.00 | ['missense_variant'] | MODERATE | FGFR3 | ENST00000340107.8:c.349G>A | ENSP00000339824.4:p.Val117Ile | 7.21E-05 | 0 |
| 4 | 1799496 | ['C', 'G'] | 1 | 1.00 | ['missense_variant'] | MODERATE | FGFR3 | ENST00000340107.8:c.352C>G | ENSP00000339824.4:p.Leu118Val | 1.22E-05 | 12 |
| 4 | 1799756 | ['C', 'A'] | 3 | 1.00 | ['missense_variant'] | MODERATE | FGFR3 | ENST00000340107.8:c.389C>A | ENSP00000339824.4:p.Ser130Tyr | | 19 |
| 4 | 1799756 | ['C', 'T'] | 1 | 1.00 | ['missense_variant'] | MODERATE | FGFR3 | ENST00000340107.8:c.389C>T | ENSP00000339824.4:p.Ser130Phe | 1.20E-05 | 17 |
| 4 | 1799758 | ['T', 'C'] | 1 | 1.00 | ['missense_variant'] | MODERATE | FGFR3 | ENST00000340107.8:c.391T>C | ENSP00000339824.4:p.Ser131Pro | 8.01E-06 | 22 |
| 4 | 1799770 | ['G', 'A'] | 6 | 1.00 | ['missense_variant'] | MODERATE | FGFR3 | ENST00000340107.8:c.403G>A | ENSP00000339824.4:p.Glu135Lys | 1.20E-05 | 21 |
| 4 | 1799776 | ['G', 'A'] | 1 | 1.00 | ['missense_variant'] | MODERATE | FGFR3 | ENST00000340107.8:c.409G>A | ENSP00000339824.4:p.Gly137Arg | 1.60E-05 | 15 |
| 4 | 1799792 | ['AGGACA( | 1 | 1.00 | ['inframe_deletion'] | MODERATE | FGFR3 | ENST00000340107.8:c.436_445+2del | | 1.20E-05 | |
| 4 | 1801373 | ['C', 'T'] | 1 | 1.00 | ['missense_variant'] | MODERATE | FGFR3 | ENST00000340107.8:c.452C>T | ENSP00000339824.4:p.Pro151Leu | | 27 |
| 4 | 1801384 | ['C', 'T'] | 1 | 1.00 | ['missense_variant'] | MODERATE | FGFR3 | ENST00000340107.8:c.463C>T | ENSP00000339824.4:p.Arg155Trp | 6.53E-06 | 24 |
| 4 | 1801393 | ['C', 'T'] | 1 | 1.00 | ['missense_variant'] | MODERATE | FGFR3 | ENST00000340107.8:c.472C>T | ENSP00000339824.4:p.Arg158Trp | 6.50E-06 | 23 |
| 4 | 1801394 | ['G', 'A'] | 5 | 1.00 | ['missense_variant'] | MODERATE | FGFR3 | ENST00000340107.8:c.473G>A | ENSP00000339824.4:p.Arg158Gln | 5.85E-05 | 27 |
| 4 | 1801402 | ['A', 'G'] | 1 | 1.00 | ['missense_variant'] | MODERATE | FGFR3 | ENST00000340107.8:c.481A>G | ENSP00000339824.4:p.Lys161Glu | | 31 |
| 4 | 1801435 | ['G', 'A'] | 3 | 1.00 | ['missense_variant'] | MODERATE | FGFR3 | ENST00000340107.8:c.514G>A | ENSP00000339824.4:p.Val172Ile | 6.67E-05 | 24 |
| 4 | 1801439 | ['G', 'A'] | 1 | 1.00 | ['missense_variant'] | MODERATE | FGFR3 | ENST00000340107.8:c.518G>A | ENSP00000339824.4:p.Arg173His | 4.00E-05 | 28 |
| 4 | 1801445 | ['G', 'A'] | 2 | 1.00 | ['missense_variant'] | MODERATE | FGFR3 | ENST00000340107.8:c.524G>A | ENSP00000339824.4:p.Arg175His | 3.98E-05 | 27 |
| 4 | 1801456 | ['G', 'A'] | 2 | 1.00 | ['missense_variant'] | MODERATE | FGFR3 | ENST00000340107.8:c.535G>A | ENSP00000339824.4:p.Ala179Thr | | 14 |
| 4 | 1801477 | ['A', 'G'] | 1 | 1.00 | ['missense_variant'] | MODERATE | FGFR3 | ENST00000340107.8:c.556A>G | ENSP00000339824.4:p.Ile186Val | | 23 |
| 4 | 1801478 | ['T', 'A'] | 1 | 1.00 | ['missense_variant'] | MODERATE | FGFR3 | ENST00000340107.8:c.557T>A | ENSP00000339824.4:p.Ile186Asn | | 25 |
| 4 | 1801481 | ['C', 'A'] | 4 | 1.00 | ['missense_variant'] | MODERATE | FGFR3 | ENST00000340107.8:c.560C>A | ENSP00000339824.4:p.Ser187Tyr | 2.52E-05 | 19 |
| 4 | 1801495 | ['G', 'A'] | 1 | 1.00 | ['missense_variant'] | MODERATE | FGFR3 | ENST00000340107.8:c.574G>A | ENSP00000339824.4:p.Gly192Ser | | 26 |

Supplementary Table 5 Genetic variants found in five height genes

| chr | position | alleles | AC | nt_qc.call_ | vep.consequence | vep.impact | vep.symbol | vep.hgvsc | vep.hgvsp | vep.gnom | vep.cadd_phred |
|---|---|---|---|---|---|---|---|---|---|---|---|
| 4 | 1801506 | ['C', 'G'] | 4 | 1.00 | ['missense_variant'] | MODERATE | FGFR3 | ENST00000340107.8:c.585C>G | ENSP00000339824.4:p.Phe195Leu | 1.84E-05 | 23 |
| 4 | 1801510 | ['G', 'A'] | 2 | 1.00 | ['missense_variant'] | MODERATE | FGFR3 | ENST00000340107.8:c.589G>A | ENSP00000339824.4:p.Gly197Ser | 6.04E-06 | 24 |
| 4 | 1801634 | ['G', 'T'] | 1 | 1.00 | ['missense_variant'] | MODERATE | FGFR3 | ENST00000340107.8:c.630G>T | ENSP00000339824.4:p.Gln210His | | 16 |
| 4 | 1801689 | ['G', 'A'] | 1 | 1.00 | ['missense_variant'] | MODERATE | FGFR3 | ENST00000340107.8:c.685G>A | ENSP00000339824.4:p.Val229Ile | | 12 |
| 4 | 1801717 | ['G', 'A'] | 7 | 1.00 | ['missense_variant'] | MODERATE | FGFR3 | ENST00000340107.8:c.713G>A | ENSP00000339824.4:p.Arg238Gln | 0.0005 | 9 |
| 4 | 1801743 | ['G', 'A'] | 1 | 1.00 | _variant', 'splice_regi | MODERATE | FGFR3 | ENST00000340107.8:c.739G>A | ENSP00000339824.4:p.Glu247Lys | 1.34E-05 | 34 |
| 4 | 1801841 | ['C', 'T'] | 1 | 1.00 | ['missense_variant'] | MODERATE | FGFR3 | ENST00000340107.8:c.746C>T | ENSP00000339824.4:p.Ser249Phe | 1.26E-05 | 25 |
| 4 | 1801849 | ['C', 'T'] | 1 | 1.00 | ['missense_variant'] | MODERATE | FGFR3 | ENST00000340107.8:c.754C>T | ENSP00000339824.4:p.Arg252Trp | | 25 |
| 4 | 1801874 | ['C', 'T'] | 8 | 1.00 | ['missense_variant'] | MODERATE | FGFR3 | ENST00000340107.8:c.779C>T | ENSP00000339824.4:p.Pro260Leu | 2.51E-05 | 28 |
| 4 | 1801903 | ['G', 'A'] | 1 | 1.00 | ['missense_variant'] | MODERATE | FGFR3 | ENST00000340107.8:c.808G>A | ENSP00000339824.4:p.Asp270Asn | 9.32E-05 | 22 |
| 4 | 1801948 | ['A', 'G'] | 4 | 1.00 | ['missense_variant'] | MODERATE | FGFR3 | ENST00000340107.8:c.853A>G | ENSP00000339824.4:p.Ile285Val | 2.41E-05 | 26 |
| 4 | 1802005 | ['C', 'T'] | 1 | 1.00 | ['missense_variant'] | MODERATE | FGFR3 | ENST00000340107.8:c.910C>T | ENSP00000339824.4:p.Pro304Ser | | 26 |
| 4 | 1802941 | ['G', 'A'] | 1 | 1.00 | ['missense_variant'] | MODERATE | FGFR3 | ENST00000340107.8:c.958G>A | ENSP00000339824.4:p.Asp320Asn | 4.33E-06 | 23 |
| 4 | 1802944 | ['G', 'A'] | 1 | 1.00 | ['missense_variant'] | MODERATE | FGFR3 | ENST00000340107.8:c.961G>A | ENSP00000339824.4:p.Val321Met | 2.60E-05 | 20 |
| 4 | 1802948 | ['G', 'A'] | 3 | 1.00 | ['missense_variant'] | MODERATE | FGFR3 | ENST00000340107.8:c.965G>A | ENSP00000339824.4:p.Arg322His | 8.68E-06 | 23 |
| 4 | 1802953 | ['C', 'T'] | 1 | 1.00 | ['missense_variant'] | MODERATE | FGFR3 | ENST00000340107.8:c.970C>T | ENSP00000339824.4:p.Arg324Cys | 1.30E-05 | 22 |
| 4 | 1802954 | ['G', 'A'] | 1 | 1.00 | ['missense_variant'] | MODERATE | FGFR3 | ENST00000340107.8:c.971G>A | ENSP00000339824.4:p.Arg324His | 1.73E-05 | 18 |
| 4 | 1802969 | ['C', 'T'] | 1 | 1.00 | ['missense_variant'] | MODERATE | FGFR3 | ENST00000340107.8:c.986C>T | ENSP00000339824.4:p.Ser329Leu | 3.07E-05 | 24 |
| 4 | 1802975 | ['G', 'A'] | 3 | 1.00 | ['missense_variant'] | MODERATE | FGFR3 | ENST00000340107.8:c.992G>A | ENSP00000339824.4:p.Arg331Gln | 2.19E-05 | 18 |
| 4 | 1802986 | ['G', 'A'] | 1 | 1.00 | ['missense_variant'] | MODERATE | FGFR3 | ENST00000340107.8:c.1003G>A | ENSP00000339824.4:p.Glu335Lys | 4.89E-05 | 20 |
| 4 | 1803005 | ['C', 'T'] | 1 | 1.00 | ['missense_variant'] | MODERATE | FGFR3 | ENST00000340107.8:c.1022C>T | ENSP00000339824.4:p.Thr341Ile | | 22 |
| 4 | 1803019 | ['G', 'A'] | 1 | 1.00 | ['missense_variant'] | MODERATE | FGFR3 | ENST00000340107.8:c.1036G>A | ENSP00000339824.4:p.Val346Met | 1.74E-05 | 22 |
| 4 | 1803046 | ['G', 'A'] | 3 | 1.00 | ['missense_variant'] | MODERATE | FGFR3 | ENST00000340107.8:c.1063G>A | ENSP00000339824.4:p.Val355Ile | 1.34E-05 | 24 |
| 4 | 1803052 | ['G', 'A'] | 1 | 1.00 | ['missense_variant'] | MODERATE | FGFR3 | ENST00000340107.8:c.1069G>A | ENSP00000339824.4:p.Gly357Arg | 2.25E-05 | 17 |
| 4 | 1803054 | ['G', 'GC'] | 3 | 1.00 | ['frameshift_variant'] | HIGH | FGFR3 | ENST00000340107.8:c.1075dup | ENSP00000339824.4:p.Arg359ProfsTer11 | | |
| 4 | 1803058 | ['C', 'T'] | 1 | 1.00 | ['stop_gained'] | HIGH | FGFR3 | ENST00000340107.8:c.1075C>T | ENSP00000339824.4:p.Arg359Ter | 1.82E-05 | 39 |
| 4 | 1803059 | ['G', 'A'] | 5 | 1.00 | ['missense_variant'] | MODERATE | FGFR3 | ENST00000340107.8:c.1076G>A | ENSP00000339824.4:p.Arg359Gln | 3.64E-05 | 9 |
| 4 | 1803061 | ['G', 'A'] | 1 | 1.00 | ['missense_variant'] | MODERATE | FGFR3 | ENST00000340107.8:c.1078G>A | ENSP00000339824.4:p.Ala360Thr | | 18 |
| 4 | 1804332 | ['G', 'C'] | 5 | 1.00 | _variant', 'splice_regi | MODERATE | FGFR3 | ENST00000340107.8:c.1084G>C | ENSP00000339824.4:p.Glu362Gln | 1.24E-05 | 15 |
| 4 | 1804353 | ['G', 'T'] | 1 | 1.00 | ['missense_variant'] | MODERATE | FGFR3 | ENST00000340107.8:c.1105G>T | ENSP00000339824.4:p.Asp369Tyr | | 20 |
| 4 | 1804360 | ['C', 'T'] | 4 | 1.00 | ['missense_variant'] | MODERATE | FGFR3 | ENST00000340107.8:c.1112C>T | ENSP00000339824.4:p.Ala371Val | 2.81E-05 | 18 |
| 4 | 1804375 | ['C', 'G'] | 1 | 1.00 | ['missense_variant'] | MODERATE | FGFR3 | ENST00000340107.8:c.1127C>G | ENSP00000339824.4:p.Ala376Gly | | 19 |
| 4 | 1804381 | ['T', 'G'] | 1 | 1.00 | ['missense_variant'] | MODERATE | FGFR3 | ENST00000340107.8:c.1133T>G | ENSP00000339824.4:p.Ile378Ser | | 26 |
| 4 | 1804383 | ['C', 'A'] | 1 | 1.00 | ['missense_variant'] | MODERATE | FGFR3 | ENST00000340107.8:c.1135C>A | ENSP00000339824.4:p.Leu379Ile | | 9 |
| 4 | 1804388 | ['C', 'A'] | 1 | 1.00 | ['missense_variant'] | MODERATE | FGFR3 | ENST00000340107.8:c.1140C>A | ENSP00000339824.4:p.Ser380Arg | | 20 |
| 4 | 1804401 | ['T', 'C'] | 2 | 1.00 | ['missense_variant'] | MODERATE | FGFR3 | ENST00000340107.8:c.1153T>C | ENSP00000339824.4:p.Phe385Leu | 7.99E-06 | 19 |
| 4 | 1804404 | ['T', 'A'] | 2 | 0.99 | ['missense_variant'] | MODERATE | FGFR3 | ENST00000340107.8:c.1156T>A | ENSP00000339824.4:p.Phe386Ile | 7.99E-06 | 18 |
| 4 | 1804413 | ['A', 'T'] | 1 | 1.00 | ['missense_variant'] | MODERATE | FGFR3 | ENST00000340107.8:c.1165A>T | ENSP00000339824.4:p.Ile389Phe | | 18 |
| 4 | 1804414 | ['T', 'A'] | 1 | 1.00 | ['missense_variant'] | MODERATE | FGFR3 | ENST00000340107.8:c.1166T>A | ENSP00000339824.4:p.Ile389Asn | | 24 |
| 4 | 1804423 | ['T', 'TGGC | 1 | 1.00 | ['inframe_insertion'] | MODERATE | FGFR3 | ENST00000340107.8:c.1179_1181dup | ENSP00000339824.4:p.Ala394dup | | |
| 4 | 1804443 | ['C', 'T'] | 1 | 1.00 | ['missense_variant'] | MODERATE | FGFR3 | ENST00000340107.8:c.1195C>T | ENSP00000339824.4:p.Arg399Cys | 3.21E-05 | 25 |
| 4 | 1804449 | ['C', 'T'] | 5 | 1.00 | ['missense_variant'] | MODERATE | FGFR3 | ENST00000340107.8:c.1201C>T | ENSP00000339824.4:p.Arg401Cys | 1.60E-05 | 23 |
| 4 | 1804455 | ['C', 'G'] | 4 | 1.00 | ['missense_variant'] | MODERATE | FGFR3 | ENST00000340107.8:c.1207C>G | ENSP00000339824.4:p.Pro403Ala | 2.40E-05 | 9 |
| 4 | 1804464 | ['A', 'G'] | 1 | 1.00 | ['missense_variant'] | MODERATE | FGFR3 | ENST00000340107.8:c.1216A>G | ENSP00000339824.4:p.Lys406Glu | 4.00E-06 | 23 |
| 4 | 1804486 | ['T', 'G'] | 2 | 1.00 | ['missense_variant'] | MODERATE | FGFR3 | ENST00000340107.8:c.1238T>G | ENSP00000339824.4:p.Val413Gly | | 31 |
| 4 | 1804492 | ['A', 'G'] | 1 | 1.00 | ['missense_variant'] | MODERATE | FGFR3 | ENST00000340107.8:c.1244A>G | ENSP00000339824.4:p.Lys415Arg | 8.01E-06 | 25 |
| 4 | 1804493 | ['G', 'C'] | 1 | 1.00 | ['missense_variant'] | MODERATE | FGFR3 | ENST00000340107.8:c.1245G>C | ENSP00000339824.4:p.Lys415Asn | 1.20E-05 | 26 |
| 4 | 1804496 | ['C', 'G'] | 7 | 1.00 | ['missense_variant'] | MODERATE | FGFR3 | ENST00000340107.8:c.1248C>G | ENSP00000339824.4:p.Ile416Met | | 20 |
| 4 | 1804516 | ['G', 'A'] | 3 | 1.00 | ['missense_variant'] | MODERATE | FGFR3 | ENST00000340107.8:c.1268G>A | ENSP00000339824.4:p.Arg423Gln | 2.41E-05 | 29 |
| 4 | 1804824 | ['G', 'C'] | 1 | 1.00 | _variant', 'splice_regi | MODERATE | FGFR3 | ENST00000340107.8:c.1273G>C | ENSP00000339824.4:p.Val425Leu | 6.41E-06 | 28 |
| 4 | 1804828 | ['C', 'T'] | 2 | 1.00 | ['missense_variant'] | MODERATE | FGFR3 | ENST00000340107.8:c.1277C>T | ENSP00000339824.4:p.Ser426Phe | 1.28E-05 | 25 |
| 4 | 1804846 | ['C', 'T'] | 1 | 1.00 | ['missense_variant'] | MODERATE | FGFR3 | ENST00000340107.8:c.1295C>T | ENSP00000339824.4:p.Ser432Phe | | 29 |
| 4 | 1804872 | ['C', 'G'] | 6 | 1.00 | ['missense_variant'] | MODERATE | FGFR3 | ENST00000340107.8:c.1321C>G | ENSP00000339824.4:p.Arg441Gly | | 18 |
| 4 | 1804872 | ['C', 'T'] | 1 | 1.00 | ['missense_variant'] | MODERATE | FGFR3 | ENST00000340107.8:c.1321C>T | ENSP00000339824.4:p.Arg441Cys | 2.56E-05 | 23 |
| 4 | 1804873 | ['G', 'A'] | 3 | 1.00 | ['missense_variant'] | MODERATE | FGFR3 | ENST00000340107.8:c.1322G>A | ENSP00000339824.4:p.Arg441His | 3.84E-05 | 27 |
| 4 | 1804878 | ['G', 'A'] | 3 | 1.00 | ['missense_variant'] | MODERATE | FGFR3 | ENST00000340107.8:c.1327G>A | ENSP00000339824.4:p.Ala443Thr | 1.28E-05 | 11 |
| 4 | 1804888 | ['C', 'T'] | 1 | 1.00 | ['missense_variant'] | MODERATE | FGFR3 | ENST00000340107.8:c.1337C>T | ENSP00000339824.4:p.Ser446Phe | 2.56E-05 | 24 |
| 4 | 1804902 | ['C', 'G'] | 1 | 0.99 | ['missense_variant'] | MODERATE | FGFR3 | ENST00000340107.8:c.1351C>G | ENSP00000339824.4:p.Pro451Ala | | 20 |
| 4 | 1804906 | ['C', 'T'] | 4 | 1.00 | ['missense_variant'] | MODERATE | FGFR3 | ENST00000340107.8:c.1355C>T | ENSP00000339824.4:p.Thr452Met | 0.001288 | 6 |
| 4 | 1804911 | ['G', 'T'] | 2 | 1.00 | ['missense_variant'] | MODERATE | FGFR3 | ENST00000340107.8:c.1360G>T | ENSP00000339824.4:p.Ala454Ser | 2.56E-05 | 20 |
| 4 | 1804929 | ['G', 'C'] | 5 | 1.00 | ['missense_variant'] | MODERATE | FGFR3 | ENST00000340107.8:c.1378G>C | ENSP00000339824.4:p.Glu460Gln | 6.39E-06 | 26 |
| 4 | 1804941 | ['G', 'A'] | 1 | 1.00 | ['missense_variant'] | MODERATE | FGFR3 | ENST00000340107.8:c.1390G>A | ENSP00000339824.4:p.Asp464Asn | | 26 |
| 4 | 1804968 | ['C', 'G'] | 2 | 1.00 | _variant', 'splice_regi | MODERATE | FGFR3 | ENST00000340107.8:c.1417C>G | ENSP00000339824.4:p.Arg473Gly | | 23 |
| 4 | 1805405 | ['T', 'C'] | 1 | 1.00 | ['missense_variant'] | MODERATE | FGFR3 | ENST00000340107.8:c.1469T>C | ENSP00000339824.4:p.Met490Thr | | 25 |
| 4 | 1805416 | ['A', 'G'] | 1 | 1.00 | ['missense_variant'] | MODERATE | FGFR3 | ENST00000340107.8:c.1480A>G | ENSP00000339824.4:p.Ile494Val | 4.03E-06 | 13 |
| 4 | 1805428 | ['A', 'G'] | 1 | 1.00 | ['missense_variant'] | MODERATE | FGFR3 | ENST00000340107.8:c.1492A>G | ENSP00000339824.4:p.Lys498Glu | 8.07E-06 | 29 |
| 4 | 1805434 | ['C', 'T'] | 1 | 1.00 | ['missense_variant'] | MODERATE | FGFR3 | ENST00000340107.8:c.1498C>T | ENSP00000339824.4:p.Arg500Trp | 8.09E-05 | 24 |
| 4 | 1805435 | ['G', 'A'] | 2 | 1.00 | ['missense_variant'] | MODERATE | FGFR3 | ENST00000340107.8:c.1499G>A | ENSP00000339824.4:p.Arg500Gln | 8.08E-06 | 23 |
| 4 | 1805440 | ['G', 'A'] | 4 | 1.00 | ['missense_variant'] | MODERATE | FGFR3 | ENST00000340107.8:c.1504G>A | ENSP00000339824.4:p.Ala502Thr | 5.66E-05 | 21 |

Supplementary Table 5 Genetic variants found in five height genes

| chr | position | alleles | AC | nt_qc.call_ | vep.consequence | vep.impact | vep.symbol | vep.hgvsc | vep.hgvsp | vep.gnom | vep.cadd_phred |
|---|---|---|---|---|---|---|---|---|---|---|---|
| 4 | 1805561 | ['G', 'A'] | 1 | 1.00 | _variant', 'splice_regi | MODERATE | FGFR3 | ENST00000340107.8:c.1543G>A | ENSP00000339824.4:p.Asp515Asn | 4.79E-05 | 23 |
| 4 | 1805571 | ['A', 'C'] | 1 | 1.00 | ['missense_variant'] | MODERATE | FGFR3 | ENST00000340107.8:c.1553A>C | ENSP00000339824.4:p.Asp518Ala | 3.99E-06 | 27 |
| 4 | 1805571 | ['A', 'T'] | 1 | 1.00 | ['missense_variant'] | MODERATE | FGFR3 | ENST00000340107.8:c.1553A>T | ENSP00000339824.4:p.Asp518Val | | 27 |
| 4 | 1805573 | ['A', 'C'] | 1 | 1.00 | ['missense_variant'] | MODERATE | FGFR3 | ENST00000340107.8:c.1555A>C | ENSP00000339824.4:p.Lys519Gln | 1.20E-05 | 25 |
| 4 | 1805573 | ['A', 'G'] | 1 | 1.00 | ['missense_variant'] | MODERATE | FGFR3 | ENST00000340107.8:c.1555A>G | ENSP00000339824.4:p.Lys519Glu | 3.99E-06 | 23 |
| 4 | 1805574 | ['A', 'G'] | 3 | 1.00 | ['missense_variant'] | MODERATE | FGFR3 | ENST00000340107.8:c.1556A>G | ENSP00000339824.4:p.Lys519Arg | 5.58E-05 | 24 |
| 4 | 1805583 | ['C', 'T'] | 3 | 1.00 | ['missense_variant'] | MODERATE | FGFR3 | ENST00000340107.8:c.1565C>T | ENSP00000339824.4:p.Ser522Leu | 7.98E-06 | 25 |
| 4 | 1805636 | ['A', 'G'] | 3 | 1.00 | ['missense_variant'] | MODERATE | FGFR3 | ENST00000340107.8:c.1618A>G | ENSP00000339824.4:p.Ile540Val | 1.20E-05 | 22 |
| 4 | 1805642 | ['A', 'G'] | 1 | 1.00 | ['missense_variant'] | MODERATE | FGFR3 | ENST00000340107.8:c.1624A>G | ENSP00000339824.4:p.Asn542Asp | 3.99E-06 | 28 |
| 4 | 1805643 | ['A', 'G'] | 2 | 1.00 | ['missense_variant'] | MODERATE | FGFR3 | ENST00000340107.8:c.1625A>G | ENSP00000339824.4:p.Asn542Ser | 3.99E-06 | 26 |
| 4 | 1805649 | ['T', 'G'] | 2 | 1.00 | ['missense_variant'] | MODERATE | FGFR3 | ENST00000340107.8:c.1631T>G | ENSP00000339824.4:p.Leu544Arg | | 32 |
| 4 | 1805661 | ['C', 'T'] | 1 | 1.00 | ['missense_variant'] | MODERATE | FGFR3 | ENST00000340107.8:c.1643C>T | ENSP00000339824.4:p.Thr548Met | | 25 |
| 4 | 1805666 | ['G', 'A'] | 1 | 1.00 | ['missense_variant'] | MODERATE | FGFR3 | ENST00000340107.8:c.1648G>A | ENSP00000339824.4:p.Gly550Ser | | 25 |
| 4 | 1805761 | ['G', 'A'] | 2 | 1.00 | ['missense_variant'] | MODERATE | FGFR3 | ENST00000340107.8:c.1663G>A | ENSP00000339824.4:p.Val555Met | 0.000146 | 25 |
| 4 | 1805761 | ['G', 'C'] | 1 | 1.00 | ['missense_variant'] | MODERATE | FGFR3 | ENST00000340107.8:c.1663G>C | ENSP00000339824.4:p.Val555Leu | 4.05E-06 | 24 |
| 4 | 1805774 | ['A', 'G'] | 1 | 1.00 | ['missense_variant'] | MODERATE | FGFR3 | ENST00000340107.8:c.1676A>G | ENSP00000339824.4:p.Tyr559Cys | | 21 |
| 4 | 1805779 | ['G', 'C'] | 4 | 1.00 | ['missense_variant'] | MODERATE | FGFR3 | ENST00000340107.8:c.1681G>C | ENSP00000339824.4:p.Ala561Pro | 2.02E-05 | 23 |
| 4 | 1805807 | ['G', 'A'] | 1 | 1.00 | ['missense_variant'] | MODERATE | FGFR3 | ENST00000340107.8:c.1709G>A | ENSP00000339824.4:p.Arg570Gln | | 24 |
| 4 | 1805810 | ['C', 'T'] | 7 | 1.00 | ['missense_variant'] | MODERATE | FGFR3 | ENST00000340107.8:c.1712C>T | ENSP00000339824.4:p.Ala571Val | 0.00015 | 26 |
| 4 | 1805819 | ['C', 'T'] | 3 | 1.00 | ['missense_variant'] | MODERATE | FGFR3 | ENST00000340107.8:c.1721C>T | ENSP00000339824.4:p.Pro574Leu | 2.03E-05 | 25 |
| 4 | 1805822 | ['C', 'T'] | 6 | 1.00 | ['missense_variant'] | MODERATE | FGFR3 | ENST00000340107.8:c.1724C>T | ENSP00000339824.4:p.Pro575Leu | | 26 |
| 4 | 1805837 | ['C', 'T'] | 1 | 1.00 | ['missense_variant'] | MODERATE | FGFR3 | ENST00000340107.8:c.1739C>T | ENSP00000339824.4:p.Ser580Phe | 9.69E-05 | 24 |
| 4 | 1805860 | ['G', 'A'] | 3 | 1.00 | ['missense_variant'] | MODERATE | FGFR3 | ENST00000340107.8:c.1762G>A | ENSP00000339824.4:p.Glu588Lys | 6.83E-05 | 23 |
| 4 | 1805911 | ['C', 'T'] | 1 | 1.00 | ['missense_variant'] | MODERATE | FGFR3 | ENST00000340107.8:c.1813C>T | ENSP00000339824.4:p.Arg605Trp | 4.01E-06 | 25 |
| 4 | 1805920 | ['G', 'C'] | 1 | 1.00 | ['missense_variant'] | MODERATE | FGFR3 | ENST00000340107.8:c.1822G>C | ENSP00000339824.4:p.Glu608Gln | | 27 |
| 4 | 1805921 | ['A', 'G'] | 2 | 1.00 | ['missense_variant'] | MODERATE | FGFR3 | ENST00000340107.8:c.1823A>G | ENSP00000339824.4:p.Glu608Gly | 4.02E-06 | 33 |
| 4 | 1805929 | ['G', 'A'] | 1 | 1.00 | ['missense_variant'] | MODERATE | FGFR3 | ENST00000340107.8:c.1831G>A | ENSP00000339824.4:p.Ala611Thr | 4.03E-06 | 33 |
| 4 | 1805930 | ['C', 'T'] | 3 | 1.00 | ['missense_variant'] | MODERATE | FGFR3 | ENST00000340107.8:c.1832C>T | ENSP00000339824.4:p.Ala611Val | 2.42E-05 | 25 |
| 4 | 1805933 | ['C', 'A'] | 5 | 1.00 | ['missense_variant'] | MODERATE | FGFR3 | ENST00000340107.8:c.1835C>A | ENSP00000339824.4:p.Ser612Tyr | 1.21E-05 | 27 |
| 4 | 1806093 | ['G', 'A'] | 6 | 1.00 | ['missense_variant'] | MODERATE | FGFR3 | ENST00000340107.8:c.1885G>A | ENSP00000339824.4:p.Glu629Lys | 0.00018 | 33 |
| 4 | 1806169 | ['C', 'G'] | 4 | 1.00 | ['missense_variant'] | MODERATE | FGFR3 | ENST00000340107.8:c.1961C>G | ENSP00000339824.4:p.Thr654Ser | 8.04E-06 | 20 |
| 4 | 1806281 | ['G', 'C'] | 1 | 1.00 | ['missense_variant'] | MODERATE | FGFR3 | ENST00000340107.8:c.1990G>C | ENSP00000339824.4:p.Ala664Pro | 4.04E-06 | 33 |
| 4 | 1806290 | ['G', 'T'] | 1 | 1.00 | ['missense_variant'] | MODERATE | FGFR3 | ENST00000340107.8:c.1999G>T | ENSP00000339824.4:p.Ala667Ser | 1.21E-05 | 25 |
| 4 | 1806300 | ['ACC', 'A'] | 1 | 1.00 | ['frameshift_variant'] | HIGH | FGFR3 | ENST00000340107.8:c.2010_2011del | ENSP00000339824.4:p.Asp670GlufsTer8 | | |
| 4 | 1806315 | ['A', 'G'] | 1 | 1.00 | ['missense_variant'] | MODERATE | FGFR3 | ENST00000340107.8:c.2024A>G | ENSP00000339824.4:p.His675Arg | | 25 |
| 4 | 1806326 | ['G', 'A'] | 1 | 1.00 | _variant', 'splice_regi | MODERATE | FGFR3 | ENST00000340107.8:c.2035G>A | ENSP00000339824.4:p.Val679Ile | 4.02E-06 | 27 |
| 4 | 1806548 | ['G', 'T'] | 1 | 1.00 | _variant', 'splice_regi | MODERATE | FGFR3 | ENST00000340107.8:c.2039G>T | ENSP00000339824.4:p.Trp680Leu | | 34 |
| 4 | 1806581 | ['C', 'T'] | 2 | 1.00 | ['missense_variant'] | MODERATE | FGFR3 | ENST00000340107.8:c.2072C>T | ENSP00000339824.4:p.Thr691Met | 4.00E-06 | 26 |
| 4 | 1806593 | ['C', 'T'] | 6 | 1.00 | ['missense_variant'] | MODERATE | FGFR3 | ENST00000340107.8:c.2084C>T | ENSP00000339824.4:p.Ser695Phe | 1.20E-05 | 25 |
| 4 | 1806619 | ['G', 'A'] | 1 | 1.00 | ['missense_variant'] | MODERATE | FGFR3 | ENST00000340107.8:c.2110G>A | ENSP00000339824.4:p.Glu704Lys | 4.00E-06 | 27 |
| 4 | 1806668 | ['A', 'G'] | 5 | 1.00 | ['missense_variant'] | MODERATE | FGFR3 | ENST00000340107.8:c.2159A>G | ENSP00000339824.4:p.Asn720Ser | 6.81E-05 | 23 |
| 4 | 1806669 | ['C', 'G'] | 1 | 1.00 | ['missense_variant'] | MODERATE | FGFR3 | ENST00000340107.8:c.2160C>G | ENSP00000339824.4:p.Asn720Lys | | 24 |
| 4 | 1806842 | ['C', 'T'] | 1 | 1.00 | ['missense_variant'] | MODERATE | FGFR3 | ENST00000340107.8:c.2188C>T | ENSP00000339824.4:p.Arg730Trp | 4.20E-06 | 23 |
| 4 | 1806843 | ['G', 'A'] | 5 | 1.00 | ['missense_variant'] | MODERATE | FGFR3 | ENST00000340107.8:c.2189G>A | ENSP00000339824.4:p.Arg730Gln | 3.35E-05 | 26 |
| 4 | 1806855 | ['A', 'G'] | 2 | 1.00 | ['missense_variant'] | MODERATE | FGFR3 | ENST00000340107.8:c.2201A>G | ENSP00000339824.4:p.His734Arg | 4.12E-06 | 21 |
| 4 | 1806860 | ['G', 'A'] | 1 | 1.00 | ['missense_variant'] | MODERATE | FGFR3 | ENST00000340107.8:c.2206G>A | ENSP00000339824.4:p.Ala736Thr | 4.93E-05 | 22 |
| 4 | 1806861 | ['C', 'T'] | 1 | 1.00 | ['missense_variant'] | MODERATE | FGFR3 | ENST00000340107.8:c.2207C>T | ENSP00000339824.4:p.Ala736Val | 3.29E-05 | 3 |
| 4 | 1806876 | ['C', 'T'] | 1 | 1.00 | ['missense_variant'] | MODERATE | FGFR3 | ENST00000340107.8:c.2222C>T | ENSP00000339824.4:p.Pro741Leu | | 26 |
| 4 | 1806935 | ['G', 'A'] | 2 | 1.00 | splice_donor_variant' | HIGH | FGFR3 | ENST00000340107.8:c.2280+1G>A | | 4.58E-06 | 25 |
| 4 | 1807132 | ['C', 'T'] | 7 | 1.00 | ['missense_variant'] | MODERATE | FGFR3 | ENST00000340107.8:c.2297C>T | ENSP00000339824.4:p.Ser766Leu | 5.47E-05 | 27 |
| 4 | 1807135 | ['C', 'T'] | 5 | 1.00 | ['missense_variant'] | MODERATE | FGFR3 | ENST00000340107.8:c.2300C>T | ENSP00000339824.4:p.Ala767Val | 6.36E-05 | 13 |
| 4 | 1807156 | ['C', 'T'] | 2 | 1.00 | ['missense_variant'] | MODERATE | FGFR3 | ENST00000340107.8:c.2321C>T | ENSP00000339824.4:p.Pro774Leu | 1.84E-05 | 24 |
| 4 | 1807159 | ['G', 'A'] | 3 | 1.00 | ['missense_variant'] | MODERATE | FGFR3 | ENST00000340107.8:c.2324G>A | ENSP00000339824.4:p.Gly775Asp | 3.19E-05 | 22 |
| 4 | 1807165 | ['A', 'G'] | 1 | 1.00 | ['missense_variant'] | MODERATE | FGFR3 | ENST00000340107.8:c.2330A>G | ENSP00000339824.4:p.Gln777Arg | 4.50E-06 | 20 |
| 4 | 1807191 | ['G', 'C'] | 2 | 1.00 | ['missense_variant'] | MODERATE | FGFR3 | ENST00000340107.8:c.2356G>C | ENSP00000339824.4:p.Gly786Arg | | 24 |
| 4 | 1807194 | ['G', 'GA'] | 1 | 1.00 | ['frameshift_variant'] | HIGH | FGFR3 | ENST00000340107.8:c.2360dup | ENSP00000339824.4:p.Asp787GlufsTer32 | | |
| 4 | 1807197 | ['G', 'A'] | 1 | 1.00 | ['missense_variant'] | MODERATE | FGFR3 | ENST00000340107.8:c.2362G>A | ENSP00000339824.4:p.Asp788Asn | 8.61E-06 | 27 |
| 4 | 1807201 | ['CCGT', 'C | 1 | 1.00 | ['inframe_deletion'] | MODERATE | FGFR3 | ENST00000340107.8:c.2367_2369del | ENSP00000339824.4:p.Val790del | | |
| 4 | 1807204 | ['T', 'C'] | 2 | 1.00 | ['missense_variant'] | MODERATE | FGFR3 | ENST00000340107.8:c.2369T>C | ENSP00000339824.4:p.Val790Ala | | 27 |
| 4 | 1807230 | ['G', 'A'] | 1 | 1.00 | ['missense_variant'] | MODERATE | FGFR3 | ENST00000340107.8:c.2395G>A | ENSP00000339824.4:p.Ala799Thr | | 17 |
| 4 | 1807237 | ['C', 'T'] | 1 | 1.00 | ['missense_variant'] | MODERATE | FGFR3 | ENST00000340107.8:c.2402C>T | ENSP00000339824.4:p.Pro801Leu | 8.24E-05 | 22 |
| 4 | 1807254 | ['C', 'T'] | 5 | 1.00 | ['missense_variant'] | MODERATE | FGFR3 | ENST00000340107.8:c.2419C>T | ENSP00000339824.4:p.Arg807Trp | 1.83E-05 | 14 |
| 4 | 1807255 | ['G', 'A'] | 2 | 1.00 | ['missense_variant'] | MODERATE | FGFR3 | ENST00000340107.8:c.2420G>A | ENSP00000339824.4:p.Arg807Gln | 2.79E-05 | 15 |
| 4 | 1807258 | ['C', 'A'] | 2 | 1.00 | ['missense_variant'] | MODERATE | FGFR3 | ENST00000340107.8:c.2423C>A | ENSP00000339824.4:p.Thr808Lys | 4.67E-06 | 24 |
| 9 | 35792461 | ['G', 'A'] | 2 | 1.00 | ['missense_variant'] | MODERATE | NPR2 | ENST00000342694.6:c.53G>A | ENSP00000341083.2:p.Arg18His | | 23 |
| 9 | 35792469 | ['G', 'C'] | 1 | 1.00 | ['missense_variant'] | MODERATE | NPR2 | ENST00000342694.6:c.61G>C | ENSP00000341083.2:p.Gly21Arg | | 19 |
| 9 | 35792475 | ['C', 'T'] | 1 | 1.00 | ['missense_variant'] | MODERATE | NPR2 | ENST00000342694.6:c.67C>T | ENSP00000341083.2:p.Arg23Trp | | 24 |
| 9 | 35792580 | ['C', 'T'] | 1 | 1.00 | ['missense_variant'] | MODERATE | NPR2 | ENST00000342694.6:c.172C>T | ENSP00000341083.2:p.Arg58Trp | 3.99E-06 | 25 |
| 9 | 35792581 | ['G', 'T'] | 1 | 1.00 | ['missense_variant'] | MODERATE | NPR2 | ENST00000342694.6:c.173G>T | ENSP00000341083.2:p.Arg58Leu | 1.20E-05 | 23 |

Supplementary Table 5 Genetic variants found in five height genes

| chr | position | alleles | AC | nt_qc.call_ | vep.consequence | vep.impact | vep.symbol | vep.hgvsc | vep.hgvsp | vep.gnoma | vep.cadd_phred |
|---|---|---|---|---|---|---|---|---|---|---|---|
| 9 | 35792583 | ['G', 'A'] | 2 | 1.00 | ['missense_variant'] | MODERATE | NPR2 | ENST00000342694.6:c.175G>A | ENSP00000341083.2:p.Ala59Thr | | 21 |
| 9 | 35792601 | ['C', 'T'] | 1 | 1.00 | ['missense_variant'] | MODERATE | NPR2 | ENST00000342694.6:c.193C>T | ENSP00000341083.2:p.Arg65Trp | 7.97E-06 | 23 |
| 9 | 35792652 | ['C', 'A'] | 1 | 1.00 | ['missense_variant'] | MODERATE | NPR2 | ENST00000342694.6:c.244C>A | ENSP00000341083.2:p.Leu82Met | 2.39E-05 | 23 |
| 9 | 35792688 | ['G', 'A'] | 1 | 1.00 | ['missense_variant'] | MODERATE | NPR2 | ENST00000342694.6:c.280G>A | ENSP00000341083.2:p.Asp94Asn | 7.96E-06 | 19 |
| 9 | 35792709 | ['T', 'G'] | 1 | 1.00 | ['missense_variant'] | MODERATE | NPR2 | ENST00000342694.6:c.301T>G | ENSP00000341083.2:p.Cys101Gly | | 27 |
| 9 | 35792713 | ['T', 'C'] | 1 | 1.00 | ['missense_variant'] | MODERATE | NPR2 | ENST00000342694.6:c.305T>C | ENSP00000341083.2:p.Val102Ala | | 20 |
| 9 | 35792736 | ['C', 'G'] | 1 | 1.00 | ['missense_variant'] | MODERATE | NPR2 | ENST00000342694.6:c.328C>G | ENSP00000341083.2:p.Arg110Gly | | 27 |
| 9 | 35792818 | ['G', 'A'] | 1 | 1.00 | ['missense_variant'] | MODERATE | NPR2 | ENST00000342694.6:c.410G>A | ENSP00000341083.2:p.Arg137His | | 21 |
| 9 | 35792833 | ['C', 'T'] | 1 | 1.00 | ['missense_variant'] | MODERATE | NPR2 | ENST00000342694.6:c.425C>T | ENSP00000341083.2:p.Thr142Ile | | 21 |
| 9 | 35792839 | ['C', 'T'] | 1 | 1.00 | ['missense_variant'] | MODERATE | NPR2 | ENST00000342694.6:c.431C>T | ENSP00000341083.2:p.Pro144Leu | | 24 |
| 9 | 35792883 | ['C', 'T'] | 1 | 1.00 | ['missense_variant'] | MODERATE | NPR2 | ENST00000342694.6:c.475C>T | ENSP00000341083.2:p.His159Tyr | | 22 |
| 9 | 35792885 | ['C', 'G'] | 2 | 1.00 | ['missense_variant'] | MODERATE | NPR2 | ENST00000342694.6:c.477C>G | ENSP00000341083.2:p.His159Gln | | 19 |
| 9 | 35792890 | ['A', 'G'] | 5 | 1.00 | ['missense_variant'] | MODERATE | NPR2 | ENST00000342694.6:c.482A>G | ENSP00000341083.2:p.Asn161Ser | 1.59E-05 | 23 |
| 9 | 35792899 | ['C', 'T'] | 1 | 1.00 | ['missense_variant'] | MODERATE | NPR2 | ENST00000342694.6:c.491C>T | ENSP00000341083.2:p.Ala164Val | | 17 |
| 9 | 35792901 | ['C', 'T'] | 2 | 1.00 | ['missense_variant'] | MODERATE | NPR2 | ENST00000342694.6:c.493C>T | ENSP00000341083.2:p.Arg165Cys | 3.98E-06 | 26 |
| 9 | 35792928 | ['C', 'T'] | 1 | 1.00 | ['missense_variant'] | MODERATE | NPR2 | ENST00000342694.6:c.520C>T | ENSP00000341083.2:p.Arg174Cys | 3.98E-06 | 27 |
| 9 | 35792992 | ['A', 'G'] | 1 | 1.00 | ['missense_variant'] | MODERATE | NPR2 | ENST00000342694.6:c.584A>G | ENSP00000341083.2:p.Asn195Ser | 7.99E-06 | 23 |
| 9 | 35793021 | ['C', 'T'] | 1 | 1.00 | ['stop_gained'] | HIGH | NPR2 | ENST00000342694.6:c.613C>T | ENSP00000341083.2:p.Arg205Ter | | 35 |
| 9 | 35793022 | ['G', 'A'] | 1 | 1.00 | ['missense_variant'] | MODERATE | NPR2 | ENST00000342694.6:c.614G>A | ENSP00000341083.2:p.Arg205Gln | | 20 |
| 9 | 35793033 | ['G', 'A'] | 2 | 1.00 | ['missense_variant'] | MODERATE | NPR2 | ENST00000342694.6:c.625G>A | ENSP00000341083.2:p.Gly209Ser | 4.09E-06 | 21 |
| 9 | 35793039 | ['G', 'A'] | 3 | 1.00 | ['missense_variant'] | MODERATE | NPR2 | ENST00000342694.6:c.631G>A | ENSP00000341083.2:p.Glu211Lys | 4.13E-06 | 21 |
| 9 | 35793904 | ['A', 'C'] | 1 | 1.00 | ['missense_variant'] | MODERATE | NPR2 | ENST00000342694.6:c.674A>C | ENSP00000341083.2:p.Tyr225Ser | | 24 |
| 9 | 35793906 | ['A', 'G'] | 1 | 1.00 | ['missense_variant'] | MODERATE | NPR2 | ENST00000342694.6:c.676A>G | ENSP00000341083.2:p.Ile226Val | 1.19E-05 | 21 |
| 9 | 35793952 | ['A', 'T'] | 1 | 1.00 | ['missense_variant'] | MODERATE | NPR2 | ENST00000342694.6:c.722A>T | ENSP00000341083.2:p.Gln241Leu | | 21 |
| 9 | 35793955 | ['G', 'A'] | 6 | 1.00 | ['missense_variant'] | MODERATE | NPR2 | ENST00000342694.6:c.725G>A | ENSP00000341083.2:p.Arg242Lys | 2.78E-05 | 21 |
| 9 | 35793980 | ['T', 'A'] | 1 | 1.00 | ['stop_gained'] | HIGH | NPR2 | ENST00000342694.6:c.750T>A | ENSP00000341083.2:p.Tyr250Ter | | 35 |
| 9 | 35794018 | ['G', 'A'] | 2 | 1.00 | ['missense_variant'] | MODERATE | NPR2 | ENST00000342694.6:c.788G>A | ENSP00000341083.2:p.Arg263His | 3.58E-05 | 24 |
| 9 | 35794024 | ['G', 'A'] | 1 | 1.00 | ['missense_variant'] | MODERATE | NPR2 | ENST00000342694.6:c.794G>A | ENSP00000341083.2:p.Gly265Asp | | 20 |
| 9 | 35794029 | ['A', 'G'] | 1 | 1.00 | ['missense_variant'] | MODERATE | NPR2 | ENST00000342694.6:c.799A>G | ENSP00000341083.2:p.Thr267Ala | 2.39E-05 | 21 |
| 9 | 35794044 | ['C', 'T'] | 2 | 1.00 | ['missense_variant'] | MODERATE | NPR2 | ENST00000342694.6:c.814C>T | ENSP00000341083.2:p.Arg272Trp | 2.78E-05 | 24 |
| 9 | 35794045 | ['G', 'A'] | 1 | 1.00 | ['missense_variant'] | MODERATE | NPR2 | ENST00000342694.6:c.815G>A | ENSP00000341083.2:p.Arg272Gln | 1.19E-05 | 20 |
| 9 | 35794056 | ['G', 'C'] | 1 | 1.00 | ['missense_variant'] | MODERATE | NPR2 | ENST00000342694.6:c.826G>C | ENSP00000341083.2:p.Asp276His | 3.98E-06 | 24 |
| 9 | 35794060 | ['A', 'G'] | 2 | 1.00 | ['missense_variant'] | MODERATE | NPR2 | ENST00000342694.6:c.830A>G | ENSP00000341083.2:p.Asn277Ser | 4.38E-05 | 21 |
| 9 | 35794063 | ['G', 'A'] | 1 | 1.00 | ['missense_variant'] | MODERATE | NPR2 | ENST00000342694.6:c.833G>A | ENSP00000341083.2:p.Arg278His | 2.79E-05 | 22 |
| 9 | 35794068 | ['C', 'T'] | 3 | 1.00 | ['missense_variant'] | MODERATE | NPR2 | ENST00000342694.6:c.838C>T | ENSP00000341083.2:p.Arg280Trp | 1.20E-05 | 24 |
| 9 | 35794093 | ['A', 'G'] | 1 | 1.00 | ['missense_variant'] | MODERATE | NPR2 | ENST00000342694.6:c.863A>G | ENSP00000341083.2:p.Glu288Gly | 1.20E-05 | 25 |
| 9 | 35799630 | ['A', 'G'] | 1 | 1.00 | ['missense_variant'] | MODERATE | NPR2 | ENST00000342694.6:c.886A>G | ENSP00000341083.2:p.Ile296Val | | 19 |
| 9 | 35799634 | ['C', 'T'] | 2 | 1.00 | ['missense_variant'] | MODERATE | NPR2 | ENST00000342694.6:c.890C>T | ENSP00000341083.2:p.Thr297Met | 3.98E-06 | 26 |
| 9 | 35799645 | ['C', 'T'] | 1 | 1.00 | ['missense_variant'] | MODERATE | NPR2 | ENST00000342694.6:c.901C>T | ENSP00000341083.2:p.Pro301Ser | | 25 |
| 9 | 35799660 | ['T', 'G'] | 1 | 1.00 | ['missense_variant'] | MODERATE | NPR2 | ENST00000342694.6:c.916T>G | ENSP00000341083.2:p.Tyr306Asp | | 28 |
| 9 | 35799679 | ['G', 'C'] | 1 | 1.00 | ['missense_variant'] | MODERATE | NPR2 | ENST00000342694.6:c.935G>C | ENSP00000341083.2:p.Arg312Pro | | 24 |
| 9 | 35799697 | ['G', 'A'] | 7 | 1.00 | ['missense_variant'] | MODERATE | NPR2 | ENST00000342694.6:c.953G>A | ENSP00000341083.2:p.Arg318Gln | 7.95E-05 | 22 |
| 9 | 35799699 | ['G', 'C'] | 2 | 1.00 | ['missense_variant'] | MODERATE | NPR2 | ENST00000342694.6:c.955G>C | ENSP00000341083.2:p.Glu319Gln | | 17 |
| 9 | 35799714 | ['G', 'C'] | 1 | 1.00 | ['missense_variant'] | MODERATE | NPR2 | ENST00000342694.6:c.970G>C | ENSP00000341083.2:p.Glu324Gln | 1.19E-05 | 22 |
| 9 | 35799715 | ['A', 'T'] | 1 | 1.00 | ['missense_variant'] | MODERATE | NPR2 | ENST00000342694.6:c.971A>T | ENSP00000341083.2:p.Glu324Val | | 23 |
| 9 | 35800041 | ['G', 'GCTT | 1 | 1.00 | ['inframe_insertion'] | MODERATE | NPR2 | ENST00000342694.6:c.1010_1012dup | ENSP00000341083.2:p.Phe337dup | 3.98E-06 | |
| 9 | 35800104 | ['C', 'T'] | 1 | 1.00 | ['missense_variant'] | MODERATE | NPR2 | ENST00000342694.6:c.1070C>T | ENSP00000341083.2:p.Thr357Ile | 5.17E-05 | 21 |
| 9 | 35800110 | ['A', 'C'] | 2 | 1.00 | ['missense_variant'] | MODERATE | NPR2 | ENST00000342694.6:c.1076A>C | ENSP00000341083.2:p.Glu359Ala | | 19 |
| 9 | 35800122 | ['G', 'A'] | 4 | 1.00 | ['missense_variant'] | MODERATE | NPR2 | ENST00000342694.6:c.1088G>A | ENSP00000341083.2:p.Arg363Gln | 5.57E-05 | 22 |
| 9 | 35800124 | ['AT', 'A'] | 3 | 1.00 | ['frameshift_variant'] | HIGH | NPR2 | ENST00000342694.6:c.1092del | ENSP00000341083.2:p.Ile364MetfsTer13 | 7.95E-06 | |
| 9 | 35800133 | ['A', 'G'] | 1 | 1.00 | ['missense_variant'] | MODERATE | NPR2 | ENST00000342694.6:c.1099A>G | ENSP00000341083.2:p.Lys367Glu | | 24 |
| 9 | 35800142 | ['G', 'A'] | 2 | 1.00 | ['missense_variant'] | MODERATE | NPR2 | ENST00000342694.6:c.1108G>A | ENSP00000341083.2:p.Gly370Arg | 1.19E-05 | 24 |
| 9 | 35800427 | ['C', 'T'] | 2 | 1.00 | ['stop_gained'] | HIGH | NPR2 | ENST00000342694.6:c.1162C>T | ENSP00000341083.2:p.Arg388Ter | 1.19E-05 | 35 |
| 9 | 35800720 | ['C', 'G'] | 1 | 1.00 | ['missense_variant'] | MODERATE | NPR2 | ENST00000342694.6:c.1230C>G | ENSP00000341083.2:p.His410Gln | | 18 |
| 9 | 35800733 | ['G', 'A'] | 4 | 1.00 | ['missense_variant'] | MODERATE | NPR2 | ENST00000342694.6:c.1243G>A | ENSP00000341083.2:p.Glu415Lys | | 21 |
| 9 | 35800752 | ['C', 'T'] | 7 | 1.00 | ['missense_variant'] | MODERATE | NPR2 | ENST00000342694.6:c.1262C>T | ENSP00000341083.2:p.Thr421Met | 7.16E-05 | 21 |
| 9 | 35800757 | ['C', 'T'] | 6 | 1.00 | ['missense_variant'] | MODERATE | NPR2 | ENST00000342694.6:c.1267C>T | ENSP00000341083.2:p.Arg423Trp | 2.39E-05 | 24 |
| 9 | 35800758 | ['G', 'A'] | 1 | 1.00 | ['missense_variant'] | MODERATE | NPR2 | ENST00000342694.6:c.1268G>A | ENSP00000341083.2:p.Arg423Gln | 1.19E-05 | 22 |
| 9 | 35800782 | ['C', 'T'] | 2 | 1.00 | ['missense_variant'] | MODERATE | NPR2 | ENST00000342694.6:c.1292C>T | ENSP00000341083.2:p.Ala431Val | | 12 |
| 9 | 35800791 | ['C', 'T'] | 2 | 1.00 | ['missense_variant'] | MODERATE | NPR2 | ENST00000342694.6:c.1301C>T | ENSP00000341083.2:p.Ser434Leu | 1.99E-05 | 15 |
| 9 | 35800803 | ['C', 'A'] | 1 | 1.00 | ['missense_variant'] | MODERATE | NPR2 | ENST00000342694.6:c.1313C>A | ENSP00000341083.2:p.Pro438His | | 23 |
| 9 | 35800814 | ['G', 'A'] | 2 | 1.00 | ['missense_variant'] | MODERATE | NPR2 | ENST00000342694.6:c.1324G>A | ENSP00000341083.2:p.Asp442Asn | 3.98E-06 | 23 |
| 9 | 35800815 | ['A', 'G'] | 1 | 1.00 | ['missense_variant'] | MODERATE | NPR2 | ENST00000342694.6:c.1325A>G | ENSP00000341083.2:p.Asp442Gly | 3.98E-06 | 24 |
| 9 | 35801090 | ['A', 'G'] | 1 | 1.00 | ['missense_variant'] | MODERATE | NPR2 | ENST00000342694.6:c.1372A>G | ENSP00000341083.2:p.Ile458Val | 2.39E-05 | 21 |
| 9 | 35801091 | ['T', 'C'] | 1 | 1.00 | ['missense_variant'] | MODERATE | NPR2 | ENST00000342694.6:c.1373T>C | ENSP00000341083.2:p.Ile458Thr | 7.95E-06 | 24 |
| 9 | 35801118 | ['T', 'G'] | 1 | 1.00 | ['missense_variant'] | MODERATE | NPR2 | ENST00000342694.6:c.1400T>G | ENSP00000341083.2:p.Phe467Cys | | 25 |
| 9 | 35801125 | ['G', 'A'] | 3 | 1.00 | ['missense_variant'] | MODERATE | NPR2 | ENST00000342694.6:c.1407G>A | ENSP00000341083.2:p.Met469Ile | 1.99E-05 | 16 |
| 9 | 35801154 | ['G', 'A'] | 1 | 1.00 | _variant', 'splice_regi | MODERATE | NPR2 | ENST00000342694.6:c.1436G>A | ENSP00000341083.2:p.Arg479Gln | 7.96E-06 | 35 |
| 9 | 35801646 | ['G', 'C'] | 3 | 1.00 | ['missense_variant'] | MODERATE | NPR2 | ENST00000342694.6:c.1440G>C | ENSP00000341083.2:p.Lys480Asn | 1.59E-05 | 23 |

Supplementary Table 5 Genetic variants found in five height genes

| chr | position | alleles | AC | nt_qc.call_ | vep.consequence | vep.impact | vep.symbol | vep.hgvsc | vep.hgvsp | vep.gnoma | vep.cadd_phred |
|---|---|---|---|---|---|---|---|---|---|---|---|
| 9 | 35801684 | ['G', 'A'] | 4 | 1.00 | ['missense_variant'] | MODERATE | NPR2 | ENST00000342694.6:c.1478G>A | ENSP00000341083.2:p.Arg493His | 7.96E-06 | 33 |
| 9 | 35801722 | ['C', 'T'] | 1 | 1.00 | ['missense_variant'] | MODERATE | NPR2 | ENST00000342694.6:c.1516C>T | ENSP00000341083.2:p.Arg506Cys | 1.59E-05 | 24 |
| 9 | 35801723 | ['G', 'A'] | 2 | 1.00 | ['missense_variant'] | MODERATE | NPR2 | ENST00000342694.6:c.1517G>A | ENSP00000341083.2:p.Arg506His | 0.000119 | 28 |
| 9 | 35801728 | ['C', 'G'] | 2 | 1.00 | ['missense_variant'] | MODERATE | NPR2 | ENST00000342694.6:c.1522C>G | ENSP00000341083.2:p.His508Asp | 3.98E-06 | 22 |
| 9 | 35801744 | ['G', 'A'] | 1 | 1.00 | ['missense_variant'] | MODERATE | NPR2 | ENST00000342694.6:c.1538G>A | ENSP00000341083.2:p.Ser513Asn | | 29 |
| 9 | 35801759 | ['C', 'T'] | 2 | 1.00 | ['missense_variant'] | MODERATE | NPR2 | ENST00000342694.6:c.1553C>T | ENSP00000341083.2:p.Ser518Leu | 7.95E-06 | 25 |
| 9 | 35801932 | ['T', 'C'] | 1 | 1.00 | ['missense_variant'] | MODERATE | NPR2 | ENST00000342694.6:c.1564T>C | ENSP00000341083.2:p.Ser522Pro | | 31 |
| 9 | 35801941 | ['G', 'A'] | 1 | 1.00 | ['missense_variant'] | MODERATE | NPR2 | ENST00000342694.6:c.1573G>A | ENSP00000341083.2:p.Gly525Ser | | 24 |
| 9 | 35801989 | ['G', 'A'] | 2 | 1.00 | ['missense_variant'] | MODERATE | NPR2 | ENST00000342694.6:c.1621G>A | ENSP00000341083.2:p.Gly541Ser | 0.000151 | 32 |
| 9 | 35802209 | ['A', 'T'] | 6 | 1.00 | ['missense_variant'] | MODERATE | NPR2 | ENST00000342694.6:c.1636A>T | ENSP00000341083.2:p.Asn546Tyr | 3.18E-05 | 27 |
| 9 | 35802218 | ['G', 'A'] | 2 | 1.00 | ['missense_variant'] | MODERATE | NPR2 | ENST00000342694.6:c.1645G>A | ENSP00000341083.2:p.Ala549Thr | 1.19E-05 | 30 |
| 9 | 35802230 | ['G', 'C'] | 1 | 1.00 | ['missense_variant'] | MODERATE | NPR2 | ENST00000342694.6:c.1657G>C | ENSP00000341083.2:p.Val553Leu | | 22 |
| 9 | 35802246 | ['T', 'C'] | 1 | 1.00 | ['missense_variant'] | MODERATE | NPR2 | ENST00000342694.6:c.1673T>C | ENSP00000341083.2:p.Ile558Thr | 7.97E-06 | 27 |
| 9 | 35802249 | ['A', 'G'] | 1 | 1.00 | ['missense_variant'] | MODERATE | NPR2 | ENST00000342694.6:c.1676A>G | ENSP00000341083.2:p.Glu559Gly | | 29 |
| 9 | 35802263 | ['G', 'C'] | 1 | 1.00 | ['missense_variant'] | MODERATE | NPR2 | ENST00000342694.6:c.1690G>C | ENSP00000341083.2:p.Val564Leu | | 23 |
| 9 | 35802545 | ['G', 'A'] | 1 | 1.00 | ['missense_variant'] | MODERATE | NPR2 | ENST00000342694.6:c.1753G>A | ENSP00000341083.2:p.Ala585Thr | 3.98E-06 | 28 |
| 9 | 35802609 | ['T', 'G'] | 1 | 1.00 | splice_donor_variant' | HIGH | NPR2 | ENST00000342694.6:c.1815+2T>G | | | 34 |
| 9 | 35802749 | ['C', 'G'] | 1 | 1.00 | ['missense_variant'] | MODERATE | NPR2 | ENST00000342694.6:c.1833C>G | ENSP00000341083.2:p.Asp611Glu | | 18 |
| 9 | 35802761 | ['G', 'C'] | 2 | 1.00 | ['missense_variant'] | MODERATE | NPR2 | ENST00000342694.6:c.1845G>C | ENSP00000341083.2:p.Leu615Phe | | 27 |
| 9 | 35802774 | ['C', 'T'] | 4 | 1.00 | ['missense_variant'] | MODERATE | NPR2 | ENST00000342694.6:c.1858C>T | ENSP00000341083.2:p.Arg620Cys | 3.98E-06 | 28 |
| 9 | 35802802 | ['A', 'G'] | 1 | 1.00 | _variant', 'splice_regi | MODERATE | NPR2 | ENST00000342694.6:c.1886A>G | ENSP00000341083.2:p.Lys629Arg | | 28 |
| 9 | 35805565 | ['T', 'G'] | 1 | 1.00 | ['missense_variant'] | MODERATE | NPR2 | ENST00000342694.6:c.1942T>G | ENSP00000341083.2:p.Ser648Ala | | 26 |
| 9 | 35805638 | ['C', 'T'] | 1 | 1.00 | ['missense_variant'] | MODERATE | NPR2 | ENST00000342694.6:c.2015C>T | ENSP00000341083.2:p.Ala672Val | | 16 |
| 9 | 35805648 | ['T', 'A'] | 2 | 1.00 | ['missense_variant'] | MODERATE | NPR2 | ENST00000342694.6:c.2025T>A | ENSP00000341083.2:p.Asp675Glu | | 17 |
| 9 | 35805656 | ['A', 'G'] | 1 | 1.00 | ['missense_variant'] | MODERATE | NPR2 | ENST00000342694.6:c.2033A>G | ENSP00000341083.2:p.His678Arg | | 24 |
| 9 | 35805661 | ['C', 'T'] | 1 | 1.00 | ['missense_variant'] | MODERATE | NPR2 | ENST00000342694.6:c.2038C>T | ENSP00000341083.2:p.Leu680Phe | 3.98E-06 | 21 |
| 9 | 35805664 | ['T', 'C'] | 1 | 1.00 | ['missense_variant'] | MODERATE | NPR2 | ENST00000342694.6:c.2041T>C | ENSP00000341083.2:p.Tyr681His | | 25 |
| 9 | 35805833 | ['A', 'C'] | 1 | 1.00 | ['missense_variant'] | MODERATE | NPR2 | ENST00000342694.6:c.2051A>C | ENSP00000341083.2:p.Lys684Thr | | 27 |
| 9 | 35805844 | ['G', 'T'] | 3 | 1.00 | ['missense_variant'] | MODERATE | NPR2 | ENST00000342694.6:c.2062G>T | ENSP00000341083.2:p.Ala688Ser | 7.96E-06 | 25 |
| 9 | 35805869 | ['C', 'T'] | 1 | 1.00 | ['missense_variant'] | MODERATE | NPR2 | ENST00000342694.6:c.2087C>T | ENSP00000341083.2:p.Pro696Leu | 3.98E-06 | 16 |
| 9 | 35805873 | ['G', 'T'] | 1 | 1.00 | ['missense_variant'] | MODERATE | NPR2 | ENST00000342694.6:c.2091G>T | ENSP00000341083.2:p.Leu697Phe | 3.98E-06 | 19 |
| 9 | 35805895 | ['G', 'A'] | 1 | 1.00 | ['missense_variant'] | MODERATE | NPR2 | ENST00000342694.6:c.2113G>A | ENSP00000341083.2:p.Ala705Thr | 3.98E-06 | 28 |
| 9 | 35805901 | ['G', 'A'] | 1 | 1.00 | ['missense_variant'] | MODERATE | NPR2 | ENST00000342694.6:c.2119G>A | ENSP00000341083.2:p.Val707Ile | 1.99E-05 | 22 |
| 9 | 35805932 | ['T', 'C'] | 1 | 1.00 | ['missense_variant'] | MODERATE | NPR2 | ENST00000342694.6:c.2150T>C | ENSP00000341083.2:p.Ile717Thr | 1.20E-05 | 28 |
| 9 | 35805941 | ['G', 'A'] | 1 | 1.00 | ['missense_variant'] | MODERATE | NPR2 | ENST00000342694.6:c.2159G>A | ENSP00000341083.2:p.Arg720His | 2.39E-05 | 32 |
| 9 | 35805945 | ['T', 'A'] | 3 | 1.00 | ['missense_variant'] | MODERATE | NPR2 | ENST00000342694.6:c.2163T>A | ENSP00000341083.2:p.Ser721Arg | 7.97E-06 | 18 |
| 9 | 35805961 | ['G', 'C'] | 1 | 1.00 | ['missense_variant'] | MODERATE | NPR2 | ENST00000342694.6:c.2179G>C | ENSP00000341083.2:p.Glu727Gln | | 24 |
| 9 | 35806069 | ['T', 'G'] | 1 | 1.00 | ['missense_variant'] | MODERATE | NPR2 | ENST00000342694.6:c.2208T>G | ENSP00000341083.2:p.Ile736Met | 7.95E-06 | 24 |
| 9 | 35806090 | ['TCAG', 'T | 1 | 1.00 | ['inframe_deletion'] | MODERATE | NPR2 | ENST00000342694.6:c.2231_2233del | ENSP00000341083.2:p.Gln744del | | |
| 9 | 35806095 | ['G', 'A'] | 3 | 1.00 | ['missense_variant'] | MODERATE | NPR2 | ENST00000342694.6:c.2234G>A | ENSP00000341083.2:p.Arg745Gln | 2.39E-05 | 22 |
| 9 | 35806103 | ['T', 'A'] | 1 | 1.00 | ['missense_variant'] | MODERATE | NPR2 | ENST00000342694.6:c.2242T>A | ENSP00000341083.2:p.Phe748Ile | | 27 |
| 9 | 35806121 | ['C', 'T'] | 3 | 1.00 | ['missense_variant'] | MODERATE | NPR2 | ENST00000342694.6:c.2260C>T | ENSP00000341083.2:p.Arg754Trp | 7.95E-05 | 24 |
| 9 | 35806129 | ['A', 'C'] | 2 | 1.00 | ['missense_variant'] | MODERATE | NPR2 | ENST00000342694.6:c.2268A>C | ENSP00000341083.2:p.Gln756His | | 15 |
| 9 | 35806157 | ['G', 'C'] | 4 | 1.00 | ['missense_variant'] | MODERATE | NPR2 | ENST00000342694.6:c.2296G>C | ENSP00000341083.2:p.Glu766Gln | 7.95E-06 | 21 |
| 9 | 35806161 | ['G', 'A'] | 1 | 1.00 | ['missense_variant'] | MODERATE | NPR2 | ENST00000342694.6:c.2300G>A | ENSP00000341083.2:p.Arg767Gln | 1.19E-05 | 26 |
| 9 | 35806187 | ['C', 'T'] | 1 | 1.00 | ['missense_variant'] | MODERATE | NPR2 | ENST00000342694.6:c.2326C>T | ENSP00000341083.2:p.Arg776Trp | 7.96E-06 | 26 |
| 9 | 35806220 | ['C', 'T'] | 1 | 1.00 | ['missense_variant'] | MODERATE | NPR2 | ENST00000342694.6:c.2359C>T | ENSP00000341083.2:p.Arg787Trp | 0.000291 | 28 |
| 9 | 35806221 | ['G', 'C'] | 2 | 1.00 | ['missense_variant'] | MODERATE | NPR2 | ENST00000342694.6:c.2360G>C | ENSP00000341083.2:p.Arg787Pro | | 28 |
| 9 | 35806221 | ['G', 'A'] | 1 | 1.00 | ['missense_variant'] | MODERATE | NPR2 | ENST00000342694.6:c.2360G>A | ENSP00000341083.2:p.Arg787Gln | 3.98E-06 | 27 |
| 9 | 35806223 | ['C', 'T'] | 1 | 1.00 | ['missense_variant'] | MODERATE | NPR2 | ENST00000342694.6:c.2362C>T | ENSP00000341083.2:p.Arg788Cys | 2.79E-05 | 27 |
| 9 | 35806429 | ['C', 'T'] | 1 | 1.00 | ['missense_variant'] | MODERATE | NPR2 | ENST00000342694.6:c.2410C>T | ENSP00000341083.2:p.Arg804Cys | 3.98E-06 | 28 |
| 9 | 35806448 | ['A', 'T'] | 1 | 1.00 | ['missense_variant'] | MODERATE | NPR2 | ENST00000342694.6:c.2429A>T | ENSP00000341083.2:p.Asn810Ile | | 25 |
| 9 | 35806474 | ['C', 'T'] | 1 | 1.00 | ['missense_variant'] | MODERATE | NPR2 | ENST00000342694.6:c.2455C>T | ENSP00000341083.2:p.Arg819Cys | 1.99E-05 | 27 |
| 9 | 35806501 | ['C', 'T'] | 1 | 1.00 | ['missense_variant'] | MODERATE | NPR2 | ENST00000342694.6:c.2482C>T | ENSP00000341083.2:p.Arg828Cys | | 29 |
| 9 | 35806506 | ['G', 'C'] | 1 | 1.00 | ['missense_variant'] | MODERATE | NPR2 | ENST00000342694.6:c.2487G>C | ENSP00000341083.2:p.Lys829Asn | | 25 |
| 9 | 35807045 | ['C', 'T'] | 1 | 1.00 | ['missense_variant'] | MODERATE | NPR2 | ENST00000342694.6:c.2542C>T | ENSP00000341083.2:p.Arg848Trp | 1.59E-05 | 24 |
| 9 | 35807053 | ['G', 'C'] | 3 | 1.00 | ['missense_variant'] | MODERATE | NPR2 | ENST00000342694.6:c.2550G>C | ENSP00000341083.2:p.Glu850Asp | 3.98E-06 | 24 |
| 9 | 35807085 | ['C', 'T'] | 1 | 1.00 | ['missense_variant'] | MODERATE | NPR2 | ENST00000342694.6:c.2582C>T | ENSP00000341083.2:p.Thr861Ile | | 27 |
| 9 | 35807129 | ['G', 'A'] | 1 | 1.00 | ['missense_variant'] | MODERATE | NPR2 | ENST00000342694.6:c.2626G>A | ENSP00000341083.2:p.Glu876Lys | | 31 |
| 9 | 35807144 | ['C', 'G'] | 2 | 1.00 | _variant', 'splice_regi | MODERATE | NPR2 | ENST00000342694.6:c.2641C>G | ENSP00000341083.2:p.Gln881Glu | | 23 |
| 9 | 35808509 | ['G', 'A'] | 1 | 1.00 | _variant', 'splice_regi | MODERATE | NPR2 | ENST00000342694.6:c.2713G>A | ENSP00000341083.2:p.Val905Met | 4.02E-06 | 35 |
| 9 | 35808519 | ['T', 'C'] | 3 | 1.00 | ['missense_variant'] | MODERATE | NPR2 | ENST00000342694.6:c.2723T>C | ENSP00000341083.2:p.Ile908Thr | 6.42E-05 | 29 |
| 9 | 35808545 | ['G', 'C'] | 1 | 1.00 | ['missense_variant'] | MODERATE | NPR2 | ENST00000342694.6:c.2749G>C | ENSP00000341083.2:p.Gly917Arg | | 32 |
| 9 | 35808558 | ['G', 'A'] | 1 | 1.00 | ['missense_variant'] | MODERATE | NPR2 | ENST00000342694.6:c.2762G>A | ENSP00000341083.2:p.Arg921Gln | 1.20E-05 | 33 |
| 9 | 35808591 | ['G', 'A'] | 7 | 1.00 | ['missense_variant'] | MODERATE | NPR2 | ENST00000342694.6:c.2795G>A | ENSP00000341083.2:p.Arg932His | 3.19E-05 | 29 |
| 9 | 35808608 | ['C', 'G'] | 1 | 1.00 | ['missense_variant'] | MODERATE | NPR2 | ENST00000342694.6:c.2812C>G | ENSP00000341083.2:p.Leu938Val | | 24 |
| 9 | 35808613 | ['T', 'A'] | 1 | 1.00 | ['missense_variant'] | MODERATE | NPR2 | ENST00000342694.6:c.2817T>A | ENSP00000341083.2:p.Asp939Glu | | 17 |
| 9 | 35808621 | ['C', 'A'] | 1 | 1.00 | ['missense_variant'] | MODERATE | NPR2 | ENST00000342694.6:c.2825C>A | ENSP00000341083.2:p.Ser942Tyr | | 18 |
| 9 | 35808629 | ['C', 'T'] | 1 | 1.00 | ['missense_variant'] | MODERATE | NPR2 | ENST00000342694.6:c.2833C>T | ENSP00000341083.2:p.Arg945Cys | 3.18E-05 | 25 |

Supplementary Table 5 Genetic variants found in five height genes

| chr | position | alleles | AC | int_qc.call_ | vep.consequence | vep.impact | vep.symbol | vep.hgvsc | vep.hgvsp | vep.gnoma | vep.cadd_phred |
|---|---|---|---|---|---|---|---|---|---|---|---|
| 9 | 35808644 | ['C', 'A'] | 1 | 1.00 | ['missense_variant'] | MODERATE | NPR2 | ENST00000342694.6:c.2848C>A | ENSP00000341083.2:p.Pro950Thr | | 26 |
| 9 | 35808663 | ['T', 'A'] | 3 | 1.00 | ['missense_variant'] | MODERATE | NPR2 | ENST00000342694.6:c.2867T>A | ENSP00000341083.2:p.Leu956Gln | 1.99E-05 | 32 |
| 9 | 35808665 | ['C', 'T'] | 1 | 1.00 | ['missense_variant'] | MODERATE | NPR2 | ENST00000342694.6:c.2869C>T | ENSP00000341083.2:p.Arg957Cys | 1.19E-05 | 31 |
| 9 | 35808669 | ['T', 'C'] | 1 | 1.00 | ['missense_variant'] | MODERATE | NPR2 | ENST00000342694.6:c.2873T>C | ENSP00000341083.2:p.Ile958Thr | 7.95E-06 | 29 |
| 9 | 35809177 | ['C', 'A'] | 1 | 1.00 | ['missense_variant'] | MODERATE | NPR2 | ENST00000342694.6:c.3008C>A | ENSP00000341083.2:p.Ser1003Tyr | | 25 |
| 9 | 35809180 | ['C', 'T'] | 4 | 1.00 | ['missense_variant'] | MODERATE | NPR2 | ENST00000342694.6:c.3011C>T | ENSP00000341083.2:p.Thr1004Ile | 2.39E-05 | 24 |
| 9 | 35809186 | ['A', 'C'] | 1 | 1.00 | ['missense_variant'] | MODERATE | NPR2 | ENST00000342694.6:c.3017A>C | ENSP00000341083.2:p.Lys1006Thr | 3.98E-06 | 27 |
| 9 | 35809194 | ['C', 'G'] | 2 | 1.00 | ['missense_variant'] | MODERATE | NPR2 | ENST00000342694.6:c.3025C>G | ENSP00000341083.2:p.Leu1009Val | | 24 |
| 9 | 35809206 | ['G', 'A'] | 1 | 1.00 | ['missense_variant'] | MODERATE | NPR2 | ENST00000342694.6:c.3037G>A | ENSP00000341083.2:p.Gly1013Arg | | 28 |
| 9 | 35809227 | ['CGG', 'C' | 1 | 1.00 | ['frameshift_variant'] | HIGH | NPR2 | ENST00000342694.6:c.3063_3064del | ENSP00000341083.2:p.Asp1022CysfsTer35 | | |
| 9 | 35809244 | ['G', 'A'] | 1 | 1.00 | ['missense_variant'] | MODERATE | NPR2 | ENST00000342694.6:c.3075G>A | ENSP00000341083.2:p.Met1025Ile | 1.19E-05 | 24 |
| 9 | 35809395 | ['C', 'T'] | 2 | 1.00 | ['stop_gained'] | HIGH | NPR2 | ENST00000342694.6:c.3094C>T | ENSP00000341083.2:p.Arg1032Ter | | 40 |
| 9 | 35809396 | ['G', 'A'] | 3 | 1.00 | ['missense_variant'] | MODERATE | NPR2 | ENST00000342694.6:c.3095G>A | ENSP00000341083.2:p.Arg1032Gln | 3.18E-05 | 24 |
| 9 | 35809413 | ['G', 'A'] | 1 | 1.00 | ['missense_variant'] | MODERATE | NPR2 | ENST00000342694.6:c.3112G>A | ENSP00000341083.2:p.Gly1038Arg | | 29 |
| 9 | 35809419 | ['C', 'T'] | 7 | 1.00 | ['missense_variant'] | MODERATE | NPR2 | ENST00000342694.6:c.3118C>T | ENSP00000341083.2:p.Arg1040Trp | 3.18E-05 | 25 |
| 9 | 35809420 | ['G', 'A'] | 6 | 1.00 | ['missense_variant'] | MODERATE | NPR2 | ENST00000342694.6:c.3119G>A | ENSP00000341083.2:p.Arg1040Gln | 7.56E-05 | 19 |
| 9 | 35809428 | ['C', 'T'] | 1 | 1.00 | ['missense_variant'] | MODERATE | NPR2 | ENST00000342694.6:c.3127C>T | ENSP00000341083.2:p.Pro1043Ser | | 18 |
| 9 | 35809434 | ['G', 'A'] | 1 | 1.00 | ['missense_variant'] | MODERATE | NPR2 | ENST00000342694.6:c.3133G>A | ENSP00000341083.2:p.Gly1045Arg | | 37 |
| 9 | 35809436 | ['AC', 'A'] | 1 | 1.00 | ['frameshift_variant'] | HIGH | NPR2 | ENST00000342694.6:c.3136del | ENSP00000341083.2:p.Leu1046SerfsTer? | | |
| 9 | 35809441 | ['T', 'C'] | 1 | 1.00 | ['missense_variant'] | MODERATE | NPR2 | ENST00000342694.6:c.3140T>C | ENSP00000341083.2:p.Leu1047Pro | | 25 |
| 15 | 98649583 | ['T', 'C'] | 1 | 1.00 | ['start_lost'] | HIGH | IGF1R | ENST00000650285.1:c.2T>C | ENSP00000497069.1:p.Met1? | | 23 |
| 15 | 98649597 | ['G', 'A'] | 3 | 1.00 | ['missense_variant'] | MODERATE | IGF1R | ENST00000650285.1:c.16G>A | ENSP00000497069.1:p.Gly6Arg | 0.000605 | 22 |
| 15 | 98649609 | ['C', 'T'] | 1 | 1.00 | ['missense_variant'] | MODERATE | IGF1R | ENST00000650285.1:c.28C>T | ENSP00000497069.1:p.Pro10Ser | 1.21E-05 | 21 |
| 15 | 98649610 | ['C', 'T'] | 1 | 1.00 | ['missense_variant'] | MODERATE | IGF1R | ENST00000650285.1:c.29C>T | ENSP00000497069.1:p.Pro10Leu | 4.05E-06 | 20 |
| 15 | 98649636 | ['C', 'G'] | 1 | 1.00 | ['missense_variant'] | MODERATE | IGF1R | ENST00000650285.1:c.55C>G | ENSP00000497069.1:p.Leu19Val | 1.21E-05 | 15 |
| 15 | 98649636 | ['C', 'T'] | 1 | 1.00 | ['missense_variant'] | MODERATE | IGF1R | ENST00000650285.1:c.55C>T | ENSP00000497069.1:p.Leu19Phe | 1.62E-05 | 15 |
| 15 | 98649640 | ['C', 'T'] | 1 | 1.00 | ['missense_variant'] | MODERATE | IGF1R | ENST00000650285.1:c.59C>T | ENSP00000497069.1:p.Ser20Phe | 3.24E-05 | 20 |
| 15 | 98649652 | ['C', 'G'] | 1 | 1.00 | ['missense_variant'] | MODERATE | IGF1R | ENST00000650285.1:c.71C>G | ENSP00000497069.1:p.Ser24Trp | 4.05E-06 | 21 |
| 15 | 98649654 | ['C', 'T'] | 1 | 1.00 | ['missense_variant'] | MODERATE | IGF1R | ENST00000650285.1:c.73C>T | ENSP00000497069.1:p.Leu25Phe | | 21 |
| 15 | 98649661 | ['C', 'T'] | 1 | 1.00 | ['missense_variant'] | MODERATE | IGF1R | ENST00000650285.1:c.80C>T | ENSP00000497069.1:p.Pro27Leu | 1.62E-05 | 23 |
| 15 | 98649674 | ['AAGTG', | 1 | 1.00 | 'coding_sequence_va | HIGH | IGF1R | ENST00000650285.1:c.94+3_94+6del | | 3.64E-05 | |
| 15 | 98707651 | ['C', 'A'] | 1 | 1.00 | ['missense_variant'] | MODERATE | IGF1R | ENST00000650285.1:c.184C>A | ENSP00000497069.1:p.Leu62Met | | 25 |
| 15 | 98707667 | ['C', 'G'] | 1 | 1.00 | ['missense_variant'] | MODERATE | IGF1R | ENST00000650285.1:c.200C>G | ENSP00000497069.1:p.Ala67Gly | | 19 |
| 15 | 98707679 | ['G', 'A'] | 2 | 1.00 | ['missense_variant'] | MODERATE | IGF1R | ENST00000650285.1:c.212G>A | ENSP00000497069.1:p.Arg71His | 2.39E-05 | 25 |
| 15 | 98707703 | ['C', 'T'] | 2 | 1.00 | ['missense_variant'] | MODERATE | IGF1R | ENST00000650285.1:c.236C>T | ENSP00000497069.1:p.Thr79Met | 1.59E-05 | 26 |
| 15 | 98707808 | ['A', 'G'] | 1 | 1.00 | ['missense_variant'] | MODERATE | IGF1R | ENST00000650285.1:c.341A>G | ENSP00000497069.1:p.Asn114Ser | | 23 |
| 15 | 98707838 | ['A', 'G'] | 1 | 1.00 | ['missense_variant'] | MODERATE | IGF1R | ENST00000650285.1:c.371A>G | ENSP00000497069.1:p.Asn124Ser | 1.59E-05 | 19 |
| 15 | 98707849 | ['A', 'G'] | 2 | 1.00 | ['missense_variant'] | MODERATE | IGF1R | ENST00000650285.1:c.382A>G | ENSP00000497069.1:p.Ile128Val | 3.98E-06 | 22 |
| 15 | 98707868 | ['G', 'A'] | 1 | 1.00 | ['missense_variant'] | MODERATE | IGF1R | ENST00000650285.1:c.401G>A | ENSP00000497069.1:p.Arg134Lys | 3.98E-06 | 21 |
| 15 | 98707958 | ['T', 'G'] | 1 | 1.00 | ['missense_variant'] | MODERATE | IGF1R | ENST00000650285.1:c.491T>G | ENSP00000497069.1:p.Val164Gly | | 22 |
| 15 | 98707976 | ['T', 'C'] | 2 | 1.00 | ['missense_variant'] | MODERATE | IGF1R | ENST00000650285.1:c.509T>C | ENSP00000497069.1:p.Val170Ala | 3.98E-06 | 23 |
| 15 | 98708024 | ['TGGA', 'T | 1 | 1.00 | ['inframe_deletion'] | MODERATE | IGF1R | ENST00000650285.1:c.562_564del | ENSP00000497069.1:p.Glu188del | | |
| 15 | 98708072 | ['A', 'G'] | 2 | 1.00 | ['missense_variant'] | MODERATE | IGF1R | ENST00000650285.1:c.605A>G | ENSP00000497069.1:p.Asn202Ser | 0.000137 | 17 |
| 15 | 98891348 | ['C', 'T'] | 1 | 1.00 | ['missense_variant'] | MODERATE | IGF1R | ENST00000650285.1:c.664C>T | ENSP00000497069.1:p.Arg222Trp | 0.00011 | 23 |
| 15 | 98891352 | ['C', 'T'] | 1 | 1.00 | ['missense_variant'] | MODERATE | IGF1R | ENST00000650285.1:c.668C>T | ENSP00000497069.1:p.Ala223Val | 4.05E-06 | 23 |
| 15 | 98891390 | ['C', 'G'] | 1 | 1.00 | ['missense_variant'] | MODERATE | IGF1R | ENST00000650285.1:c.706C>G | ENSP00000497069.1:p.Leu236Val | | 24 |
| 15 | 98891405 | ['G', 'A'] | 2 | 1.00 | ['missense_variant'] | MODERATE | IGF1R | ENST00000650285.1:c.721G>A | ENSP00000497069.1:p.Ala241Thr | 8.00E-06 | 16 |
| 15 | 98891417 | ['G', 'A'] | 3 | 1.00 | ['missense_variant'] | MODERATE | IGF1R | ENST00000650285.1:c.733G>A | ENSP00000497069.1:p.Asp245Asn | 4.00E-06 | 18 |
| 15 | 98891420 | ['A', 'G'] | 6 | 1.00 | ['missense_variant'] | MODERATE | IGF1R | ENST00000650285.1:c.736A>G | ENSP00000497069.1:p.Thr246Ala | 2.80E-05 | 17 |
| 15 | 98891439 | ['G', 'A'] | 4 | 1.00 | ['missense_variant'] | MODERATE | IGF1R | ENST00000650285.1:c.755G>A | ENSP00000497069.1:p.Arg252His | 2.79E-05 | 22 |
| 15 | 98891469 | ['C', 'G'] | 1 | 1.00 | ['missense_variant'] | MODERATE | IGF1R | ENST00000650285.1:c.785C>G | ENSP00000497069.1:p.Pro262Arg | | 15 |
| 15 | 98891478 | ['C', 'T'] | 1 | 1.00 | ['missense_variant'] | MODERATE | IGF1R | ENST00000650285.1:c.794C>T | ENSP00000497069.1:p.Pro265Leu | 7.96E-06 | 26 |
| 15 | 98891484 | ['A', 'C'] | 1 | 1.00 | ['missense_variant'] | MODERATE | IGF1R | ENST00000650285.1:c.800A>C | ENSP00000497069.1:p.Asn267Thr | 1.99E-05 | 16 |
| 15 | 98891540 | ['C', 'G'] | 1 | 1.00 | ['missense_variant'] | MODERATE | IGF1R | ENST00000650285.1:c.856C>G | ENSP00000497069.1:p.Leu286Val | | 13 |
| 15 | 98891564 | ['G', 'A'] | 1 | 1.00 | ['missense_variant'] | MODERATE | IGF1R | ENST00000650285.1:c.880G>A | ENSP00000497069.1:p.Glu294Lys | 8.09E-06 | 17 |
| 15 | 98891606 | ['C', 'T'] | 1 | 1.00 | ['missense_variant'] | MODERATE | IGF1R | ENST00000650285.1:c.922C>T | ENSP00000497069.1:p.Pro308Ser | | 26 |
| 15 | 98891622 | ['G', 'A'] | 1 | 1.00 | ['missense_variant'] | MODERATE | IGF1R | ENST00000650285.1:c.938G>A | ENSP00000497069.1:p.Arg313His | 4.18E-06 | 24 |
| 15 | 98891625 | ['A', 'G'] | 1 | 1.00 | ['missense_variant'] | MODERATE | IGF1R | ENST00000650285.1:c.941A>G | ENSP00000497069.1:p.Asn314Ser | | 18 |
| 15 | 98891628 | ['G', 'A'] | 1 | 1.00 | ['missense_variant'] | MODERATE | IGF1R | ENST00000650285.1:c.944G>A | ENSP00000497069.1:p.Gly315Asp | | 12 |
| 15 | 98891633 | ['C', 'G'] | 1 | 1.00 | ['missense_variant'] | MODERATE | IGF1R | ENST00000650285.1:c.949C>G | ENSP00000497069.1:p.Gln317Glu | | 16 |
| 15 | 98896760 | ['G', 'A'] | 1 | 1.00 | ['missense_variant'] | MODERATE | IGF1R | ENST00000650285.1:c.957G>A | ENSP00000497069.1:p.Met319Ile | | 23 |
| 15 | 98896789 | ['C', 'T'] | 1 | 1.00 | ['missense_variant'] | MODERATE | IGF1R | ENST00000650285.1:c.986C>T | ENSP00000497069.1:p.Pro329Leu | 7.96E-06 | 23 |
| 15 | 98896794 | ['G', 'A'] | 3 | 1.00 | ['missense_variant'] | MODERATE | IGF1R | ENST00000650285.1:c.991G>A | ENSP00000497069.1:p.Val331Ile | 1.99E-05 | 21 |
| 15 | 98896816 | ['C', 'G'] | 2 | 1.00 | ['missense_variant'] | MODERATE | IGF1R | ENST00000650285.1:c.1013C>G | ENSP00000497069.1:p.Thr338Arg | | 17 |
| 15 | 98896829 | ['T', 'G'] | 1 | 1.00 | ['missense_variant'] | MODERATE | IGF1R | ENST00000650285.1:c.1026T>G | ENSP00000497069.1:p.Asp342Glu | | 22 |
| 15 | 98896879 | ['A', 'G'] | 1 | 1.00 | ['missense_variant'] | MODERATE | IGF1R | ENST00000650285.1:c.1076A>G | ENSP00000497069.1:p.Asn359Ser | 1.59E-05 | 23 |
| 15 | 98896884 | ['C', 'T'] | 1 | 1.00 | ['missense_variant'] | MODERATE | IGF1R | ENST00000650285.1:c.1081C>T | ENSP00000497069.1:p.Leu361Phe | 3.98E-06 | 22 |
| 15 | 98896897 | ['G', 'A'] | 1 | 1.00 | ['missense_variant'] | MODERATE | IGF1R | ENST00000650285.1:c.1094G>A | ENSP00000497069.1:p.Arg365Gln | | 33 |
| 15 | 98896899 | ['C', 'T'] | 1 | 1.00 | ['missense_variant'] | MODERATE | IGF1R | ENST00000650285.1:c.1096C>T | ENSP00000497069.1:p.Arg366Trp | 1.19E-05 | 24 |

Supplementary Table 5 Genetic variants found in five height genes

| chr | position | alleles | AC | nt_qc.call_ | vep.consequence | vep.impact | vep.symbol | vep.hgvsc | vep.hgvsp | vep.gnom | vep.cadd_phred |
|---|---|---|---|---|---|---|---|---|---|---|---|
| 15 | 98899488 | ['T', 'A'] | 1 | 1.00 | ['missense_variant'] | MODERATE | IGF1R | ENST00000650285.1:c.1114T>A | ENSP00000497069.1:p.Ser372Thr | | 22 |
| 15 | 98899518 | ['G', 'A'] | 1 | 1.00 | ['missense_variant'] | MODERATE | IGF1R | ENST00000650285.1:c.1144G>A | ENSP00000497069.1:p.Glu382Lys | | 28 |
| 15 | 98899536 | ['G', 'A'] | 4 | 1.00 | ['missense_variant'] | MODERATE | IGF1R | ENST00000650285.1:c.1162G>A | ENSP00000497069.1:p.Val388Met | 0.000577 | 26 |
| 15 | 98899542 | ['A', 'G'] | 1 | 1.00 | ['missense_variant'] | MODERATE | IGF1R | ENST00000650285.1:c.1168A>G | ENSP00000497069.1:p.Ile390Val | | 23 |
| 15 | 98899568 | ['C', 'CT'] | 1 | 1.00 | ['frameshift_variant'] | HIGH | IGF1R | ENST00000650285.1:c.1196dup | ENSP00000497069.1:p.Leu399PhefsTer71 | 3.98E-06 | |
| 15 | 98899591 | ['G', 'A'] | 7 | 1.00 | ['missense_variant'] | MODERATE | IGF1R | ENST00000650285.1:c.1217G>A | ENSP00000497069.1:p.Arg406His | 3.18E-05 | 24 |
| 15 | 98908740 | ['G', 'A'] | 1 | 1.00 | ['missense_variant'] | MODERATE | IGF1R | ENST00000650285.1:c.1303G>A | ENSP00000497069.1:p.Asp435Asn | 7.96E-06 | 22 |
| 15 | 98908800 | ['T', 'C'] | 1 | 1.00 | ['missense_variant'] | MODERATE | IGF1R | ENST00000650285.1:c.1363T>C | ENSP00000497069.1:p.Cys455Arg | | 28 |
| 15 | 98908818 | ['C', 'T'] | 2 | 1.00 | ['missense_variant'] | MODERATE | IGF1R | ENST00000650285.1:c.1381C>T | ENSP00000497069.1:p.Arg461Cys | | 28 |
| 15 | 98908819 | ['G', 'A'] | 4 | 1.00 | ['missense_variant'] | MODERATE | IGF1R | ENST00000650285.1:c.1382G>A | ENSP00000497069.1:p.Arg461His | 1.99E-05 | 23 |
| 15 | 98908834 | ['C', 'T'] | 2 | 1.00 | ['missense_variant'] | MODERATE | IGF1R | ENST00000650285.1:c.1397C>T | ENSP00000497069.1:p.Thr466Met | 3.98E-06 | 27 |
| 15 | 98908848 | ['C', 'T'] | 3 | 1.00 | ['missense_variant'] | MODERATE | IGF1R | ENST00000650285.1:c.1411C>T | ENSP00000497069.1:p.Arg471Cys | 1.20E-05 | 27 |
| 15 | 98908849 | ['G', 'A'] | 4 | 1.00 | ['missense_variant'] | MODERATE | IGF1R | ENST00000650285.1:c.1412G>A | ENSP00000497069.1:p.Arg471His | 4.78E-05 | 25 |
| 15 | 98911326 | ['G', 'A'] | 1 | 1.00 | ['missense_variant'] | MODERATE | IGF1R | ENST00000650285.1:c.1474G>A | ENSP00000497069.1:p.Val492Ile | 0.000569 | 17 |
| 15 | 98911333 | ['A', 'T'] | 2 | 1.00 | ['missense_variant'] | MODERATE | IGF1R | ENST00000650285.1:c.1481A>T | ENSP00000497069.1:p.His494Leu | 3.98E-06 | 24 |
| 15 | 98911354 | ['C', 'T'] | 1 | 1.00 | ['missense_variant'] | MODERATE | IGF1R | ENST00000650285.1:c.1502C>T | ENSP00000497069.1:p.Ser501Leu | 0.000358 | 15 |
| 15 | 98911357 | ['A', 'G'] | 1 | 1.00 | ['missense_variant'] | MODERATE | IGF1R | ENST00000650285.1:c.1505A>G | ENSP00000497069.1:p.Lys502Arg | | 23 |
| 15 | 98911362 | ['C', 'T'] | 4 | 1.00 | ['missense_variant'] | MODERATE | IGF1R | ENST00000650285.1:c.1510C>T | ENSP00000497069.1:p.Arg504Cys | 1.19E-05 | 28 |
| 15 | 98911383 | ['C', 'T'] | 2 | 1.00 | ['missense_variant'] | MODERATE | IGF1R | ENST00000650285.1:c.1531C>T | ENSP00000497069.1:p.Arg511Trp | 2.39E-05 | 25 |
| 15 | 98911390 | ['G', 'A'] | 1 | 1.00 | ['missense_variant'] | MODERATE | IGF1R | ENST00000650285.1:c.1538G>A | ENSP00000497069.1:p.Arg513Gln | 7.95E-06 | 23 |
| 15 | 98911414 | ['T', 'C'] | 1 | 1.00 | ['missense_variant'] | MODERATE | IGF1R | ENST00000650285.1:c.1562T>C | ENSP00000497069.1:p.Ile521Thr | | 25 |
| 15 | 98911417 | ['G', 'C'] | 1 | 1.00 | ['missense_variant'] | MODERATE | IGF1R | ENST00000650285.1:c.1565G>C | ENSP00000497069.1:p.Ser522Thr | 3.98E-06 | 23 |
| 15 | 98911425 | ['G', 'A'] | 2 | 1.00 | ['missense_variant'] | MODERATE | IGF1R | ENST00000650285.1:c.1573G>A | ENSP00000497069.1:p.Val525Ile | 3.98E-05 | 25 |
| 15 | 98911441 | ['C', 'T'] | 5 | 1.00 | _variant', 'splice_regi | MODERATE | IGF1R | ENST00000650285.1:c.1589C>T | ENSP00000497069.1:p.Ala530Val | 1.19E-05 | 34 |
| 15 | 98913124 | ['C', 'G'] | 1 | 1.00 | ['missense_variant'] | MODERATE | IGF1R | ENST00000650285.1:c.1670C>G | ENSP00000497069.1:p.Pro557Arg | | 19 |
| 15 | 98913124 | ['C', 'T'] | 1 | 1.00 | ['missense_variant'] | MODERATE | IGF1R | ENST00000650285.1:c.1670C>T | ENSP00000497069.1:p.Pro557Leu | 7.95E-06 | 24 |
| 15 | 98913138 | ['G', 'A'] | 1 | 1.00 | ['missense_variant'] | MODERATE | IGF1R | ENST00000650285.1:c.1684G>A | ENSP00000497069.1:p.Val562Met | 1.59E-05 | 9 |
| 15 | 98913138 | ['G', 'T'] | 1 | 1.00 | ['missense_variant'] | MODERATE | IGF1R | ENST00000650285.1:c.1684G>T | ENSP00000497069.1:p.Val562Leu | 3.98E-06 | 7 |
| 15 | 98913147 | ['G', 'A'] | 2 | 1.00 | ['missense_variant'] | MODERATE | IGF1R | ENST00000650285.1:c.1693G>A | ENSP00000497069.1:p.Gly565Ser | 2.78E-05 | 22 |
| 15 | 98913160 | ['A', 'G'] | 1 | 1.00 | ['missense_variant'] | MODERATE | IGF1R | ENST00000650285.1:c.1706A>G | ENSP00000497069.1:p.His569Arg | 1.19E-05 | 19 |
| 15 | 98913189 | ['G', 'A'] | 1 | 1.00 | ['missense_variant'] | MODERATE | IGF1R | ENST00000650285.1:c.1735G>A | ENSP00000497069.1:p.Val579Ile | 6.76E-05 | 16 |
| 15 | 98913204 | ['G', 'A'] | 4 | 1.00 | ['missense_variant'] | MODERATE | IGF1R | ENST00000650285.1:c.1750G>A | ENSP00000497069.1:p.Val584Met | 1.99E-05 | 24 |
| 15 | 98913216 | ['A', 'G'] | 6 | 1.00 | ['missense_variant'] | MODERATE | IGF1R | ENST00000650285.1:c.1762A>G | ENSP00000497069.1:p.Met588Val | | 21 |
| 15 | 98913228 | ['G', 'A'] | 1 | 1.00 | ['missense_variant'] | MODERATE | IGF1R | ENST00000650285.1:c.1774G>A | ENSP00000497069.1:p.Asp592Asn | 2.39E-05 | 22 |
| 15 | 98913237 | ['C', 'G'] | 1 | 1.00 | ['missense_variant'] | MODERATE | IGF1R | ENST00000650285.1:c.1783C>G | ENSP00000497069.1:p.Arg595Gly | | 23 |
| 15 | 98913264 | ['A', 'T'] | 1 | 1.00 | ['missense_variant'] | MODERATE | IGF1R | ENST00000650285.1:c.1810A>T | ENSP00000497069.1:p.Ile604Phe | | 22 |
| 15 | 98913277 | ['C', 'CT'] | 1 | 1.00 | ['frameshift_variant'] | HIGH | IGF1R | ENST00000650285.1:c.1825dup | ENSP00000497069.1:p.Ser609PhefsTer73 | | |
| 15 | 98915984 | ['G', 'A'] | 1 | 1.00 | ['missense_variant'] | MODERATE | IGF1R | ENST00000650285.1:c.1849G>A | ENSP00000497069.1:p.Val617Ile | 2.78E-05 | 17 |
| 15 | 98916017 | ['A', 'C'] | 1 | 1.00 | ['missense_variant'] | MODERATE | IGF1R | ENST00000650285.1:c.1882A>C | ENSP00000497069.1:p.Ile628Leu | | 19 |
| 15 | 98916048 | ['A', 'G'] | 1 | 1.00 | ['missense_variant'] | MODERATE | IGF1R | ENST00000650285.1:c.1913A>G | ENSP00000497069.1:p.Asn638Ser | 1.19E-05 | 26 |
| 15 | 98916090 | ['C', 'G'] | 1 | 1.00 | ['missense_variant'] | MODERATE | IGF1R | ENST00000650285.1:c.1955C>G | ENSP00000497069.1:p.Pro652Arg | | 22 |
| 15 | 98916098 | ['G', 'A'] | 1 | 1.00 | ['missense_variant'] | MODERATE | IGF1R | ENST00000650285.1:c.1963G>A | ENSP00000497069.1:p.Gly655Ser | 8.35E-05 | 15 |
| 15 | 98916111 | ['G', 'A'] | 3 | 1.00 | ['missense_variant'] | MODERATE | IGF1R | ENST00000650285.1:c.1976G>A | ENSP00000497069.1:p.Arg659Gln | 0.00066 | 19 |
| 15 | 98916126 | ['C', 'T'] | 5 | 1.00 | ['missense_variant'] | MODERATE | IGF1R | ENST00000650285.1:c.1991C>T | ENSP00000497069.1:p.Ser664Phe | 1.19E-05 | 23 |
| 15 | 98916701 | ['G', 'A'] | 2 | 1.00 | ['missense_variant'] | MODERATE | IGF1R | ENST00000650285.1:c.2026G>A | ENSP00000497069.1:p.Gly676Ser | 1.59E-05 | 22 |
| 15 | 98916708 | ['T', 'C'] | 1 | 1.00 | ['missense_variant'] | MODERATE | IGF1R | ENST00000650285.1:c.2033T>C | ENSP00000497069.1:p.Ile678Thr | | 19 |
| 15 | 98916755 | ['G', 'T'] | 1 | 1.00 | ['missense_variant'] | MODERATE | IGF1R | ENST00000650285.1:c.2080G>T | ENSP00000497069.1:p.Gly694Trp | 3.98E-06 | 24 |
| 15 | 98916797 | ['G', 'A'] | 1 | 1.00 | ['missense_variant'] | MODERATE | IGF1R | ENST00000650285.1:c.2122G>A | ENSP00000497069.1:p.Glu708Lys | 3.98E-06 | 23 |
| 15 | 98916797 | ['G', 'C'] | 1 | 1.00 | ['missense_variant'] | MODERATE | IGF1R | ENST00000650285.1:c.2122G>C | ENSP00000497069.1:p.Glu708Gln | 3.98E-06 | 22 |
| 15 | 98916799 | ['G', 'C'] | 1 | 1.00 | ['missense_variant'] | MODERATE | IGF1R | ENST00000650285.1:c.2124G>C | ENSP00000497069.1:p.Glu708Asp | 3.98E-06 | 21 |
| 15 | 98916805 | ['G', 'C'] | 1 | 1.00 | ['missense_variant'] | MODERATE | IGF1R | ENST00000650285.1:c.2130G>C | ENSP00000497069.1:p.Gln710His | 3.98E-06 | 21 |
| 15 | 98916845 | ['A', 'G'] | 1 | 1.00 | ['missense_variant'] | MODERATE | IGF1R | ENST00000650285.1:c.2170A>G | ENSP00000497069.1:p.Asn724Asp | | 24 |
| 15 | 98916856 | ['C', 'G'] | 1 | 1.00 | ['missense_variant'] | MODERATE | IGF1R | ENST00000650285.1:c.2181C>G | ENSP00000497069.1:p.His727Gln | | 23 |
| 15 | 98916869 | ['G', 'A'] | 2 | 1.00 | ['missense_variant'] | MODERATE | IGF1R | ENST00000650285.1:c.2194G>A | ENSP00000497069.1:p.Val732Met | 7.97E-06 | 24 |
| 15 | 98922156 | ['G', 'T'] | 1 | 1.00 | ['missense_variant'] | MODERATE | IGF1R | ENST00000650285.1:c.2210G>T | ENSP00000497069.1:p.Arg737Met | | 29 |
| 15 | 98922204 | ['G', 'A'] | 1 | 1.00 | ['missense_variant'] | MODERATE | IGF1R | ENST00000650285.1:c.2258G>A | ENSP00000497069.1:p.Arg753Gln | 3.98E-06 | 23 |
| 15 | 98922221 | ['G', 'A'] | 2 | 1.00 | ['missense_variant'] | MODERATE | IGF1R | ENST00000650285.1:c.2275G>A | ENSP00000497069.1:p.Ala759Thr | | 8 |
| 15 | 98922286 | ['C', 'G'] | 2 | 1.00 | ['missense_variant'] | MODERATE | IGF1R | ENST00000650285.1:c.2340C>G | ENSP00000497069.1:p.Ser780Arg | | 20 |
| 15 | 98922326 | ['C', 'T'] | 1 | 1.00 | ['missense_variant'] | MODERATE | IGF1R | ENST00000650285.1:c.2380C>T | ENSP00000497069.1:p.Arg794Trp | 3.58E-05 | 28 |
| 15 | 98922345 | ['G', 'A'] | 1 | 1.00 | ['missense_variant'] | MODERATE | IGF1R | ENST00000650285.1:c.2399G>A | ENSP00000497069.1:p.Arg800His | 3.98E-06 | 30 |
| 15 | 98922381 | ['A', 'G'] | 1 | 1.00 | ['missense_variant'] | MODERATE | IGF1R | ENST00000650285.1:c.2435A>G | ENSP00000497069.1:p.Lys812Arg | | 17 |
| 15 | 98922396 | ['C', 'T'] | 6 | 1.00 | ['missense_variant'] | MODERATE | IGF1R | ENST00000650285.1:c.2450C>T | ENSP00000497069.1:p.Ala817Val | | 20 |
| 15 | 98922406 | ['C', 'G'] | 2 | 1.00 | ['missense_variant'] | MODERATE | IGF1R | ENST00000650285.1:c.2460C>G | ENSP00000497069.1:p.Phe820Leu | 4.00E-06 | 11 |
| 15 | 98922407 | ['G', 'A'] | 5 | 1.00 | ['missense_variant'] | MODERATE | IGF1R | ENST00000650285.1:c.2461G>A | ENSP00000497069.1:p.Val821Ile | 8.00E-06 | 24 |
| 15 | 98923948 | ['C', 'T'] | 4 | 1.00 | ['missense_variant'] | MODERATE | IGF1R | ENST00000650285.1:c.2558C>T | ENSP00000497069.1:p.Pro853Leu | 0.000167 | 19 |
| 15 | 98923997 | ['C', 'G'] | 1 | 1.00 | ['stop_gained'] | HIGH | IGF1R | ENST00000650285.1:c.2607C>G | ENSP00000497069.1:p.Tyr869Ter | | 35 |
| 15 | 98924534 | ['G', 'A'] | 4 | 1.00 | ['missense_variant'] | MODERATE | IGF1R | ENST00000650285.1:c.2632G>A | ENSP00000497069.1:p.Glu878Lys | 1.59E-05 | 23 |
| 15 | 98924551 | ['G', 'T'] | 1 | 1.00 | ['missense_variant'] | MODERATE | IGF1R | ENST00000650285.1:c.2649G>T | ENSP00000497069.1:p.Gln883His | 3.98E-06 | 22 |
| 15 | 98924558 | ['A', 'G'] | 1 | 1.00 | ['missense_variant'] | MODERATE | IGF1R | ENST00000650285.1:c.2656A>G | ENSP00000497069.1:p.Arg886Gly | 3.98E-06 | 21 |

Supplementary Table 5 Genetic variants found in five height genes

| chr | position | alleles | AC | nt_qc.call_ | vep.consequence | vep.impact | vep.symbol | vep.hgvsc | vep.hgvsp | vep.gnom | vep.cadd_phred |
|---|---|---|---|---|---|---|---|---|---|---|---|
| 15 | 98924595 | ['C', 'A'] | 5 | 1.00 | ['missense_variant'] | MODERATE | IGF1R | ENST00000650285.1:c.2693C>A | ENSP00000497069.1:p.Pro898Gln | 1.19E-05 | 26 |
| 15 | 98924625 | ['C', 'T'] | 1 | 1.00 | ['missense_variant'] | MODERATE | IGF1R | ENST00000650285.1:c.2723C>T | ENSP00000497069.1:p.Thr908Ile | | 23 |
| 15 | 98929558 | ['C', 'G'] | 1 | 1.00 | _variant', 'splice_regi | MODERATE | IGF1R | ENST00000650285.1:c.2783C>G | ENSP00000497069.1:p.Thr928Arg | | 14 |
| 15 | 98929590 | ['G', 'A'] | 2 | 1.00 | ['missense_variant'] | MODERATE | IGF1R | ENST00000650285.1:c.2815G>A | ENSP00000497069.1:p.Ala939Thr | | 17 |
| 15 | 98929602 | ['G', 'A'] | 2 | 1.00 | ['missense_variant'] | MODERATE | IGF1R | ENST00000650285.1:c.2827G>A | ENSP00000497069.1:p.Ala943Thr | 1.59E-05 | 18 |
| 15 | 98929617 | ['G', 'A'] | 1 | 1.00 | ['missense_variant'] | MODERATE | IGF1R | ENST00000650285.1:c.2842G>A | ENSP00000497069.1:p.Val948Met | 1.19E-05 | 23 |
| 15 | 98929644 | ['G', 'A'] | 1 | 1.00 | ['missense_variant'] | MODERATE | IGF1R | ENST00000650285.1:c.2869G>A | ENSP00000497069.1:p.Val957Ile | | 19 |
| 15 | 98929648 | ['T', 'C'] | 1 | 1.00 | ['missense_variant'] | MODERATE | IGF1R | ENST00000650285.1:c.2873T>C | ENSP00000497069.1:p.Phe958Ser | | 24 |
| 15 | 98930252 | ['G', 'C'] | 1 | 1.00 | ['missense_variant'] | MODERATE | IGF1R | ENST00000650285.1:c.2903G>C | ENSP00000497069.1:p.Gly968Ala | | 24 |
| 15 | 98930266 | ['T', 'C'] | 1 | 1.00 | ['missense_variant'] | MODERATE | IGF1R | ENST00000650285.1:c.2917T>C | ENSP00000497069.1:p.Tyr973His | | 32 |
| 15 | 98930299 | ['G', 'A'] | 1 | 1.00 | ['missense_variant'] | MODERATE | IGF1R | ENST00000650285.1:c.2950G>A | ENSP00000497069.1:p.Ala984Thr | 4.03E-06 | 23 |
| 15 | 98934863 | ['T', 'C'] | 1 | 1.00 | ['missense_variant'] | MODERATE | IGF1R | ENST00000650285.1:c.2996T>C | ENSP00000497069.1:p.Ile999Thr | | 30 |
| 15 | 98934875 | ['G', 'A'] | 1 | 1.00 | ['missense_variant'] | MODERATE | IGF1R | ENST00000650285.1:c.3008G>A | ENSP00000497069.1:p.Arg1003Gln | 3.18E-05 | 31 |
| 15 | 98934880 | ['C', 'A'] | 1 | 1.00 | ['missense_variant'] | MODERATE | IGF1R | ENST00000650285.1:c.3013C>A | ENSP00000497069.1:p.Leu1005Ile | | 25 |
| 15 | 98934908 | ['A', 'G'] | 1 | 1.00 | ['missense_variant'] | MODERATE | IGF1R | ENST00000650285.1:c.3041A>G | ENSP00000497069.1:p.Tyr1014Cys | 1.19E-05 | 32 |
| 15 | 98934982 | ['G', 'T'] | 2 | 1.00 | ['missense_variant'] | MODERATE | IGF1R | ENST00000650285.1:c.3115G>T | ENSP00000497069.1:p.Ala1039Ser | 7.96E-06 | 29 |
| 15 | 98934983 | ['C', 'T'] | 2 | 1.00 | ['missense_variant'] | MODERATE | IGF1R | ENST00000650285.1:c.3116C>T | ENSP00000497069.1:p.Ala1039Val | | 27 |
| 15 | 98934988 | ['A', 'G'] | 1 | 1.00 | ['missense_variant'] | MODERATE | IGF1R | ENST00000650285.1:c.3121A>G | ENSP00000497069.1:p.Met1041Val | 3.98E-06 | 22 |
| 15 | 98935328 | ['G', 'T'] | 1 | 1.00 | ['missense_variant'] | MODERATE | IGF1R | ENST00000650285.1:c.3199G>T | ENSP00000497069.1:p.Gly1067Cys | | 34 |
| 15 | 98935380 | ['G', 'A'] | 1 | 1.00 | ['missense_variant'] | MODERATE | IGF1R | ENST00000650285.1:c.3251G>A | ENSP00000497069.1:p.Arg1084Gln | 6.39E-06 | 27 |
| 15 | 98935392 | ['A', 'T'] | 1 | 1.00 | ['missense_variant'] | MODERATE | IGF1R | ENST00000650285.1:c.3263A>T | ENSP00000497069.1:p.Lys1088Ile | | 33 |
| 15 | 98935403 | ['C', 'T'] | 1 | 1.00 | ['missense_variant'] | MODERATE | IGF1R | ENST00000650285.1:c.3274C>T | ENSP00000497069.1:p.Arg1092Trp | 6.39E-06 | 24 |
| 15 | 98935404 | ['G', 'A'] | 5 | 1.00 | ['missense_variant'] | MODERATE | IGF1R | ENST00000650285.1:c.3275G>A | ENSP00000497069.1:p.Arg1092Gln | | 34 |
| 15 | 98935422 | ['T', 'C'] | 1 | 1.00 | ['missense_variant'] | MODERATE | IGF1R | ENST00000650285.1:c.3293T>C | ENSP00000497069.1:p.Met1098Thr | | 21 |
| 15 | 98939223 | ['C', 'T'] | 1 | 1.00 | ['missense_variant'] | MODERATE | IGF1R | ENST00000650285.1:c.3320C>T | ENSP00000497069.1:p.Pro1107Leu | | 28 |
| 15 | 98939233 | ['C', 'G'] | 2 | 1.00 | ['missense_variant'] | MODERATE | IGF1R | ENST00000650285.1:c.3330C>G | ENSP00000497069.1:p.Ser1110Arg | 7.95E-06 | 22 |
| 15 | 98939238 | ['T', 'G'] | 2 | 1.00 | ['missense_variant'] | MODERATE | IGF1R | ENST00000650285.1:c.3335T>G | ENSP00000497069.1:p.Met1112Arg | | 31 |
| 15 | 98942935 | ['C', 'T'] | 1 | 1.00 | ['missense_variant'] | MODERATE | IGF1R | ENST00000650285.1:c.3470C>T | ENSP00000497069.1:p.Thr1157Met | 3.98E-06 | 27 |
| 15 | 98942947 | ['A', 'G'] | 1 | 1.00 | ['missense_variant'] | MODERATE | IGF1R | ENST00000650285.1:c.3482A>G | ENSP00000497069.1:p.Tyr1161Cys | 2.39E-05 | 33 |
| 15 | 98942959 | ['A', 'G'] | 1 | 1.00 | ['missense_variant'] | MODERATE | IGF1R | ENST00000650285.1:c.3494A>G | ENSP00000497069.1:p.Tyr1165Cys | | 33 |
| 15 | 98942964 | ['C', 'T'] | 1 | 1.00 | ['missense_variant'] | MODERATE | IGF1R | ENST00000650285.1:c.3499C>T | ENSP00000497069.1:p.Arg1167Trp | | 32 |
| 15 | 98942970 | ['G', 'A'] | 1 | 1.00 | ['missense_variant'] | MODERATE | IGF1R | ENST00000650285.1:c.3505G>A | ENSP00000497069.1:p.Gly1169Arg | | 34 |
| 15 | 98948627 | ['A', 'T'] | 1 | 1.00 | ['missense_variant'] | MODERATE | IGF1R | ENST00000650285.1:c.3641A>T | ENSP00000497069.1:p.Gln1214Leu | | 32 |
| 15 | 98948645 | ['A', 'G'] | 1 | 1.00 | ['missense_variant'] | MODERATE | IGF1R | ENST00000650285.1:c.3659A>G | ENSP00000497069.1:p.Gln1220Arg | | 30 |
| 15 | 98948653 | ['C', 'T'] | 1 | 1.00 | ['missense_variant'] | MODERATE | IGF1R | ENST00000650285.1:c.3667C>T | ENSP00000497069.1:p.Arg1223Cys | | 33 |
| 15 | 98948691 | ['C', 'A'] | 2 | 1.00 | ['missense_variant'] | MODERATE | IGF1R | ENST00000650285.1:c.3705C>A | ENSP00000497069.1:p.Asp1235Glu | | 19 |
| 15 | 98957063 | ['T', 'A'] | 1 | 1.00 | _variant', 'splice_regi | MODERATE | IGF1R | ENST00000650285.1:c.3725T>A | ENSP00000497069.1:p.Phe1242Tyr | | 20 |
| 15 | 98957093 | ['A', 'G'] | 1 | 1.00 | ['missense_variant'] | MODERATE | IGF1R | ENST00000650285.1:c.3755A>G | ENSP00000497069.1:p.Asn1252Ser | 3.98E-06 | 24 |
| 15 | 98957146 | ['A', 'G'] | 1 | 1.00 | ['missense_variant'] | MODERATE | IGF1R | ENST00000650285.1:c.3808A>G | ENSP00000497069.1:p.Met1270Val | | 19 |
| 15 | 98957183 | ['G', 'A'] | 1 | 1.00 | ['missense_variant'] | MODERATE | IGF1R | ENST00000650285.1:c.3845G>A | ENSP00000497069.1:p.Ser1282Asn | 7.95E-06 | 30 |
| 15 | 98957184 | ['CGAG', '( | 1 | 1.00 | ['inframe_deletion'] | MODERATE | IGF1R | ENST00000650285.1:c.3850_3852del | ENSP00000497069.1:p.Glu1284del | 1.99E-05 | |
| 15 | 98957185 | ['G', 'A'] | 2 | 1.00 | ['missense_variant'] | MODERATE | IGF1R | ENST00000650285.1:c.3847G>A | ENSP00000497069.1:p.Glu1283Lys | 0.000103 | 25 |
| 15 | 98957206 | ['C', 'T'] | 1 | 1.00 | ['missense_variant'] | MODERATE | IGF1R | ENST00000650285.1:c.3868C>T | ENSP00000497069.1:p.Pro1290Ser | | 9 |
| 15 | 98957207 | ['C', 'T'] | 2 | 1.00 | ['missense_variant'] | MODERATE | IGF1R | ENST00000650285.1:c.3869C>T | ENSP00000497069.1:p.Pro1290Leu | 3.58E-05 | 17 |
| 15 | 98957214 | ['G', 'GCTG | 1 | 1.00 | ['inframe_insertion'] | MODERATE | IGF1R | ENST00000650285.1:c.3882_3887dup | ENSP00000497069.1:p.Asp1294_Leu1295dup | 3.98E-06 | |
| 15 | 98957221 | ['C', 'G'] | 1 | 1.00 | ['missense_variant'] | MODERATE | IGF1R | ENST00000650285.1:c.3883C>G | ENSP00000497069.1:p.Leu1295Val | | 20 |
| 15 | 98957239 | ['G', 'A'] | 1 | 1.00 | ['missense_variant'] | MODERATE | IGF1R | ENST00000650285.1:c.3901G>A | ENSP00000497069.1:p.Glu1301Lys | | 32 |
| 15 | 98957270 | ['C', 'T'] | 5 | 1.00 | ['missense_variant'] | MODERATE | IGF1R | ENST00000650285.1:c.3932C>T | ENSP00000497069.1:p.Ser1311Leu | 8.00E-05 | 22 |
| 15 | 98957273 | ['C', 'T'] | 1 | 1.00 | ['missense_variant'] | MODERATE | IGF1R | ENST00000650285.1:c.3935C>T | ENSP00000497069.1:p.Ser1312Phe | | 23 |
| 15 | 98957290 | ['G', 'C'] | 1 | 1.00 | ['missense_variant'] | MODERATE | IGF1R | ENST00000650285.1:c.3952G>C | ENSP00000497069.1:p.Asp1318His | | 24 |
| 15 | 98957290 | ['G', 'T'] | 1 | 1.00 | ['missense_variant'] | MODERATE | IGF1R | ENST00000650285.1:c.3952G>T | ENSP00000497069.1:p.Asp1318Tyr | 1.20E-05 | 24 |
| 15 | 98957300 | ['C', 'A'] | 1 | 1.00 | ['stop_gained'] | HIGH | IGF1R | ENST00000650285.1:c.3962C>A | ENSP00000497069.1:p.Ser1321Ter | 3.99E-06 | 47 |
| 15 | 98957307 | ['C', 'G'] | 1 | 1.00 | ['missense_variant'] | MODERATE | IGF1R | ENST00000650285.1:c.3969C>G | ENSP00000497069.1:p.His1323Gln | | 16 |
| 15 | 98957317 | ['A', 'G'] | 1 | 1.00 | ['missense_variant'] | MODERATE | IGF1R | ENST00000650285.1:c.3979A>G | ENSP00000497069.1:p.Asn1327Asp | 3.60E-05 | 21 |
| 15 | 98957326 | ['G', 'A'] | 5 | 1.00 | ['missense_variant'] | MODERATE | IGF1R | ENST00000650285.1:c.3988G>A | ENSP00000497069.1:p.Gly1330Ser | 5.20E-05 | 14 |
| 15 | 98957341 | ['G', 'T'] | 1 | 1.00 | ['missense_variant'] | MODERATE | IGF1R | ENST00000650285.1:c.4003G>T | ENSP00000497069.1:p.Val1335Phe | | 23 |
| 15 | 98957348 | ['G', 'A'] | 1 | 1.00 | ['missense_variant'] | MODERATE | IGF1R | ENST00000650285.1:c.4010G>A | ENSP00000497069.1:p.Arg1337His | 2.00E-05 | 26 |
| 15 | 98957362 | ['G', 'A'] | 2 | 1.00 | ['missense_variant'] | MODERATE | IGF1R | ENST00000650285.1:c.4024G>A | ENSP00000497069.1:p.Glu1342Lys | 4.00E-06 | 25 |
| 15 | 98957366 | ['G', 'T'] | 5 | 1.00 | ['missense_variant'] | MODERATE | IGF1R | ENST00000650285.1:c.4028G>T | ENSP00000497069.1:p.Arg1343Ile | 2.00E-05 | 23 |
| 15 | 98957383 | ['A', 'G'] | 2 | 1.00 | ['missense_variant'] | MODERATE | IGF1R | ENST00000650285.1:c.4045A>G | ENSP00000497069.1:p.Met1349Val | 8.02E-06 | 26 |
| 15 | 98957393 | ['G', 'C'] | 2 | 1.00 | ['missense_variant'] | MODERATE | IGF1R | ENST00000650285.1:c.4055G>C | ENSP00000497069.1:p.Gly1352Ala | | 28 |
| 15 | 98957395 | ['C', 'T'] | 2 | 1.00 | ['missense_variant'] | MODERATE | IGF1R | ENST00000650285.1:c.4057C>T | ENSP00000497069.1:p.Arg1353Cys | 8.04E-06 | 28 |
| 15 | 98957404 | ['G', 'A'] | 3 | 1.00 | ['missense_variant'] | MODERATE | IGF1R | ENST00000650285.1:c.4066G>A | ENSP00000497069.1:p.Glu1356Lys | 4.83E-05 | 25 |
| 15 | 98957404 | ['G', 'C'] | 1 | 1.00 | ['missense_variant'] | MODERATE | IGF1R | ENST00000650285.1:c.4066G>C | ENSP00000497069.1:p.Glu1356Gln | | 25 |
| 15 | 98957407 | ['C', 'T'] | 2 | 1.00 | ['missense_variant'] | MODERATE | IGF1R | ENST00000650285.1:c.4069C>T | ENSP00000497069.1:p.Arg1357Trp | 0.00025 | 25 |
| 15 | 98957408 | ['G', 'A'] | 2 | 1.00 | ['missense_variant'] | MODERATE | IGF1R | ENST00000650285.1:c.4070G>A | ENSP00000497069.1:p.Arg1357Gln | 3.22E-05 | 32 |
| 15 | 98957425 | ['C', 'T'] | 1 | 1.00 | ['stop_gained'] | HIGH | IGF1R | ENST00000650285.1:c.4087C>T | ENSP00000497069.1:p.Gln1363Ter | 8.07E-06 | 52 |
| 15 | 98957432 | ['C', 'T'] | 1 | 1.00 | ['missense_variant'] | MODERATE | IGF1R | ENST00000650285.1:c.4094C>T | ENSP00000497069.1:p.Ser1365Leu | 8.08E-06 | 26 |
| X | 630910 | ['A', 'G'] | 1 | 1.00 | ['missense_variant'] | MODERATE | SHOX | ENST00000381578.6:c.13A>G | ENSP00000370990.1:p.Thr5Ala | 7.99E-06 | 25 |

Supplementary Table 5 Genetic variants found in five height genes

| chr | position | alleles | AC | nt_qc.call_ | vep.consequence | vep.impact | vep.symbol | vep.hgvsc | vep.hgvsp | vep.gnom | vep.cadd_phred |
|---|---|---|---|---|---|---|---|---|---|---|---|
| X | 630922 | ['T', 'C'] | 1 | 1.00 | ['missense_variant'] | MODERATE | SHOX | ENST00000381578.6:c.25T>C | ENSP00000370990.1:p.Ser9Pro | | 25 |
| X | 630950 | ['A', 'T'] | 1 | 1.00 | ['missense_variant'] | MODERATE | SHOX | ENST00000381578.6:c.53A>T | ENSP00000370990.1:p.Asp18Val | 3.99E-06 | 22 |
| X | 630981 | ['TAAG', 'T | 3 | 1.00 | ['inframe_deletion'] | MODERATE | SHOX | ENST00000381578.6:c.88_90del | ENSP00000370990.1:p.Lys30del | | |
| X | 631004 | ['G', 'A'] | 1 | 1.00 | ['missense_variant'] | MODERATE | SHOX | ENST00000381578.6:c.107G>A | ENSP00000370990.1:p.Arg36Gln | | 26 |
| X | 631028 | ['C', 'A'] | 5 | 1.00 | ['missense_variant'] | MODERATE | SHOX | ENST00000381578.6:c.131C>A | ENSP00000370990.1:p.Ala44Glu | | 23 |
| X | 631045 | ['G', 'A'] | 1 | 1.00 | ['missense_variant'] | MODERATE | SHOX | ENST00000381578.6:c.148G>A | ENSP00000370990.1:p.Gly50Arg | | 22 |
| X | 631074 | ['C', 'G'] | 1 | 1.00 | ['missense_variant'] | MODERATE | SHOX | ENST00000381578.6:c.177C>G | ENSP00000370990.1:p.Ile59Met | | 0 |
| X | 631087 | ['G', 'C'] | 5 | 1.00 | ['missense_variant'] | MODERATE | SHOX | ENST00000381578.6:c.190G>C | ENSP00000370990.1:p.Gly64Arg | | 20 |
| X | 631097 | ['C', 'T'] | 1 | 1.00 | ['missense_variant'] | MODERATE | SHOX | ENST00000381578.6:c.200C>T | ENSP00000370990.1:p.Pro67Leu | | 23 |
| X | 631108 | ['T', 'C'] | 4 | 1.00 | ['missense_variant'] | MODERATE | SHOX | ENST00000381578.6:c.211T>C | ENSP00000370990.1:p.Phe71Leu | | 21 |
| X | 631115 | ['A', 'G'] | 1 | 1.00 | ['missense_variant'] | MODERATE | SHOX | ENST00000381578.6:c.218A>G | ENSP00000370990.1:p.Asp73Gly | | 20 |
| X | 631163 | ['G', 'C'] | 1 | 1.00 | ['missense_variant'] | MODERATE | SHOX | ENST00000381578.6:c.266G>C | ENSP00000370990.1:p.Arg89Thr | | 22 |
| X | 631169 | ['C', 'T'] | 1 | 1.00 | ['missense_variant'] | MODERATE | SHOX | ENST00000381578.6:c.272C>T | ENSP00000370990.1:p.Ala91Val | | 10 |
| X | 634631 | ['C', 'G'] | 1 | 1.00 | ['missense_variant'] | MODERATE | SHOX | ENST00000381578.6:c.291C>G | ENSP00000370990.1:p.Cys97Trp | | 24 |
| X | 634635 | ['G', 'A'] | 1 | 1.00 | ['missense_variant'] | MODERATE | SHOX | ENST00000381578.6:c.295G>A | ENSP00000370990.1:p.Glu99Lys | | 23 |
| X | 634637 | ['G', 'C'] | 1 | 1.00 | ['missense_variant'] | MODERATE | SHOX | ENST00000381578.6:c.297G>C | ENSP00000370990.1:p.Glu99Asp | | 7 |
| X | 634668 | ['G', 'T'] | 1 | 1.00 | ['missense_variant'] | MODERATE | SHOX | ENST00000381578.6:c.328G>T | ENSP00000370990.1:p.Asp110Tyr | | 26 |
| X | 634671 | ['G', 'C'] | 5 | 1.00 | ['missense_variant'] | MODERATE | SHOX | ENST00000381578.6:c.331G>C | ENSP00000370990.1:p.Gly111Arg | | 22 |
| X | 634676 | ['G', 'C'] | 1 | 1.00 | ['missense_variant'] | MODERATE | SHOX | ENST00000381578.6:c.336G>C | ENSP00000370990.1:p.Gln112His | | 24 |
| X | 634689 | ['CAG', 'C'] | 1 | 1.00 | ['frameshift_variant'] | HIGH | SHOX | ENST00000381578.6:c.352_353del | ENSP00000370990.1:p.Arg118AlafsTer63 | | |
| X | 634696 | ['G', 'A'] | 1 | 1.00 | ['missense_variant'] | MODERATE | SHOX | ENST00000381578.6:c.356G>A | ENSP00000370990.1:p.Arg119His | | 24 |
| X | 634705 | ['C', 'G'] | 1 | 1.00 | ['missense_variant'] | MODERATE | SHOX | ENST00000381578.6:c.365C>G | ENSP00000370990.1:p.Thr122Ser | 4.01E-06 | 24 |
| X | 634719 | ['G', 'C'] | 1 | 1.00 | ['missense_variant'] | MODERATE | SHOX | ENST00000381578.6:c.379G>C | ENSP00000370990.1:p.Glu127Gln | | 25 |
| X | 634741 | ['G', 'A'] | 1 | 1.00 | ['missense_variant'] | MODERATE | SHOX | ENST00000381578.6:c.401G>A | ENSP00000370990.1:p.Arg134Gln | | 25 |
| X | 634743 | ['C', 'T'] | 1 | 1.00 | ['missense_variant'] | MODERATE | SHOX | ENST00000381578.6:c.403C>T | ENSP00000370990.1:p.Leu135Phe | | 23 |
| X | 634771 | ['C', 'G'] | 1 | 1.00 | ['missense_variant'] | MODERATE | SHOX | ENST00000381578.6:c.431C>G | ENSP00000370990.1:p.Ala144Gly | | 24 |
| X | 634776 | ['A', 'C'] | 1 | 1.00 | ['missense_variant'] | MODERATE | SHOX | ENST00000381578.6:c.436A>C | ENSP00000370990.1:p.Met146Leu | | 24 |
| X | 634782 | ['G', 'A'] | 1 | 1.00 | ['missense_variant'] | MODERATE | SHOX | ENST00000381578.6:c.442G>A | ENSP00000370990.1:p.Glu148Lys | | 25 |
| X | 640851 | ['C', 'A'] | 1 | 1.00 | ['missense_variant'] | MODERATE | SHOX | ENST00000381578.6:c.517C>A | ENSP00000370990.1:p.Arg173Ser | | 19 |
| X | 640877 | ['A', 'C'] | 4 | 1.00 | _variant', 'splice_regi | MODERATE | SHOX | ENST00000381578.6:c.543A>C | ENSP00000370990.1:p.Lys181Asn | 3.58E-05 | 24 |
| X | 641001 | ['G', 'A'] | 4 | 1.00 | _variant', 'splice_regi | MODERATE | SHOX | ENST00000381578.6:c.547G>A | ENSP00000370990.1:p.Val183Ile | | 17 |
| X | 641037 | ['C', 'T'] | 2 | 1.00 | ['stop_gained'] | HIGH | SHOX | ENST00000381578.6:c.583C>T | ENSP00000370990.1:p.Arg195Ter | | 35 |
| X | 641038 | ['G', 'A'] | 2 | 1.00 | ['missense_variant'] | MODERATE | SHOX | ENST00000381578.6:c.584G>A | ENSP00000370990.1:p.Arg195Gln | | 25 |
| X | 641052 | ['G', 'A'] | 3 | 1.00 | ['missense_variant'] | MODERATE | SHOX | ENST00000381578.6:c.598G>A | ENSP00000370990.1:p.Val200Ile | | 22 |
| X | 641065 | ['C', 'T'] | 4 | 1.00 | ['missense_variant'] | MODERATE | SHOX | ENST00000381578.6:c.611C>T | ENSP00000370990.1:p.Ala204Val | | 23 |
| X | 641065 | ['C', 'A'] | 1 | 1.00 | ['missense_variant'] | MODERATE | SHOX | ENST00000381578.6:c.611C>A | ENSP00000370990.1:p.Ala204Asp | | 23 |
| X | 641068 | ['T', 'C'] | 1 | 1.00 | ['missense_variant'] | MODERATE | SHOX | ENST00000381578.6:c.614T>C | ENSP00000370990.1:p.Leu205Ser | | 23 |
| X | 644397 | ['G', 'A'] | 1 | 1.00 | ['missense_variant'] | MODERATE | SHOX | ENST00000381578.6:c.640G>A | ENSP00000370990.1:p.Ala214Thr | | 25 |
| X | 644418 | ['G', 'A'] | 1 | 1.00 | ['missense_variant'] | MODERATE | SHOX | ENST00000381578.6:c.661G>A | ENSP00000370990.1:p.Val221Met | | 26 |
| X | 644422 | ['C', 'T'] | 1 | 1.00 | ['missense_variant'] | MODERATE | SHOX | ENST00000381578.6:c.665C>T | ENSP00000370990.1:p.Ala222Val | | 23 |
| X | 644423 | ['C', 'CCAC | 1 | 1.00 | ['inframe_insertion'] | MODERATE | SHOX | ENST00000381578.6:c.670_681dup | ENSP00000370990.1:p.Ala224_His227dup | 3.36E-05 | |
| X | 644427 | ['G', 'A'] | 1 | 1.00 | ['missense_variant'] | MODERATE | SHOX | ENST00000381578.6:c.670G>A | ENSP00000370990.1:p.Ala224Thr | | 22 |
| X | 644428 | ['CGCACCC | 3 | 1.00 | ['inframe_deletion'] | MODERATE | SHOX | ENST00000381578.6:c.685_696del | ENSP00000370990.1:p.His229_Leu232del | | |
| X | 644434 | ['C', 'CGCA | 1 | 1.00 | ['inframe_insertion'] | MODERATE | SHOX | ENST00000381578.6:c.683_688dup | ENSP00000370990.1:p.Leu228_His229dup | | |
| X | 644464 | ['C', 'T'] | 1 | 1.00 | ['missense_variant'] | MODERATE | SHOX | ENST00000381578.6:c.707C>T | ENSP00000370990.1:p.Ala236Val | | 25 |
| X | 644484 | ['C', 'G'] | 1 | 1.00 | ['missense_variant'] | MODERATE | SHOX | ENST00000381578.6:c.727C>G | ENSP00000370990.1:p.Pro243Ala | | 21 |
| X | 644524 | ['CCGCCTC | 1 | 1.00 | ['inframe_deletion'] | MODERATE | SHOX | ENST00000381578.6:c.772_792del | ENSP00000370990.1:p.Ser258_Ala264del | | |
| X | 644539 | ['C', 'G'] | 1 | 1.00 | ['missense_variant'] | MODERATE | SHOX | ENST00000381578.6:c.782C>G | ENSP00000370990.1:p.Ala261Gly | | 19 |
| X | 644563 | ['G', 'T'] | 1 | 1.00 | ['missense_variant'] | MODERATE | SHOX | ENST00000381578.6:c.806G>T | ENSP00000370990.1:p.Ser269Ile | | 24 |
| X | 644578 | ['C', 'T'] | 1 | 1.00 | ['missense_variant'] | MODERATE | SHOX | ENST00000381578.6:c.821C>T | ENSP00000370990.1:p.Ser274Phe | | 26 |
| X | 644586 | ['G', 'A'] | 5 | 1.00 | ['missense_variant'] | MODERATE | SHOX | ENST00000381578.6:c.829G>A | ENSP00000370990.1:p.Ala277Thr | | 27 |
| X | 644598 | ['C', 'T'] | 1 | 1.00 | ['missense_variant'] | MODERATE | SHOX | ENST00000381578.6:c.841C>T | ENSP00000370990.1:p.Leu281Phe | | 25 |
| X | 644604 | ['G', 'C'] | 1 | 1.00 | ['missense_variant'] | MODERATE | SHOX | ENST00000381578.6:c.847G>C | ENSP00000370990.1:p.Ala283Pro | | 26 |
| X | 644608 | ['G', 'A'] | 2 | 1.00 | ['missense_variant'] | MODERATE | SHOX | ENST00000381578.6:c.851G>A | ENSP00000370990.1:p.Arg284Gln | | 25 |
| X | 644622 | ['G', 'T'] | 1 | 1.00 | ['missense_variant'] | MODERATE | SHOX | ENST00000381578.6:c.865G>T | ENSP00000370990.1:p.Ala289Ser | | 24 |
| X | 644622 | ['G', 'T'] | 1 | 1.00 | ['missense_variant'] | MODERATE | SHOX | ENST00000381578.6:c.865G>T | ENSP00000370990.1:p.Ala289Ser | | 24 |

Supplementary Table 6 Number of mutations described in HGMD for each of the five height genes

| | Complex | Deletion | | | | Gross Deletion | Gross Insertion | Indel | | | Insertion | | | | Mutation | | | | | | Prom | Splice | | | Grand Total |
|---|---|---|---|---|---|---|---|---|---|---|---|---|---|---|---|---|---|---|---|---|---|---|---|---|---|
| Gene/Disease Category | gross | frameshift | inframe | noncoding | total | gross | gross | frameshift | inframe | total | frameshift | inframe | noncoding | total | initiation | missense | nonsense | nonstop | synonymous | total | regulatory | canonical-splice | splice | total | |
| ***FGFR3*** | | | | | | | | | 3 | 3 | | | | | | 52 | | 7 | | 59 | | | | | 62 |
| Acanthosis nigricans | | | | | | | | | | | | | | | | 1 | | | | 1 | | | | | 1 |
| Achondroplasia | | | | | | | | | 1 | 1 | | | | | | 10 | | | | 10 | | | | | 11 |
| Achondroplasia with developmental delay & acanthosis nigricans | | | | | | | | | 1 | 1 | | | | | | | | | | | | | | | 1 |
| Achondroplasia with severe Platyspondyly | | | | | | | | | | | | | | | | 1 | | | | 1 | | | | | 1 |
| Camptodactyly, tall stature and hearing loss syndrome | | | | | | | | | | | | | | | | 1 | | | | 1 | | | | | 1 |
| Hypochondroplasia | | | | | | | | | 1 | 1 | | | | | | 27 | | | | 27 | | | | | 28 |
| Short stature | | | | | | | | | | | | | | | | 1 | | | | 1 | | | | | 1 |
| Skeletal dysplasia | | | | | | | | | | | | | | | | 1 | | | | 1 | | | | | 1 |
| Tall stature, lateral tibial deviation, scoliosis, hearing impairment, camptodactyly and arachnodactyly | | | | | | | | | | | | | | | | 1 | | | | 1 | | | | | 1 |
| Thanatophoric dwarfism | | | | | | | | | | | | | | | | 2 | | | | 2 | | | | | 2 |
| Thanatophoric dysplasia | | | | | | | | | | | | | | | | 6 | | 7 | | 13 | | | | | 13 |
| Thanatophoric dysplasia, type 2 | | | | | | | | | | | | | | | | 1 | | | | 1 | | | | | 1 |
| ***IGF1R*** | 2 | 1 | 1 | | 2 | 11 | 3 | | | | 3 | | | 3 | | 22 | 5 | | | 27 | | 1 | | 1 | 49 |
| Growth impairment | | | | | | | | | | | | | | | | 4 | | | | 4 | | | | | 4 |
| Growth retardation | | 1 | | | 1 | 3 | | | | | | | | | | 4 | 3 | | | 7 | | | | | 11 |
| Growth retardation & microcephaly | | | | | | | | | | | | | | | | 3 | | | | 3 | | | | | 3 |
| Growth retardation, intrauterine & postnatal | | | | | | | | | | | | | | | | 3 | | | | 3 | | | | | 3 |
| Growth retardation, microcephaly & Silver-Russell syndrome features | | | | | | | | | | | 1 | | | 1 | | | | | | | | | | | 1 |
| Insulin-like growth factor deficiency | | | | | | | | | | | | | | | | 1 | | | | 1 | | | | | 1 |
| Overgrowth | 1 | | | | | | 1 | | | | | | | | | | | | | | | | | | 2 |
| Overgrowth & variable intellectual disability | 1 | | | | | | 1 | | | | | | | | | | | | | | | | | | 2 |
| Short stature | | | | | | 8 | | | | | 2 | | | 2 | | 6 | 2 | | | 8 | | | | | 18 |
| Short stature, intellectual disability, Chiari malformation, microcephaly, hypotonia & lack of speech | | | | | | | 1 | | | | | | | | | | | | | | | | | | 1 |
| Short staure & intrauterine growth retardation | | | | | | | | | | | | | | | | 1 | | | | 1 | | | | | 1 |
| SHORT syndrome | | | | | | | | | | | | | | | | | | | | | | 1 | | 1 | 1 |
| Small for gestational age | | | 1 | | 1 | | | | | | | | | | | | | | | | | | | | 1 |
| ***NPPC*** | 4 | | | | | 1 | | | | | | | | | | 2 | | | | 2 | | | | | 7 |
| Overgrowth and bone anomalies | 3 | | | | | | | | | | | | | | | | | | | | | | | | 3 |
| Short stature, autosomal dominant | | | | | | | | | | | | | | | | 2 | | | | 2 | | | | | 2 |
| Skeletal overgrowth | 1 | | | | | 1 | | | | | | | | | | | | | | | | | | | 2 |
| ***NPR2*** | | 7 | 1 | | 8 | | | 1 | | 1 | | | | | | 49 | 9 | | | 58 | | 5 | | 5 | 72 |
| Abnormality of the skeletal system | | | | | | | | | | | | | | | | 2 | | | | 2 | | | | | 2 |
| Acromesomelic dysplasia, Maroteaux type | | 6 | | | 6 | | | 1 | | 1 | | | | | | 24 | 8 | | | 32 | | 4 | | 4 | 43 |
| Short stature | | 1 | 1 | | 2 | | | | | | | | | | | 19 | 1 | | | 20 | | 1 | | 1 | 23 |
| Tall stature | | | | | | | | | | | | | | | | 2 | | | | 2 | | | | | 2 |
| Tall stature, scoliosis & macrodactyly of great toes | | | | | | | | | | | | | | | | 2 | | | | 2 | | | | | 2 |
| ***SHOX*** | 8 | 15 | 2 | 1 | 18 | 157 | 32 | 1 | 1 | 2 | 6 | 2 | | 8 | 1 | 71 | 20 | 1 | | 93 | 1 | 5 | | 5 | 324 |
| Dyschondrosteosis | 1 | | | | | | | | | | 1 | | | 1 | | 2 | | | | 2 | | | | | 4 |
| Langer mesomelic dysplasia | | | | | | 10 | | | | | 2 | | | 2 | | 2 | | | | 2 | | | | | 14 |
| Langer mesomelic dysplasia, milder phenotype | | | | | | 1 | | | | | | | | | | | | | | | | | | | 1 |
| Leri-Weill dyschondrosteosis | 3 | 9 | | | 9 | 78 | 9 | 1 | 1 | 2 | 2 | 1 | | 3 | 1 | 33 | 12 | 1 | | 47 | 1 | 1 | | 1 | 153 |
| Leri-Weill dyschondrosteosis with normal stature | | | | | | 1 | | | | | | | | | | | | | | | | | | | 1 |
| Leri-Weill dyschondrosteosis with short stature | | | | | | 2 | | | | | | | | | | | | | | | | | | | 2 |
| Madelung deformity | 1 | 1 | | | 1 | 1 | | | | | | | | | | | 1 | | | 1 | | 1 | | 1 | 5 |
| Short stature | 2 | 5 | 2 | 1 | 8 | 60 | 23 | | | | | 1 | | 1 | | 34 | 6 | | | 40 | | 3 | | 3 | 137 |
| Short stature with Cornelia de Lange-like syndrome | | | | | | 1 | | | | | | | | | | | | | | | | | | | 1 |
| Short stature, mental retardation & facial dysmorphisms | 1 | | | | | | | | | | | | | | | | | | | | | | | | 1 |
| SHOX-related disease | | | | | | 3 | | | | | 1 | | | 1 | | | 1 | | | 1 | | | | | 5 |
| Grand Total | 27 | 372 | 38 | 21 | 431 | 269 | 44 | 18 | 13 | 31 | 141 | 10 | 8 | 159 | 4 | 1324 | 316 | 8 | 2 | 1654 | 1 | 253 | 23 | 276 | 2892 |
| Grand Total | 27 | 372 | 38 | 21 | 431 | 269 | 44 | 18 | 13 | 31 | 141 | 10 | 8 | 159 | 4 | 1324 | 316 | 8 | 2 | 1654 | 1 | 253 | 23 | 276 | 2892 |

**Supplementary Table 7.** Proposed mechanisms of identified genes for growth regulation. Lines of evidence from bi-directional genetic effects (gain of function, loss of function) and the associated impact on growth in human, mouse and cell line models.

| | | | Human | | Mouse | | Cell Line | |
|---|---|---|---|---|---|---|---|---|
| **Gene** | **Protein** | **Mechanism** | **GOF** | **LOF** | **GOF** | **LOF** | **GOF** | **LOF** |
| *IGF1R* | Insulin-like growth factor 1 receptor | Regulates cell growth and proliferation after activation by IGF1, IGF2, and insulin | ↑ [1] | ↓ [2] | | ↓ [3] | ↑ [4] | ↓ [4] |
| *FGFR3* | *Fibroblast growth factor receptor 3* | *Activates kinase (ERK)/mitogen-activated protein kinase (MAPK) pathway. Regulation of chondrocyte proliferation.* | ↓ [5] | ↑ [6] | ↓ [7] | ↑ [8] | ↓ [9] | |
| *NPPC* | C-type natriuretic peptide (CNP) | Promotes endochondral ossification in chondrocytes after binding NPR2. | ↑ [10] | ↓ [11] | ↑ [10] | ↓ [12] | ↑ [13] | ↓ [13] |
| *NPR2* | Natriuretic peptide receptor B. | Primary receptor for CNP. Increases guanylyl cyclase activity (cGMP) and counteracts FGFR3 signaling. | ↑ [14,15] | ↓ [16,17] | | ↓ [18] | ↑ [15] | ↓ [17] |
| *SHOX* | Short stature homeobox | Multiple. Represses FGFR3 transcription. Up-regulates NPPB. Regulates cell-death. | ↑ [19] | ↓ [20] | | ↓ [21] | | ↓ [22] |

Supplementary Table 8 NPR2 mutations tested in functional experiments

| | Replicates | | | | | | |
|---|---|---|---|---|---|---|---|
| **Variant** | **1** | **2** | **3** | **Mean 95%CI** | **Source** | **b** | **ac** |
| G21R | 2.61 | 2.60 | 3.13 | 2.78 [2.19 , 3.37 ] | This_GOF | 1.99 | 1 |
| A48S | 0.07 | 0.07 | 0.06 | 0.07 [0.05 , 0.08 ] | HGMD_ISS | NA | NA |
| A59T | 4.24 | 3.18 | 3.79 | 3.74 [2.69 , 4.78 ] | This_GOF | 1.76 | 2 |
| V102A | 0.13 | 0.09 | 0.12 | 0.11 [0.07 , 0.15 ] | This_LoF | -2.20 | 1 |
| R110G | 0.68 | 0.80 | 0.62 | 0.7 [0.52 , 0.88 ] | This_GOF | 3.24 | 1 |
| A164G | 0.35 | 0.26 | 0.28 | 0.3 [0.2 , 0.39 ] | This_Low | -0.21 | 13 |
| I226V | 0.43 | 0.46 | 0.48 | 0.46 [0.41 , 0.5 ] | This_LoF | -2.02 | 1 |
| Y250X | 0.02 | 0.02 | 0.02 | 0.02 [0.01 , 0.02 ] | This_LoF | -2.19 | 1 |
| R263H | 1.73 | 1.82 | 2.04 | 1.86 [1.55 , 2.18 ] | This_GOF | 1.25 | 2 |
| T297M | 0.03 | 0.02 | 0.04 | 0.03 [0.01 , 0.05 ] | HGMD_ADMD | -2.16 | 2 |
| P301S | 0.05 | 0.08 | 0.13 | 0.08 [0 , 0.17 ] | This_LoF | -2.67 | 1 |
| R318Q | 1.00 | 0.92 | 0.87 | 0.93 [0.79 , 1.06 ] | This_Neutral | -0.03 | 7 |
| R358W | 1.02 | 1.01 | 1.03 | 1.02 [1 , 1.04 ] | This_High | 0.16 | 44 |
| E359A | 3.20 | 2.88 | 2.91 | 3 [2.65 , 3.34 ] | This_GOF | 1.33 | 2 |
| I364fs | 0.01 | 0.01 | 0.02 | 0.01 [0.01 , 0.02 ] | HGMD_ADMD | NA | NA |
| E389D | 0.06 | 0.08 | 0.07 | 0.07 [0.05 , 0.09 ] | HGMD_ISS | NA | NA |
| E415K | 0.49 | 0.53 | 0.40 | 0.47 [0.35 , 0.6 ] | This_Neutral | -0.08 | 4 |
| K480N | 1.77 | 1.62 | 1.68 | 1.69 [1.54 , 1.84 ] | This_GOF | 1.57 | 3 |
| A488P | 1.27 | 1.60 | 0.95 | 1.27 [0.63 , 1.91 ] | HGMD_GOF | NA | NA |
| I494S | 0.06 | 0.02 | 0.03 | 0.04 [0 , 0.08 ] | HGMD_ISS | NA | NA |
| Q500X | 0.02 | 0.01 | 0.02 | 0.02 [0.01 , 0.03 ] | HGMD_ADMD | NA | NA |
| H508D | 1.80 | 1.70 | 1.72 | 1.74 [1.64 , 1.84 ] | This_LoF | -2.28 | 2 |
| G541S | 0.73 | 0.34 | 0.53 | 0.53 [0.16 , 0.91 ] | This_Neutral | 0.67 | 2 |
| A549T | 0.71 | 0.71 | 0.70 | 0.71 [0.69 , 0.72 ] | HGMD_ISS | -0.21 | 2 |
| R562Q | 1.89 | 1.16 | 1.61 | 1.55 [0.83 , 2.28 ] | HGMD_GOF | NA | NA |
| R601H | 0.50 | 0.50 | 0.43 | 0.48 [0.4 , 0.55 ] | This_Low | -0.50 | 14 |
| R601S | 0.04 | 0.05 | 0.09 | 0.06 [0.01 , 0.11 ] | HGMD_ADMD | NA | NA |
| E609K | 0.97 | 1.07 | 0.98 | 1.01 [0.9 , 1.12 ] | This_Low | -0.47 | 13 |
| S721R | 0.97 | 1.05 | 0.85 | 0.96 [0.76 , 1.16 ] | This_Neutral | -0.04 | 3 |

#### Supplementary Table 8 NPR2 mutations tested in functional experiments

| | Replicates | | | | | | |
|---|---|---|---|---|---|---|---|
| **Variant** | **1** | **2** | **3** | **Mean 95%CI** | **Source** | **b** | **ac** |
| E727Q | 1.80 | 1.65 | 1.66 | 1.7 [1.53 , 1.87 ] | This_GOF | 1.35 | 1 |
| R745Q | 0.71 | 0.53 | 0.66 | 0.63 [0.44 , 0.82 ] | This_Neutral | 0.02 | 3 |
| R819C | 0.14 | 0.15 | 0.13 | 0.14 [0.11 , 0.17 ] | HGMD_ISS | -0.58 | 1 |
| T861I | 0.09 | 0.08 | 0.07 | 0.08 [0.06 , 0.1 ] | This_LoF | -2.12 | 1 |
| V883M | 0.32 | 0.38 | 0.31 | 0.33 [0.26 , 0.41 ] | HGMD_GOF | NA | NA |
| T907M | 0.01 | 0.01 | 0.02 | 0.01 [0.01 , 0.02 ] | HGMD_ADMD | NA | NA |
| G917R | 0.02 | 0.02 | 0.01 | 0.02 [0.01 , 0.02 ] | This_LoF | -2.74 | 1 |
| R921Q | 0.23 | 0.23 | 0.21 | 0.22 [0.2 , 0.25 ] | This_LoF | -1.91 | 1 |
| R957C | 0.03 | 0.03 | 0.03 | 0.03 [0.03 , 0.03 ] | HGMD_ADMD | -0.86 | 1 |
| L1009V | 0.09 | 0.08 | 0.07 | 0.08 [0.07 , 0.09 ] | This_LoF | -2.28 | 2 |
| Wild-Type | 0.95 | 1.05 | 1.00 | 1 [0.91 , 1.09 ] | | | |

Abreviations: source, Source for choosing this variant for the assay; b, beta estimate for height in the ukbiobank data; AC, allele count in the ukbiobank data

Supplementary Table 9 Median height of PRS x cGMP category in NPR2 carriers

| PRS group | cGMP group | n | ZPRS | cGMP | zheight |
|---|---|---|---|---|---|
| 1 | (0.0, 0.2] | 3 | -1.38 | 0.08 | -2.60 |
| 1 | (0.2, 0.5] | 8 | -1.48 | 0.47 | -1.17 |
| 1 | (0.5, 1.5] | 14 | -1.00 | 1.02 | -0.56 |
| 1 | (1.5, 4.0] | 0 | NA | NA | NA |
| 2 | (0.0, 0.2] | 1 | -0.63 | 0.03 | -2.10 |
| 2 | (0.2, 0.5] | 2 | -0.51 | 0.39 | -0.38 |
| 2 | (0.5, 1.5] | 20 | -0.47 | 1.01 | -0.06 |
| 2 | (1.5, 4.0] | 1 | -0.77 | 2.78 | 2.05 |
| 3 | (0.0, 0.2] | 2 | 0.07 | 0.05 | -2.09 |
| 3 | (0.2, 0.5] | 9 | -0.03 | 0.48 | -0.45 |
| 3 | (0.5, 1.5] | 13 | 0.12 | 1.01 | -0.05 |
| 3 | (1.5, 4.0] | 0 | NA | NA | NA |
| 4 | (0.0, 0.2] | 1 | 0.73 | 0.03 | -0.80 |
| 4 | (0.2, 0.5] | 8 | 0.53 | 0.30 | -0.21 |
| 4 | (0.5, 1.5] | 11 | 0.59 | 1.02 | 0.29 |
| 4 | (1.5, 4.0] | 4 | 0.73 | 2.43 | 1.75 |
| 5 | (0.0, 0.2] | 1 | 1.32 | 0.14 | -0.52 |
| 5 | (0.2, 0.5] | 6 | 1.43 | 0.38 | 0.31 |
| 5 | (0.5, 1.5] | 14 | 1.38 | 1.02 | 0.38 |
| 5 | (1.5, 4.0] | 4 | 1.91 | 1.72 | 1.12 |

Abreviations: n, number of individuals in group; ZPRS,Median of Polygenic risk score in group; cGMP,median of cyclic GMP in group; zheight, median of height in group

Supplementary Table 10 List of potential bidirectional effect selected tagets from HGMD

| Gene | Phenotype | Number of variants |
|---|---|---|
| ABCA1 | Altered HDL cholesterol levels | 2 |
| ABCA1 | HDL deficiency | 62 |
| ABCA1 | High HDL cholesterol | 1 |
| ABCA1 | Increased plasma HDL cholesterol | 7 |
| ABCA1 | Low HDL cholesterol | 5 |
| ABCA1 | Reduced plasma HDL cholesterol | 51 |
| ABCC8 | Hyperglycaemia | 1 |
| ABCC8 | Hyperinsulinaemic hypoglycaemia | 10 |
| ABCC8 | Hypoglycaemia, hyperinsulinaemic | 4 |
| ABCC8 | Hypoglycaemia, persistent hyperinsulinaemic | 50 |
| ABCG1 | High HDL cholesterol | 2 |
| ABCG1 | Low HDL cholesterol | 1 |
| ADORA3 | Elevated SERT activity | 1 |
| ADORA3 | Reduced SERT activity | 1 |
| AKT2 | Hypoglycaemia | 1 |
| AKT2 | Severe insulin resistance and diabetes mellitus | 1 |
| AKT3 | Macrocephaly & speech delay | 1 |
| AKT3 | Macrocephaly, megalencephaly & developmental delay | 1 |
| AKT3 | Megalencephaly-capillary malformation / megalencephaly-polymicrogyria-polydactyly-hydrocephalus syndrome | 1 |
| AKT3 | Megalencephaly-polymicrogyria-polydactyly-hydrocephalus syndrome | 1 |
| AKT3 | Megalencephaly, polymicrogyria, epilepsy and hypoglycaemia | 1 |
| AKT3 | Microcephaly | 6 |
| AKT3 | Microcephaly & abnormalities of the corpus callosum | 4 |
| AKT3 | Microcephaly, abnormalities of the corpus callosum & seizures | 2 |
| AKT3 | Microcephaly, hypotonia, feeding difficulties, developmental delay & dysmorphic features | 1 |
| ALPP | Recurrent spontaneous abortion, decreased risk | 1 |
| ALPP | Recurrent spontaneous abortion, increased risk | 1 |
| ANGPTL3 | Higher plasma triglyceride level | 2 |
| ANGPTL3 | Lower plasma triglyceride level | 12 |
| ANGPTL4 | Higher plasma triglyceride level | 1 |
| ANGPTL4 | Lower plasma triglyceride level | 21 |
| ANGPTL4 | Lower plasma triglyceride level, association with | 1 |
| ANGPTL8 | Increased HDL cholesterol levels | 1 |
| ANGPTL8 | Lower plasma LDL and HDL levels, association with | 1 |
| APOA1 | Elevated HDL-cholesterol, association with | 1 |
| APOA1 | HDL deficiency | 10 |
| APOA1 | HDL deficiency with periorbital xanthelasmas | 1 |
| APOA1 | Hyperlipidaemia | 1 |
| APOA1 | Low HDL cholesterol | 1 |
| APOA1 | Reduced plasma HDL cholesterol | 3 |
| APOB | HDL cholesterol, association with | 1 |
| APOB | High LDL cholesterol | 1 |
| APOB | Hypercholesterolaemia | 45 |
| APOB | Hypercholesterolaemia, association with | 2 |
| APOB | Hypertriglyceridaemia | 28 |
| APOB | Hypocholesterolaemia | 18 |
| APOB | Hypocholesterolaemia, association with | 1 |
| APOB | Hypocholesterolaemia, steatosis and liver cancer | 1 |
| APOB | Increased cholesterol levels | 2 |
| APOB | Low HDL cholesterol | 1 |
| APOB | Lower non-HDL cholesterol levels | 1 |
| APOB | Total cholesterol levels, association with | 1 |
| APOC3 | High triglyceride levels, association with | 1 |
| APOC3 | Hypertriglyceridaemia | 1 |
| APOC3 | Low triglyceride levels | 1 |
| BDKRB2 | Essential hypertension, increased risk, association with in Asians | 1 |
| BDKRB2 | Hypertension, association with | 1 |
| BDKRB2 | Reduced blood pressure, association with | 2 |
| CASR | Hypercalcaemia | 5 |
| CASR | Hypercalcaemia and hypercalciuria | 1 |
| CASR | Hypercalcaemia, association with | 1 |
| CASR | Hypercalcaemia, hypocalciuric | 245 |
| CASR | Hypercalcaemia, hypocalciuric & hyperparathyroidism | 1 |
| CASR | Hypercalcaemia, hypocalciuric & hypoparathyroidism | 3 |
| CASR | Hypercalciuria, hypocalcaemic | 17 |
| CASR | Hyperparathyroidism | 27 |
| CASR | Hyperparathyroidism, neonatal primary | 6 |
| CASR | Hypocalcaemia | 41 |
| CASR | Hypocalcaemia with hypercalciuria | 6 |
| CASR | Hypocalcaemia, autosomal dominant | 2 |
| CASR | Hypocalcaemia, with Bartter syndrome | 1 |
| CASR | Hypoparathyroidism | 19 |
| CASR | Hypoparathyroidism, autosomal dominant | 3 |
| CASR | Increased blood ionised calcium level | 1 |
| CCL5 | HIV 1, delayed disease progression, association | 1 |
| CCL5 | HIV 1, rapid disease progression, association | 1 |
| CCR5 | HIV 1, resistance to | 1 |
| CCR5 | HIV 1, resistance to, association with | 7 |
| CCR5 | Increased HIV1, perinatal transmission, association | 1 |
| CCR5 | Kaposi sarcoma | 1 |

Supplementary Table 10 List of potential bidirectional effect selected tagets from HGMD

| Gene | Phenotype | Number of variants |
|---|---|---|
| CD40LG | CD40L deficiency | 1 |
| CD40LG | Increased soluble CD40L levels | 1 |
| CETP | HDL-C levels, in African blacks, association with | 1 |
| CETP | Higher HDL cholesterol level | 12 |
| CETP | Higher HDL cholesterol level, association with | 3 |
| CETP | Increased HDL cholesterol levels | 1 |
| CETP | Increased HDL-C levels in males | 2 |
| CETP | Low HDL cholesterol | 1 |
| CETP | Lower HDL cholesterol level | 6 |
| CHGA | Hypertension, association with | 1 |
| CHGA | Reduced diastolic blood pressure, association with | 2 |
| CRP | Altered CRP levels | 5 |
| CRP | Increased CRP level | 2 |
| CRP | Reduced CRP levels | 1 |
| CYP11B1 | Aldosteronism | 1 |
| CYP11B1 | Hypoaldosteronism | 5 |
| CYP19A1 | Aromatase deficiency | 33 |
| CYP19A1 | Aromatase excess syndrome | 15 |
| CYP19A1 | Breast cancer, association with | 1 |
| CYP19A1 | Breast cancer, decreased risk, association with | 1 |
| CYP19A1 | Breast cancer, increased risk, association with | 2 |
| CYP2D6 | Intermediate metaboliser | 4 |
| CYP2D6 | Poor metaboliser | 52 |
| CYP2D6 | Ultrarapid metaboliser | 3 |
| EGLN1 | Erythrocytosis | 20 |
| EGLN1 | Erythrocytosis and paraganglioma | 1 |
| EGLN1 | Erythrocytosis, familial, 3 | 1 |
| EGLN1 | Pheochromocytoma/paraganglioma-polycythemia | 1 |
| EGLN1 | Polycythaemia | 4 |
| EGLN1 | Polycythaemia vera | 1 |
| EGLN1 | Reduced haemoglobin levels | 2 |
| F2 | Dysprothrombinaemia | 13 |
| F2 | Hypoprothrombinaemia | 5 |
| F2 | Increased plasma prothrombin levels | 1 |
| F2 | Prothrombin deficiency | 39 |
| F2 | Thrombosis | 2 |
| F2 | Thrombosis, venous | 4 |
| F2 | Venous thromboembolism | 1 |
| F2RL3 | Increased PAR4-induced platelet aggregation | 1 |
| F2RL3 | Reduced PAR4-induced platelet aggregation | 1 |
| F2RL3 | Reduced PAR4-induced platelet response | 1 |
| F5 | Bleeding disorder | 1 |
| F5 | Deep vein thrombosis | 5 |
| F5 | Factor V deficiency | 139 |
| F5 | Reduced factor V levels | 1 |
| F5 | Thrombosis | 4 |
| F5 | Thrombosis, increased risk | 5 |
| F5 | Thrombosis, increased risk, association with | 3 |
| F7 | Factor VII deficiency | 295 |
| F7 | Factor VII deficiency, association with | 2 |
| F7 | Increased plasma F7 levels, association with | 1 |
| F7 | Reduced plasma F7 levels, association with | 1 |
| F8 | Decreased factor VIII activity | 1 |
| F8 | Factor VIII deficiency | 3 |
| F8 | Haemophilia A | 3067 |
| F8 | Haemophilia A & moyamoya disease | 1 |
| F8 | Haemophilia A, modifier of | 1 |
| F8 | Venous thromboembolism, susceptibility to | 1 |
| F9 | Deep vein thrombosis, reduced risk | 1 |
| F9 | Haemophilia B | 1251 |
| F9 | Haemophilia B, severe | 4 |
| F9 | Reduced factor IX level in females | 1 |
| F9 | Thrombophilia | 1 |
| FGA | Afibrinogenaemia | 56 |
| FGA | Afibrinogenaemia / hypofibrinogenaemia | 1 |
| FGA | Afibrinogenaemia with recurrent venous thromboembolism | 1 |
| FGA | Decreased fibrinogen levels | 2 |
| FGA | Deep vein thrombosis | 1 |
| FGA | Dysfibrinogenaemia | 37 |
| FGA | Haemorrhages | 1 |
| FGA | Hypodysfibrinogenaemia | 2 |
| FGA | Hypofibrinogenaemia | 13 |
| FGA | Menorrhagia | 1 |
| FGA | Thrombosis | 1 |
| FGA | Venous thromboembolism | 1 |
| FGA | Venous thromboembolism, susceptibility, association with | 3 |
| FGB | Afibrinogenaemia | 26 |
| FGB | Afibrinogenaemia / hypofibrinogenaemia | 1 |
| FGB | Dysfibrinogenaemia | 16 |
| FGB | Haemorrhages | 3 |

Supplementary Table 10 List of potential bidirectional effect selected tagets from HGMD

| Gene | Phenotype | Number of variants |
|---|---|---|
| FGB | Hypodysfibrinogenaemia | 2 |
| FGB | Hypofibrinogenaemia | 31 |
| FGB | Increased clot stiffness, association with | 1 |
| FGB | Increased plasma fibrinogen levels | 1 |
| FGB | Thrombotic tendency | 1 |
| FGB | Venous thromboembolism, protection against, in Caucasians, association with | 1 |
| FGF23 | Calcium nephrolithiasis with renal phosphate leak, association with | 1 |
| FGF23 | Hyperostosis-hyperphosphatemia syndrome | 1 |
| FGF23 | Rickets, hypophosphataemic, autosomal dominant | 4 |
| FGF23 | Tumoural calcinosis with hyperphosphataemia | 9 |
| FGFR3 | Achondroplasia | 11 |
| FGFR3 | Achondroplasia with developmental delay & acanthosis nigricans | 1 |
| FGFR3 | Achondroplasia with severe Platyspondyly | 1 |
| FGFR3 | Hypochondroplasia | 25 |
| FGFR3 | Short stature | 1 |
| FGFR3 | Tall stature, lateral tibial deviation, scoliosis, hearing impairment, camptodactyly and arachnodactyly | 1 |
| FGFR3 | Thanatophoric dwarfism | 2 |
| FGFR3 | Thanatophoric dysplasia | 12 |
| FGFR3 | Thanatophoric dysplasia, type 2 | 1 |
| FGG | Afibrinogenaemia | 14 |
| FGG | Afibrinogenaemia / hypofibrinogenaemia | 1 |
| FGG | Decreased plasma fibrinogen levels | 1 |
| FGG | Deep venous thrombosis, increased risk, association with | 1 |
| FGG | Dysfibrinogenaemia | 50 |
| FGG | Hypodysfibrinogenaemia | 6 |
| FGG | Hypofibrinogenaemia | 42 |
| FGG | Hypofibrinogenaemia with hepatic storage | 3 |
| FGG | Menorrhagia | 1 |
| FSHR | FSHR activation | 2 |
| FSHR | FSHR inactivation | 1 |
| FSHR | Hypergonadotropic hypogonadism | 5 |
| FSHR | Hypergonadotropic ovarian failure | 1 |
| FSHR | Ovarian failure, primary | 1 |
| FSHR | Ovarian hyperstimulation syndrome | 8 |
| FSHR | Premature ovarian failure | 1 |
| FSHR | Primary amenorrhea | 6 |
| FSHR | Secondary amenorrhea | 2 |
| FSHR | Twinning, dizygotic | 1 |
| G6PC2 | Decreased fasting plasma glucose levels, association with | 1 |
| G6PC2 | Fasting plasma glucose level, association with | 5 |
| G6PC2 | Increased fasting plasma glucose levels, association with | 1 |
| GALNT2 | Higher plasma HDL cholesterol | 4 |
| GALNT2 | Higher plasma HDL cholesterol, modifier of | 1 |
| GALNT2 | Lower levels of phospholipid transfer protein and HDL cholesterol | 2 |
| GCK | Hyperglycaemia | 17 |
| GCK | Hyperglycaemia, exercise-induced | 1 |
| GCK | Hyperglycemia & macroglossia/umbilical hernia/large birth weight | 1 |
| GCK | Hypoglycaemia | 8 |
| GCK | Hypoglycaemia, hyperinsulinaemic | 2 |
| GCM2 | Hyperparathyroidism | 4 |
| GCM2 | Hyperparathyroidism, association with | 1 |
| GCM2 | Hypoparathyroidism | 15 |
| GNA11 | Hypercalcaemia, hypocalciuric | 1 |
| GNA11 | Hypercalcaemia, hypocalciuric type 2 | 2 |
| GNA11 | Hypocalcaemia | 2 |
| GNA11 | Hypocalcaemia type 2, autosomal dominant | 1 |
| HBA2 | Anaemia | 2 |
| HBA2 | Anaemia with hypochromia & microcytosis | 2 |
| HBA2 | Anaemia, hypochromic microcytic | 2 |
| HBA2 | Erythrocytosis | 2 |
| HBA2 | Haemolytic anaemia | 3 |
| HBA2 | Polycythaemia | 1 |
| HBB | Anaemia | 3 |
| HBB | Anaemia, hypochromic microcytic | 2 |
| HBB | Erythrocytosis | 21 |
| HBB | Haemolytic anaemia | 20 |
| HBB | Polycythaemia | 5 |
| IGF1R | Growth retardation | 9 |
| IGF1R | Growth retardation & microcephaly | 4 |
| IGF1R | Growth retardation, intrauterine & postnatal | 3 |
| IGF1R | Growth retardation, microcephaly & Silver-Russell syndrome features | 1 |
| IGF1R | Overgrowth | 2 |
| IGF1R | Overgrowth & variable intellectual disability | 2 |
| IGF1R | Short stature | 17 |
| IGF1R | Short stature & intrauterine growth retardation | 1 |
| IGF1R | Short stature, association with | 1 |
| IGF1R | Short stature, intellectual disability, Chiari malformation, microcephaly, hypotonia & lack of speech | 1 |
| IGF1R | Short staure & intrauterine growth retardation | 1 |
| IGF1R | SHORT syndrome | 1 |
| IL1RL1 | Higher sST2 levels, association with | 5 |

Supplementary Table 10 List of potential bidirectional effect selected tagets from HGMD

| Gene | Phenotype | Number of variants |
|---|---|---|
| IL1RL1 | Lower sST2 levels, association with | 1 |
| IL4R | Asthma, atopic, association with | 1 |
| IL4R | Asthma, increased risk, association with | 1 |
| IL4R | Atopic dermatitis, association with | 1 |
| IL4R | Atopy, association with | 2 |
| IL4R | Atopy, reduced risk, association with | 1 |
| INSIG2 | Obesity, association with | 1 |
| INSIG2 | Reduced BMI, association with | 1 |
| KCNJ2 | Long QT syndrome | 6 |
| KCNJ2 | Short QT syndrome | 2 |
| KCNJ2 | Short QT syndrome 3 | 1 |
| KCNJ2 | Short QT3 syndrome & autism-epilepsy phenotype | 1 |
| KCNQ1 | Long QT syndrome | 507 |
| KCNQ1 | Long QT syndrome & atrial fibrillation | 1 |
| KCNQ1 | Long QT syndrome with Takotsubo cardiomyopathy | 1 |
| KCNQ1 | Long QT syndrome, modifier of | 3 |
| KCNQ1 | Short QT syndrome | 2 |
| KITLG | Progressive hyper- and hypopigmentation | 4 |
| KITLG | Progressive hyperpigmentation | 1 |
| LCAT | Hypercholesterolaemia | 2 |
| LCAT | Increased HDL cholesterol levels | 1 |
| LCAT | Reduced high density lipoprotein-cholesterol | 5 |
| LCAT | Reduced plasma HDL cholesterol | 7 |
| LCAT | Reduced plasma HDL cholesterol, association with | 2 |
| LDLR | Coronary artery disease, protection against, association with | 1 |
| LDLR | Hypercholesterolaemia | 1990 |
| LDLR | Increased plasma LDL cholesterol | 1 |
| LDLR | Increased plasma LDL cholesterol, association with | 1 |
| LDLR | Myocardial infarction | 29 |
| LDLR | Reduced LDL cholesterol levels | 1 |
| LDLR | Stroke, increased risk, association with | 1 |
| LEP | High plasma leptin levels | 2 |
| LEP | Leptin deficiency | 8 |
| LEP | Low serum leptin levels | 2 |
| LIPG | HDL cholesterol levels, association with | 2 |
| LIPG | High HDL cholesterol | 1 |
| LIPG | Higher plasma HDL cholesterol | 13 |
| LIPG | Higher plasma HDL cholesterol, association with | 2 |
| LIPG | Higher plasma HDL cholesterol, in African Americans, association with | 1 |
| LIPG | Hypercholesterolaemia | 1 |
| LIPG | Low HDL cholesterol | 1 |
| LPL | Hypertriglyceridaemia | 64 |
| LPL | Increased HDL cholesterol, association with | 1 |
| LPL | Low HDL cholesterol | 1 |
| LPL | Lower plasma triglyceride level | 1 |
| LPL | Lower plasma triglyceride level, association with | 1 |
| LPL | Lower triglyceride level, association with | 2 |
| LPL | Reduced HDL level in atherosclerosis, association | 1 |
| LRP5 | Endosteal hyperostosis | 2 |
| LRP5 | High bone mass trait | 5 |
| LRP5 | Higher femoral neck bone mineral density, association with | 1 |
| LRP5 | Lower femoral neck bone mineral density, association with | 1 |
| LRP5 | Osteopetrosis | 5 |
| LRP5 | Osteoporosis & hyperlipidaemia | 1 |
| LRP5 | Osteoporosis-pseudoglioma syndrome | 64 |
| LRP5 | Osteoporosis, association with | 2 |
| LRP5 | Osteoporosis, primary | 4 |
| MC2R | ACTH hypersensitivity syndrome | 2 |
| MC2R | Adrenal hypoplasia, salt-losing | 1 |
| MC2R | Adrenal insufficiency, primary | 3 |
| MC2R | Cushing syndrome | 1 |
| NPC1 | High HDL cholesterol | 1 |
| NPC1 | Low HDL cholesterol | 1 |
| NPC1L1 | High dietary cholesterol absorption | 4 |
| NPC1L1 | Hypercholesterolaemia, association with | 1 |
| NPC1L1 | Increased serum cholesterol, association with | 1 |
| NPC1L1 | Low dietary cholesterol absorption | 20 |
| NPC1L1 | Reduced LDL cholesterol levels, association with | 11 |
| NPPC | Overgrowth and bone anomalies | 3 |
| NPPC | Short stature | 1 |
| NPPC | Skeletal overgrowth | 2 |
| NPR2 | Acromesomelic dysplasia, Maroteaux type | 37 |
| NPR2 | Short stature | 20 |
| NPR2 | Tall stature | 2 |
| NPR2 | Tall stature, scoliosis & macrodactyly of great toes | 2 |
| NR0B2 | Higher birth weight, association with | 1 |
| NR0B2 | Lower birth weight | 1 |
| NR3C1 | Glucocorticoid receptor deficiency | 13 |
| NR3C1 | Glucocorticoid resistance | 9 |
| NR3C1 | Increased glucocorticoid sensitivity | 2 |

Supplementary Table 10 List of potential bidirectional effect selected tagets from HGMD

| Gene | Phenotype | Number of variants |
|---|---|---|
| NTRK1 | Pain insensitivity, congenital | 54 |
| NTRK1 | Pain insensitivity, congenital with anhidrosis | 18 |
| NTRK1 | Sensory and autonomic neuropathy | 3 |
| NTRK1 | Sensory and autonomic neuropathy type IV | 11 |
| NTRK1 | Sensory and autonomic neuropathy type V | 1 |
| PCSK9 | Altered cholesterol levels | 1 |
| PCSK9 | Atherosclerosis, severity, association with | 1 |
| PCSK9 | Coronary heart disease, reduced risk, association with | 1 |
| PCSK9 | High LDL cholesterol | 8 |
| PCSK9 | High LDL cholesterol, association with | 6 |
| PCSK9 | Hypercholesterolaemia | 28 |
| PCSK9 | Hypercholesterolaemia, autosomal dominant | 13 |
| PCSK9 | Hypercholesterolaemia, modifier of | 1 |
| PCSK9 | Hyperlipidaemia | 1 |
| PCSK9 | Hypocholesterolaemia | 11 |
| PCSK9 | Low LDL cholesterol | 9 |
| PCSK9 | Low LDL cholesterol, association with | 7 |
| PPARG | Diabetes, type 2, increased risk | 8 |
| PPARG | Insulin resistance | 6 |
| PPARG | Insulin resistance, diabetes and hypertension | 2 |
| PPARG | Insulin sensitivity, association with | 1 |
| PRNP | Creutzfeldt-Jakob syndrome, increased risk | 1 |
| PRNP | Creutzfeldt-Jakob syndrome, protection, association with | 1 |
| PRNP | Prion disease, resistance to, association with | 1 |
| PTGS2 | Colorectal cancer, risk, association with | 1 |
| PTGS2 | Colorectal neoplasia, reduced risk, association | 1 |
| SCARB1 | Coronary heart disease, increased risk, association with | 1 |
| SCARB1 | HDL cholesterol levels, in women, association with | 1 |
| SCARB1 | Increased HDL cholesterol | 3 |
| SCARB1 | Increased HDL cholesterol and Lp(a) | 5 |
| SCARB1 | Increased HDL cholesterol level, association with | 1 |
| SCARB1 | Low HDL cholesterol | 1 |
| SCN4A | Hyperkalaemic periodic paralysis | 10 |
| SCN4A | Hypokalaemic periodic paralysis | 9 |
| SCN9A | Channelopathy-associated insensitivity to pain | 3 |
| SCN9A | Chronic non-paroxysmal neuropathic pain | 1 |
| SCN9A | Congenital indifference to pain | 27 |
| SCN9A | Congenital indifference to pain, partial | 2 |
| SCN9A | Erythermalgia, primary | 12 |
| SCN9A | Erythromelalgia | 17 |
| SCN9A | Erythromelalgia, late-onset | 1 |
| SCN9A | Neuropathy, hereditary sensory, type IID | 1 |
| SCN9A | Pain, dysautonomia & acromesomelia | 1 |
| SCN9A | Paroxysmal extreme pain disorder | 11 |
| SCN9A | Paroxysmal extreme pain disorder / Erythermalgia, primary | 1 |
| SCN9A | Peripheral neuropathy | 1 |
| SHOX | Dyschondrosteosis | 4 |
| SHOX | Langer mesomelic dysplasia | 13 |
| SHOX | Langer mesomelic dysplasia, milder phenotype | 1 |
| SHOX | Leri-Weill dyschondrosteosis | 140 |
| SHOX | Leri-Weill dyschondrosteosis with short stature | 2 |
| SHOX | Short stature | 121 |
| SHOX | Short stature, mental retardation & facial dysmorphisms | 1 |
| SHOX | Tall stature | 2 |
| SLC14A2 | Higher blood pressure in males, association with | 1 |
| SLC14A2 | Lower blood pressure in males, association with | 1 |
| SLC22A12 | Gout, association with | 1 |
| SLC22A12 | Gout, increased risk, association with | 1 |
| SLC22A12 | Gout, primary | 4 |
| SLC22A12 | Hyperuricaemia, association with | 1 |
| SLC22A12 | Hypouricaemia, renal | 33 |
| SLC22A12 | Hypouricaemia, renal, type 1 | 3 |
| SLC22A12 | Increased fractional excretion of uric acid, association | 1 |
| SLC22A12 | Reduced fractional excretion of uric acid, association | 2 |
| SLC26A9 | Increased current and chloride ion transport | 1 |
| SLC26A9 | Reduced current and chloride ion transport | 3 |
| SOST | Bone-mineral density, association with | 1 |
| SOST | Craniodiaphyseal dysplasia, autosomal dominant | 2 |
| SOST | High bone mass trait | 1 |
| SOST | Osteoporosis, association with | 1 |
| TFPI | Increased plasma TFPI | 2 |
| TFPI | Lower plasma TFPI | 1 |
| TGFB3 | Low muscle mass, hypotonia, growth retardation, arthrogryposis & bifid uvula | 1 |
| TGFB3 | Overgrowth & Loeys-Dietz syndrome | 1 |
| TGFB3 | Tall stature, archnodactyly, hyperextensible joints, hypertelorism, and bfid uvula | 1 |
| VDR | Higher bone mineral density, association with | 1 |
| VDR | Lower lumbar bone mineral density, association with | 1 |
| VDR | Rickets, vitamin D dependent, type II | 2 |
| VDR | Rickets, vitamin D dependent, type IIA | 2 |
| VDR | Rickets, vitamin D resistant | 49 |

Supplementary Table 10 List of potential bidirectional effect selected tagets from HGMD

| Gene | Phenotype | Number of variants |
|---|---|---|
| VDR | Rickets, vitamin D resistant, with alopecia | 2 |
| VDR | Rickets, vitamin D resistant, without alopecia | 5 |
| VWF | Altered VWF antigen/FVIII coagulant activity | 3 |
| VWF | Increased protein cleavage by ADAMTS13 | 1 |
| VWF | Increased VWF antigen levels, association with | 1 |
| VWF | Proteolysis by ADAMTS13, susceptibility to | 1 |
| VWF | Reduced protein cleavage by ADAMTS13 | 2 |
| VWF | Reduced vWF plasma protein levels | 1 |
| ABCB1 | Decreased drug resistance | 1 |
| ABCB1 | Increased drug resistance | 4 |
| ABCC1 | Increased multidrug response | 1 |
| ABCC1 | Reduced multidrug resistance | 3 |
| ADH1C | Alcoholism, increased risk, association with | 1 |
| ADH1C | Reduced alcohol metabolism, association with | 1 |
| AGT | Hypertension | 7 |
| AGT | Hypertension, association with | 6 |
| AGT | Increased angiotensinogen levels, association with | 1 |
| AGT | Reduced angiotensinogen levels | 1 |
| AGTR2 | Congenital anomalies of the kidney and urinary tract | 1 |
| AGTR2 | Increased glomerular filtration rate, in males | 1 |
| AGTR2 | Congenital anomalies of the kidney and urinary tract | 1 |
| AGTR2 | Increased glomerular filtration rate, in males | 1 |
| DIO1 | Higher plasma rT3 | 1 |
| DIO1 | Lower plasma rT3/T4 ratio | 1 |
| F2R | Coronary heart disease, association with | 1 |
| F2R | Low platelet receptor density, association with | 1 |
| F2R | Venous thromboembolism, protection (male), association | 1 |
| KISS1R | Central precocious puberty | 2 |
| KISS1R | Central precocious puberty, association with | 1 |
| KISS1R | Hypogonadotropic hypogonadism | 14 |
| KISS1R | Hypogonadotropic hypogonadism, idiopathic | 5 |
| KISS1R | Hypogonadotropic hypogonadism, isolated | 9 |
| KLF1 | Borderline HbA2 level | 8 |
| KLF1 | Dyserythropoietic anaemia | 1 |
| KLF1 | Haemolytic anaemia, nonspherocytic | 5 |
| KLF1 | Hereditary persistence of foetal haemoglobin | 4 |
| KLF1 | Increased Hb F levels | 13 |
| KLF1 | Increased Hb F levels and elevated HbA2 level | 3 |
| KLF1 | Increased HbA2 levels | 1 |
| KLF1 | Neonatal anaemia, severe | 1 |
| KLF1 | Thalassaemia beta | 1 |
| KLF1 | Thalassaemia beta, modifier of | 1 |
| KLK3 | Altered PSA level | 3 |
| KLK3 | Low PSA concentration | 3 |
| KLK3 | Prostate cancer, increased risk, association with | 2 |
| NGF | Loss of pain perception | 3 |
| NGF | Sensory and autonomic neuropathy | 2 |
| NQO2 | Breast cancer | 1 |
| NQO2 | Breast cancer, decreased risk, association with | 1 |
| PPARA | Elevated plasma lipid concentration, association in diabetes | 1 |
| PPARA | Lower total cholesterol, association with | 1 |
| REN | Prorenin elevation | 1 |
| REN | Reduced plasma renin activity | 1 |
| REN | Prorenin elevation | 1 |
| REN | Reduced plasma renin activity | 1 |
| SHMT1 | Increased folate levels, association with | 1 |
| SHMT1 | Neural tube defect | 1 |
| SLC11A2 | Anaemia, hypochromic microcytic | 5 |
| SLC11A2 | Haemochromatosis | 1 |
| SLC11A2 | Microcytic anaemia | 1 |
| SLC11A2 | Microcytic anaemia & iron overload | 1 |
| TLR4 | Acute pyelonephritis, in children, association with | 1 |
| TLR4 | Acute pyelonephritis, reduced risk in adults, association with | 1 |
| TLR4 | Acute pyelonephritis, in children, association with | 1 |
| TLR4 | Acute pyelonephritis, reduced risk in adults, association with | 1 |

#### Supplementary Table 11 List of bidirectional target - indication pairs

| symbol | ensembl_id | MSH (PharmaProjects) | lApprovedUS.EU | Phase.Latest | gwas_gene | best_mesh_gwas | best_trait_gwas | omim_gene | best_mesh_bidir | best_trait_bidir |
|---|---|---|---|---|---|---|---|---|---|---|
| AGT | ENSG00000135744 | Hypertension | TRUE | Approved | TRUE | 0.00 | cerebrospinal fluid biomarker measurement | TRUE | 1.00 | Hypertension |
| ANGPTL3 | ENSG00000132855 | Hypercholesterolemia | FALSE | Phase II Clinical Tria | TRUE | 0.90 | low density lipoprotein cholesterol measurement | TRUE | 1.00 | Hypercholesterolemia |
| APOA1 | ENSG00000118137 | Hypercholesterolemia | FALSE | Preclinical | TRUE | 0.50 | high density lipoprotein cholesterol measurement | TRUE | 1.00 | Hypercholesterolemia |
| APOB | ENSG00000084674 | Hypercholesterolemia | FALSE | Phase I Clinical Trial | TRUE | 0.90 | low density lipoprotein cholesterol measurement | TRUE | 1.00 | Hypercholesterolemia |
| BDKRB2 | ENSG00000168398 | Hypertension | FALSE | Preclinical | FALSE | 0.00 | NA | TRUE | 1.00 | Hypertension |
| CASR | ENSG00000036828 | Hypercalcemia | TRUE | Approved | TRUE | 0.00 | calcium measurement | TRUE | 1.00 | Hypercalcemia |
| CASR | ENSG00000036828 | Hypercalcemia | TRUE | Approved | TRUE | 0.00 | calcium measurement | TRUE | 1.00 | Hyperparathyroidism |
| CASR | ENSG00000036828 | Hyperparathyroidism, Secondary | TRUE | Approved | TRUE | 0.00 | calcium measurement | TRUE | 0.91 | Hypercalcemia |
| CASR | ENSG00000036828 | Hyperparathyroidism, Secondary | TRUE | Approved | TRUE | 0.00 | calcium measurement | TRUE | 0.91 | Hyperparathyroidism |
| CASR | ENSG00000036828 | Hyperparathyroidism, Primary | TRUE | Approved | TRUE | 0.00 | calcium measurement | TRUE | 0.90 | Hypercalcemia |
| CASR | ENSG00000036828 | Hyperparathyroidism, Primary | TRUE | Approved | TRUE | 0.00 | calcium measurement | TRUE | 0.90 | Hyperparathyroidism |
| CCL5 | ENSG00000271503 | HIV Infections | FALSE | Phase II Clinical Tria | TRUE | 0.00 | reticulocyte count | TRUE | 1.00 | HIV Infections |
| CCR5 | ENSG00000160791 | HIV Infections | TRUE | Approved | TRUE | 1.00 | HIV-1 infection, HIV viral set point measurement | TRUE | 1.00 | HIV Infections |
| CETP | ENSG00000087237 | Hypercholesterolemia | FALSE | Phase III Clinical Tria | TRUE | 0.90 | low density lipoprotein cholesterol measurement | TRUE | 1.00 | Hypercholesterolemia |
| CYP19A1 | ENSG00000137869 | Breast Neoplasms | TRUE | Approved | TRUE | 0.52 | endometrial carcinoma | TRUE | 1.00 | Breast Neoplasms |
| EGLN1 | ENSG00000135766 | Anemia | FALSE | Preclinical | TRUE | 0.75 | hemoglobin measurement | TRUE | 1.00 | Anemia |
| F2 | ENSG00000180210 | Hemophilia A | TRUE | Approved | TRUE | 0.18 | acne | TRUE | 1.00 | Hemophilia A |
| F5 | ENSG00000198734 | Hemophilia A | FALSE | Preclinical | TRUE | 0.17 | venous thromboembolism | TRUE | 1.00 | Hemophilia A |
| F7 | ENSG00000057593 | Hemophilia A | TRUE | Approved | TRUE | 0.00 | coagulation factor measurement | TRUE | 1.00 | Hemophilia A |
| F8 | ENSG00000185010 | Hemophilia A | TRUE | Approved | TRUE | 0.18 | type I diabetes mellitus | TRUE | 1.00 | Hemophilia A |
| F9 | ENSG00000101981 | Hemophilia A | TRUE | Approved | FALSE | 0.00 | NA | TRUE | 1.00 | Hemophilia A |
| FGFR3 | ENSG00000068078 | Dwarfism | FALSE | Preclinical | TRUE | 0.18 | bladder carcinoma | TRUE | 0.84 | Achondroplasia |
| HBB | ENSG00000244734 | Anemia | FALSE | Preclinical | TRUE | 0.75 | hemoglobin A2 measurement | TRUE | 1.00 | Anemia |
| IGF1R | ENSG00000140443 | Dwarfism | TRUE | Approved | TRUE | 0.18 | endometriosis | TRUE | 0.84 | Achondroplasia |
| IL4R | ENSG00000077238 | Asthma | FALSE | Phase III Clinical Tria | TRUE | 0.00 | basophil count, eosinophil count | TRUE | 1.00 | Asthma |
| KISS1R | ENSG00000116014 | Hypogonadism | FALSE | unknown | TRUE | 0.00 | reticulocyte count | TRUE | 1.00 | Hypogonadism |
| KLK3 | ENSG00000142515 | Prostatic Neoplasms | FALSE | Phase III Clinical Tria | TRUE | 1.00 | prostate carcinoma | TRUE | 0.90 | Prostate-Specific Antigen |
| LDLR | ENSG00000130164 | Hypercholesterolemia | FALSE | Preclinical | TRUE | 0.90 | low density lipoprotein cholesterol measurement | TRUE | 1.00 | Hypercholesterolemia |
| LEP | ENSG00000174697 | Obesity | FALSE | Preclinical | FALSE | 0.00 | NA | TRUE | 1.00 | Obesity |
| LPL | ENSG00000175445 | Hypercholesterolemia | TRUE | Approved | TRUE | 0.50 | high density lipoprotein cholesterol measurement | TRUE | 1.00 | Hypercholesterolemia |
| LRP5 | ENSG00000162337 | Osteoporosis | FALSE | Preclinical | TRUE | 0.90 | bone density | TRUE | 1.00 | Osteoporosis |
| NPC1L1 | ENSG00000015520 | Hypercholesterolemia | TRUE | Approved | TRUE | 0.90 | low density lipoprotein cholesterol measurement | TRUE | 1.00 | Hypercholesterolemia |
| NR3C1 | ENSG00000113580 | Inflammation | TRUE | Approved | FALSE | 0.00 | NA | TRUE | 1.00 | Inflammation |
| PCSK9 | ENSG00000169174 | Hypercholesterolemia | TRUE | Approved | TRUE | 0.90 | low density lipoprotein cholesterol measurement | TRUE | 1.00 | Hypercholesterolemia |
| PPARA | ENSG00000186951 | Hypercholesterolemia | TRUE | Approved | FALSE | 0.00 | NA | TRUE | 1.00 | Hypercholesterolemia |
| PPARG | ENSG00000132170 | Diabetes Complications | FALSE | Preclinical | TRUE | 0.77 | type II diabetes mellitus | TRUE | 0.83 | Diabetes Mellitus |
| PRNP | ENSG00000171867 | Creutzfeldt-Jakob Syndrome | FALSE | Phase I Clinical Trial | TRUE | 1.00 | Creutzfeldt Jacob Disease | TRUE | 1.00 | Creutzfeldt-Jakob Syndrome |
| PTGS2 | ENSG00000073756 | Colorectal Neoplasms | FALSE | Phase II Clinical Tria | FALSE | 0.00 | NA | TRUE | 1.00 | Colorectal Neoplasms |
| SCARB1 | ENSG00000073060 | Hypercholesterolemia | FALSE | Preclinical | TRUE | 0.50 | high density lipoprotein cholesterol measurement | TRUE | 1.00 | Hypercholesterolemia |
| SCN9A | ENSG00000169432 | Pain | FALSE | Preclinical | FALSE | 0.00 | NA | TRUE | 1.00 | Pain |
| SLC22A12 | ENSG00000197891 | Hyperuricemia | TRUE | Approved | TRUE | 0.90 | gout | TRUE | 0.90 | Gout |
| SOST | ENSG00000167941 | Osteoporosis | FALSE | Pre-registration | TRUE | 0.90 | bone density | TRUE | 1.00 | Osteoporosis |
| TFPI | ENSG00000003436 | Hemophilia A | FALSE | Phase II Clinical Tria | TRUE | 0.00 | BMI-adjusted waist-hip ratio | TRUE | 1.00 | Hemophilia A |
| VDR | ENSG00000111424 | Osteoporosis | TRUE | Approved | FALSE | 0.00 | NA | TRUE | 1.00 | Osteoporosis |
| VWF | ENSG00000110799 | Hemophilia A | TRUE | Approved | TRUE | 0.00 | coagulation factor measurement | TRUE | 1.00 | Hemophilia A |

List of target - indication pairs with clinical trial information from PharmaProjects for which a Bidirectional target with MESH > 0.83 was found.
